# Rapid gene exchange explains differences in bacterial pangenome structure

**DOI:** 10.64898/2026.02.04.703695

**Authors:** Samuel T. Horsfield, Anlei Peng, Matthew J. Russell, Johanna von Wachsmann, Jacqueline Toussaint, Joshua C. D’Aeth, Chuhan Qin, Henri Pesonen, Gerry Tonkin-Hill, Jukka Corander, Nicholas J. Croucher, John A. Lees

**Author notes:** Contributed equally (Nicholas J. Croucher, John A. Lees).

## Abstract

The size and diversity of bacterial gene repertoires, known as pangenomes, vary widely across species. The evolutionary forces driving the maintenance of pangenomes is an open topic of debate, with contradictory theories suggesting that pangenomes exist as a result of neutral evolution, with all genes gained and lost at random, or that all genes provide a fitness benefit to the host and are maintained by positive selection. Modelling of pangenome dynamics has provided insight into how gene exchange explains observed gene frequency distributions, and stands as the only means of jointly inferring contributions of individual gene selection effects and mobility on the maintenance of pangenomes. However, previous modelling studies have not included both gene-level selection and mobility, and do not consider broadly sampled genome datasets for many species. To differentiate neutral and selective forces maintaining pangenomes, we developed a mechanistic model of gene-level evolution, Pansim, and a scalable model fitting framework, PopPUNK-mod. Together, these tools leverage rapid genome distance calculation to fit models of pangenome dynamics to datasets containing hundreds of thousands of genomes. We used this framework to compare the pangenome dynamics of over 400 different bacterial species, using over 600,000 genomes. We find that diversity in pangenome characteristics between species is driven predominantly by variation in the number of rapidly exchanged genes, while the rate of exchange of remaining genes is conserved. We find that bacterial phylogeny, rather than ecology, correlates with pangenome dynamics. We express that pan-species gene-level analyses are now needed to understand selection across accessory genes. Our work highlights the importance of gene exchange rate differences in governing differences in pangenome characteristics between species.

## Introduction

Many bacterial species exhibit a huge diversity in their respective gene repertoires contained within a pangenome: a collection of genomic variation representing the full diversity of a population (Tettelin et al. 2005; Vernikos et al. 2015). Conventionally, genes are grouped into two categories based on their frequency within a population: core genes, which are typically found in all members of a species and define the fundamental biology of a species (Young et al. 2006; Koonin and Wolf 2008); and accessory genes, whose presence is variable across the population, and which contribute to genotypic and phenotypic diversity within a species, such as differences in drug resistance (Jaillard et al. 2017; McNally et al. 2019), vaccine susceptibility (Lo et al. 2019), host range (Dearlove et al. 2015; Weinert et al. 2015) and virulence (Alikhan et al. 2018; Hennart et al. 2020).

Previous inter-species pangenome comparisons have highlighted large variability of the size and structure of the accessory genome (Bobay and Ochman 2018; Andreani et al. 2017; Baumdicker and Pfaffelhuber 2014; Dewar et al. 2024), indicating that the relative effects of the forces driving the formation and maintenance of accessory genomes are not consistent across bacteria. An open question exists as to whether the forces shaping accessory genomes are: primarily adaptive and driven by niche-associated fitness benefits or costs (McInerney et al. 2017); associated with balancing or negative frequency-dependent selection, in which strains express phenotypes that are maximally advantageous when only present in a fraction of the population (Corander et al. 2017; Harrow et al. 2021); neutral, whereby genes are gained and lost without having a substantial phenotypic impact on the cell (Shapiro 2017; Rocha 2018); or selfish, having a negative impact on host fitness, with gene frequency tied to individual rates of transfer (Douglas and Shapiro 2021; Vos and Eyre-Walker 2017). Currently, modelling of pangenome dynamics stands as the only means to capture the individual fitness contributions of all genes in a pangenome to observed gene frequencies.

Recent work has focused on modelling gene-level dynamics to understand how genic selection and mobility gives rise to and maintains pangenomes. Across various species, genes within the same pangenome exhibit large differences in turnover rate, suggestive of non-uniform forces of selection or mobility (Sela et al. 2021; Gamblin et al. 2025), while some, but not all, rare accessory genes, show evidence of maintenance by positive selection (Douglas and Shapiro 2024). These between-gene differences in dynamics in the accessory genome strongly supports a more complex situation than accessory genes being wholly neutral or adaptive, with accessory gene frequency being an interaction between selection for genes differentiating lineages, and horizontal gene transfer (HGT) (Baumdicker and Kupczok 2023; N’Guessan et al. 2021).

We require model architectures and datasets that are sufficiently complex and representative to capture the interactions between genic selection and mobility that govern gene frequency. However, estimates of gene frequency are sensitive to errors in gene prediction, which can greatly impact the accuracy of parameter inference when modelling pangenome dynamics (Tonkin-Hill, Gladstone, et al. 2023; Tonkin-Hill, Corander, et al. 2023). Furthermore, models of pangenome dynamics benefit greatly from wide, deep population sampling in order to account for variation in gene selection and mobility in different lineages (Tonkin-Hill, Corander, et al. 2023). Vast databases of bacterial genomes now exist, containing tens to hundreds of thousands of genomes for individual species (Hunt et al. 2024; Schmidt et al. 2024; Richardson et al. 2023). However, current gene-level models scale to only tens or hundreds of genomes (Douglas and Shapiro 2024; Tonkin-Hill, Gladstone, et al. 2023; Sela et al. 2021; Gamblin et al. 2025). Therefore, we currently lack a means of fitting complex evolutionary models to large publicly-available species genome datasets (Hunt et al. 2024; Timme et al. 2019).

Previously, we developed a highly scalable genomic epidemiological analysis tool, PopPUNK (Lees et al. 2019), which represents pangenome-wide species diversity in terms of pairwise core and accessory distances between individual genomes. PopPUNK uses k-mer matching to calculate distances between tens of thousands of individual genomes. Therefore, PopPUNK provides a gene-agnostic method for distinguishing pangenome dynamics between species which is insensitive to errors in gene prediction, and can scale to the size of publicly-available genome datasets. We have also previously shown in individual species that the structure of these pairwise distances is representative of species population structure, and is shaped by the pangenome dynamics shaping their respective evolutionary histories (Croucher et al. 2014; Marttinen et al. 2015; Harrow et al. 2021). Therefore, pairwise distance information holds promise as a useful data source for modelling the molecular rates governing pangenome evolution.

In this work, we develop an approach for modelling pangenome dynamics using PopPUNK which addresses issues in gene frequency estimation and model scalability. We first qualitatively compare distributions of core and accessory pairwise distance between four bacterial species with markedly different ecologies, highlighting the differences shown in PopPUNK distance distributions that correlate with the unique biology of each species. We then describe a novel pangenome simulator, Pansim, which can model a multitude of neutral and non-neutral pangenome dynamics through molecular events that underlie pangenome diversification. We show that Pansim can generate a wide range of core and accessory distance distributions, which are sensitive to variations in pangenome dynamics parameters. Finally, we develop PopPUNK-mod, an Approximate Bayesian Computation (ABC) method for fitting Pansim to observed genome data. Applying PopPUNK-mod to over 400 different species with a variety of lifestyles, we are able to propose a new model for the observed inter-species differences in pangenome sizes, which is driven largely by differences in rapid gene turnover across the bacterial domain.

## Results

### The relationship between core and accessory genome diversification rates across species is characteristic of bacterial ecology and evolutionary history

Initially, we selected four focal species to illustrate the effect of ecology and evolutionary history on the PopPUNK core and accessory distance distributions (**Figure 1, Supplementary Table 1**). These distributions reveal key heterogeneities in population structure and overall diversity between species.

**Figure 1:**
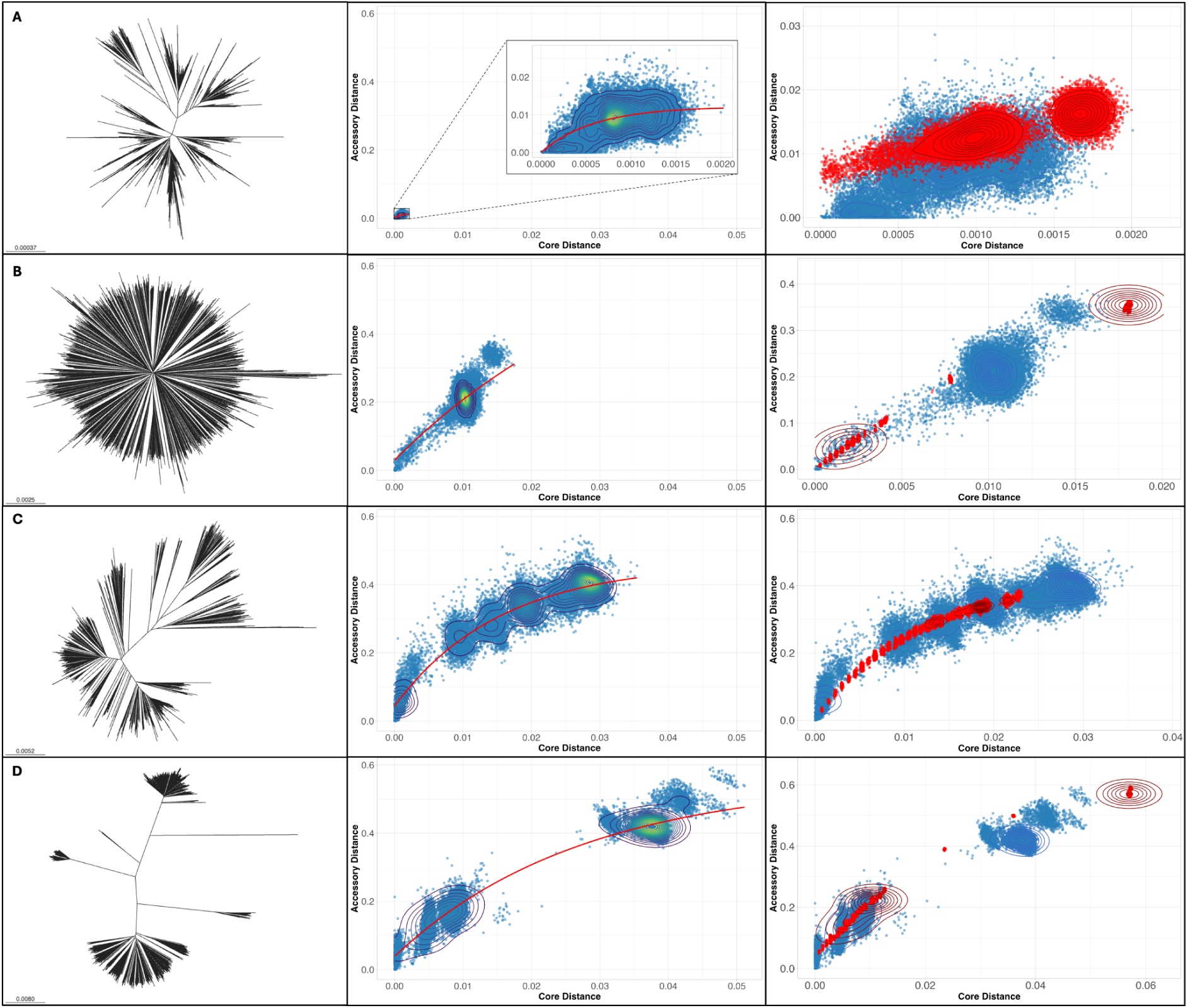
Comparison of population diversity between taxonomically distinct bacterial species; (**A**) *Mycobacterium tuberculosis*, (**B**) *Streptococcus pneumoniae*, (**C**) *Escherichia coli* and (**D**) *Listeria monocytogenes*. Left column - core genome phylogeny generated using PopPUNK and Microreact. Middle column - PopPUNK core versus accessory pairwise distance distributions. Red lines show non-linear negative exponential regression line (**Methods, Equation 1**). Right column - PopPUNK core versus accessory pairwise distance distributions from corresponding pathogen populations (blue) and simulated data generated by our evolutionary simulator, Pansim (red). Contours indicate point density.

*Mycobacterium tuberculosis* is an obligate human pathogen which is therefore reproductively-isolated and host-restricted with little to no evidence of recombination. These features result in a clonal population lacking a definitive accessory genome, and forming distinctive lineages defined by vertically-inherited core mutations (Gagneux 2018; Marin et al. 2025). The phylogenetic tree of *M. tuberculosis* shows this structured population clearly, while the PopPUNK core vs. accessory distance distributions highlight its low pangenome diversity (Figure 1A).

*S. pneumoniae* is an obligate human commensal and opportunistic invasive pathogen that, unlike *M. tuberculosis*, is naturally competent. Hence it is able to recombine with co-colonising bacteria, including other Streptococci, when inhabiting the human upper respiratory tract (D’Aeth et al. 2021; Johnston et al. 2013). Due to high rates of within-species recombination, *S. pneumoniae* has a non-heirarchical population structure, consisting of genetically equidistant strains (Marttinen et al. 2015; Harrow et al. 2021; Lees et al. 2019). The phylogenetic tree of *S. pneumoniae* has long terminal branches as a result of the clonal frame being overwritten, whilst the PopPUNK core vs. accessory distances show a single point of high density representing many equally-distant between-strain comparisons (Figure 1B).

*Escherichia coli* is a multi-host commensal found across a multitude of animal hosts, with certain strains associated with disease (Gordon 2013). *E. coli* has a highly structured population, which can be divided into multiple strains that themselves can be hierarchically clustered into phylogroups (Denamur et al. 2021; Tenaillon et al. 2010) (Figure 1C**, Supplementary** Figure 1). This structure is due to the persistence of relationships through common ancestry that have not been overwritten by more recent recombination, with *E. coli* having a lower homologous recombination rate compared to *S. pneumoniae (Torrance et al. 2024)*. This hierarchical population structure is also shown by PopPUNK distances calculated from comprehensive databases of *E. coli* isolated from human hosts and the environment (Horesh et al. 2021; Timme et al. 2019), with multiple distinct clusters of between-strain distances indicating a stable stepwise accruement of diversity from within- to between-strain comparisons.

*Listeria monocytogenes* is a free-living bacterium that also has a highly structured population separated into lineages which differ in the individual sizes of their pangenomes, habitat distribution and rates of homologous recombination (Lourenco et al. 2022; Liao et al. 2021; Orsi et al. 2011). The heterogeneity of recombination rates between lineages is highlighted in the phylogenetic tree, with lineages separated by long internal branches, and terminal branch lengths that vary in a strain-dependent manner (Figure 1D**, Supplementary** Figure 1). The PopPUNK distances further highlight this within-strain heterogeneity, with three tight clusters at low distance values, followed by a series of clusters found at very high distance values, indicating varying degrees of genetic overlap between strains.

Despite the notable differences in ecology and population structure between these four species, we observe similarities between the overall shapes of core vs. accessory pairwise distance distributions across a range of bacterial species (Figure 1, middle panels). Although core and accessory distances are positively correlated, these distributions appear to show a saturation effect in the accessory genome diversity as core genome distance increases. To further understand these differences, we expanded our analysis to 22 species with high quality population genome data available, spanning a greater breadth of the bacterial domain. To quantitatively characterise this relationship, we fitted negative exponential curves (Figure 1, middle panel, **Supplementary** Figures 2 **& 3**, see **Methods**) and compared fitted parameters between all 22 species.

We compared the parameters from these curves with annotations of lifestyle, which have been shown to correlate with population characteristics such as pangenome size and recombination rate (**Supplementary Table 1**) (Bobay and Ochman 2018; Dewar et al. 2024). We observed that the instantaneous gene turnover rate (y-intercept) was lower in single-host pathogens than in all other lifestyle categories, although not significantly different (**Supplementary** Figures 2 **& 3**). This observation suggests that non-host restricted and non-obligate pathogen species possess rapidly-exchanged genetic material which varies at a rate higher than the rate at which mutations occur in the core genome. Conversely, host-restricted obligate pathogens do not possess genetic material that is rapidly turned over. The remaining two parameters, the plateau and rate ratio, which define the saturation point of the accessory distance and the rate of asymptotic approach respectively, did not consistently vary between lifestyles.

To understand the effect of sampling bias on the regression fitted parameters, we additionally analysed a local sampling study of *S. pneumoniae* from the the Maela refugee camp (Chewapreecha et al. 2014). Confidence intervals of fitted parameters between the global and local *S. pneumoniae* sample do not overlap (**Supplementary** Figure 2), indicating that regression analysis is not robust to sampling bias, despite distributions appearing highly similar (**Supplementary** Figure 4). Therefore, over or undersampling of specific strains for other species, which is likely for disease-associated strains, will likely impact conclusions made from this analysis.

### Pansim is a novel pangenome simulator which captures complex pangenome dynamics

While PopPUNK distance distributions between different species were qualitatively similar, a simple regression model was not able to explain the relationship between ecological characteristics and population-level diversity, and was sensitive to sampling bias. To better capture the pangenome dynamics underlying the observed relationship between core and accessory diversity, we developed a flexible evolutionary simulator, Pansim, which can generate population structures of a diverse array of bacterial species (Figure 1, right panels). Several other bacterial population simulators exist which capture complex neutral population and pangenome dynamics, such as migration and recombination (Sipola et al. 2018; Brown et al. 2016), or selection in the form of selective sweeps of single alleles (Baumdicker et al. 2022; Kern and Schrider 2016; Cury et al. 2022), but do not comprehensively model neutral and selective processes.

In its basic form, Pansim is a multi-compartment Wright-Fisher model (Ewens 2004), with populations simulated forward in time with non-overlapping generations, with mutations allowed to accrue in the core genome as single nucleotide polymorphisms (SNPs) and accessory genome as gene gain/loss events (Figure 2A). Additionally, Pansim simulates neutral processes of homologous recombination (HR) in the core genome, and HGT in the accessory genome between members of the same generation (Figure 2B). As we observed that some species may accrue accessory changes at a greater rate than core mutations occur (**Supplementary** Figures 2 **& 3**), Pansim also models a two compartment accessory genome, each with their own rate of gene gain/loss. This model format has previously been shown to optimally fit observed gene-frequency distributions (Collins and Higgs 2012), and also reflects realistic molecular rate differences of mobile genetic elements compared to other accessory genes (N’Guessan et al. 2021).

**Figure 2:**
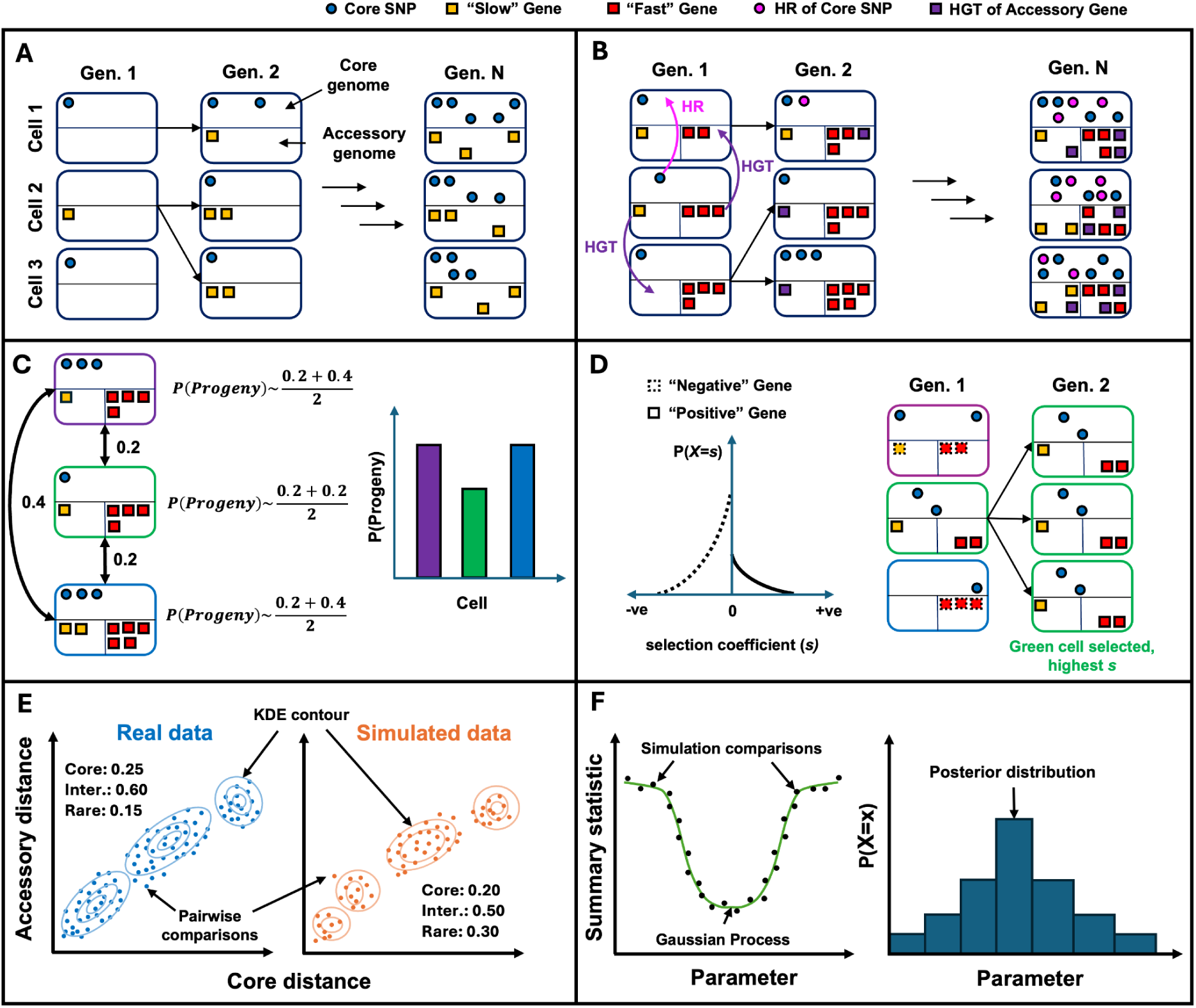
Overview of Pansim and model fitting with PopPUNK-mod. (**A**) Pansim uses a Wright-Fisher process, in which each individual cell accrues core genome SNPs, and gains and loses genes in the accessory genome. (**B**) Pansim enables sharing of alleles between members of the same generation, either through homologous recombination (HR, pink arrows) in the core genome, or horizontal gene transfer (HGT, purple) arrows in the accessory genome. (**C**) Pansim simulates competition between members of the same generation by downweighting fitness based on average pairwise distances across individuals. (**D**) Pansim models gene fitness effects as a double exponential distribution about 0, which are multiplied to generate an individual-level selection coefficient. (**E**) Summary statistic generation using Kernel Density Estimation (KDE) and Jensen-Shannon Distance (JSD) calculation between pairwise distances and pangenome statistics. (**F**) Summary statistic used in Approximate Bayesian Computation (ABC) to generate posterior distributions for each fitted parameter in PopPUNK-mod.

Pansim also models competition, under the assumption that more closely related individuals more strongly compete with each other for limited resources, following the competitive exclusion principle (Corander et al. 2017; Harrow et al. 2021; Hardin 1960) (Figure 2C), and gene-wise selection, where the selection landscape of genes is approximated by a double-exponential distribution, with the product of all gene-wise coefficients in an individual genome defines its fitness (Figure 2D).

To characterise the population dynamics underpinning observed bacterial populations, we also developed a method of fitting Pansim to observed data. Due to the complexity of Pansim, required for a biologically realistic simulation of pangenome dynamics, there is no analytical solution for computing the likelihood of a data set given a model parameterisation. Instead, we implemented a likelihood-free method of model fitting using Approximate Bayesian Computation (ABC) in BOLFI (Lintusaari et al. 2018; Gutmann and Corander 2015), which compares simulated data generated under a range of parameterisations with a target dataset using a summary statistic. Through rounds of iterative testing, we developed an informative summary statistic which describes the two-dimensional distribution of core and accessory distances, along with the proportions of core, intermediate and rare genes in a pangenome. The summary statistic is generated by creating a two-dimensional density representation of core vs. accessory distributions, using Kernel Density Estimation (KDE) for the target and simulated data, and calculating the Jensen-Shannon Distance (JSD) between them (Figure 2E). This summary statistic was able to distinguish observed bacterial species based on lifestyle, separating pathogens, commensals and free-living bacteria (**Supplementary** Figure 5). The distribution of Jensen-Shannon distances across each model parameterisation is then approximated using a Gaussian process by BOLFI-ABC, which is explored using a Markov Chain Monte Carlo (MCMC) algorithm to generate a posterior distribution for each fitted parameter (Figure 2F). This process of fitting Pansim to target distance distributions using ABC is provided by the PopPUNK-mod package (**Methods**). A sensitivity analysis of the effects of parameter values on pairwise distance distributions is detailed in **Supplementary Results**.

### Modelling pangenome dynamics across the bacterial tree of life shows rapid gene exchange rates are explained by phylogeny, not ecology

To characterise the differences in pangenome dynamics across the bacterial domain, we fitted our model to 271,939 genomes from 408 bacterial species from the AllTheBacteria dataset (Hunt et al. 2024), which contains a comprehensive representation of all publicly available sequenced bacterial isolate genomes. PopPUNK-mod was fitted to each species individually for three parameters: basal gene turnover rate, the minimum rate at which genes are gained and lost; core mutation rate, the rate at which single nucleotide polymorphisms occur in the core genome; and the proportion of fast genes, the proportion of total accessory genes that are exchanged instantaneously in each generation (for discussion of parameter choices, see **Supplementary Results;** parameter units detailed in **Supplementary Table 2**). We built a phylogenetic tree using marker genes from a single representative genome for each species, and overlaid parameter estimates for each species onto the tree (Figure 3). We calculated Pagel’s λ (Pagel 1999) for each median parameter estimate, which identifies whether a phylogenetic signal is present (λ > 0) or absent (λ = 0) for associated continuous traits.

**Figure 3:**
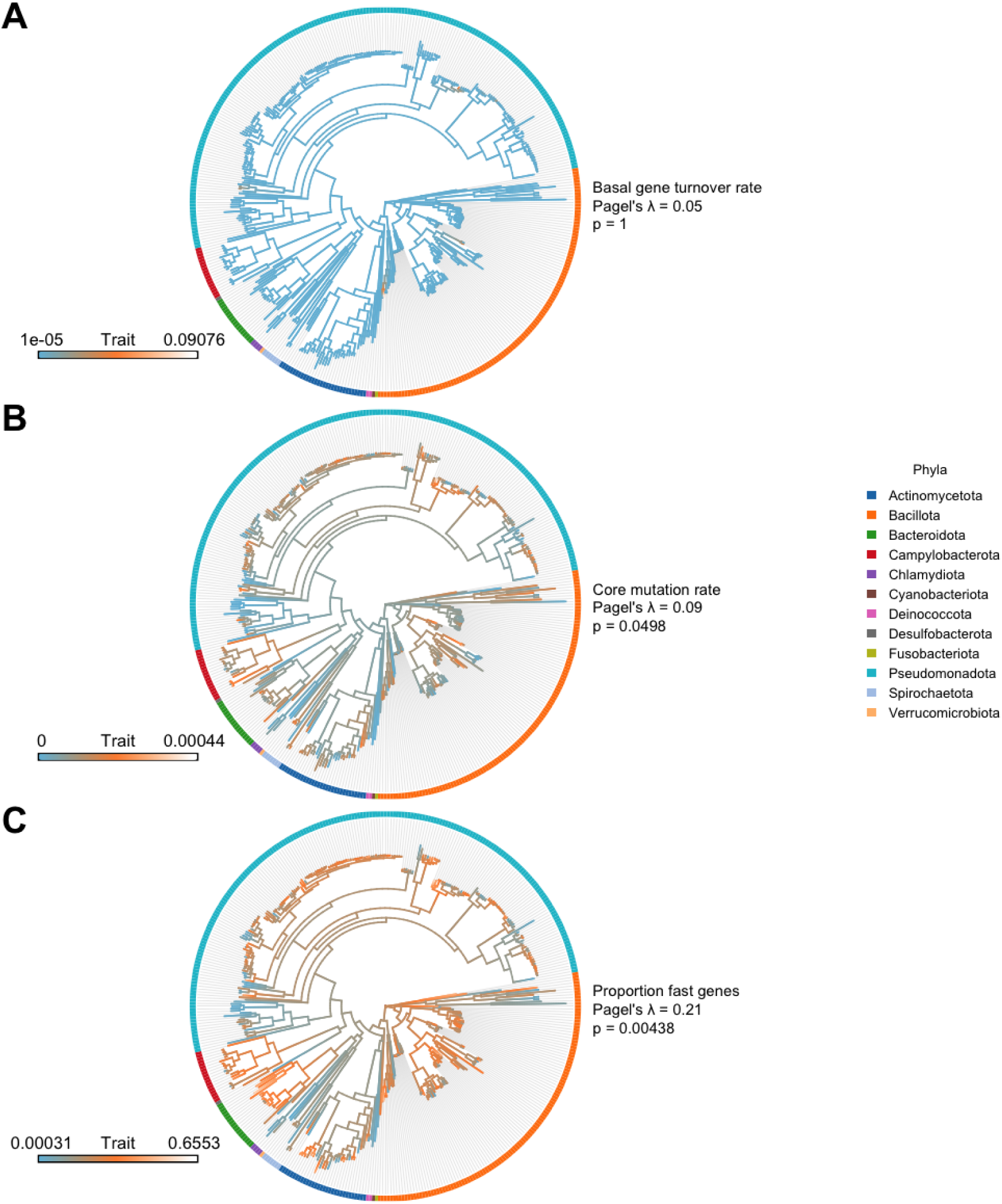
Pangenome dynamics comparison across 408 bacterial species in AllTheBacteria. Model median parameters used were (**A**) basal gene turnover rate, (**B**) core mutation rate and (**C**) proportion of fast genes. Parameter units detailed in **Supplementary Table 2**. Branches are coloured according to parameter value from blue (low) to orange/white (high), denoted by trait scale in bottom left of each plot. Pagel’s λ was used to calculate the degree of correlation between each parameter and the underlying phylogeny, and is displayed per parameter along with respective p-values calculated using phytools. Phyla designations were assigned from GTDB. Data for each species is available in **Supplementary File 1**.

Using Pagel’s λ, we identified a phylogenetic signal for the median core mutation rate and the proportion of fast genes, while no signal was found for basal gene turnover rate (Figure 3). This relationship between parameter estimates and phylogenetic signal was consistent for lower and upper credible intervals, with the exception of the lower credible interval of core mutation rate, for which no significant phylogenetic signal was detected (**Supplementary** Figures 6 **& 7**). Although basal gene turnover rate was more consistent between species than core mutation rate and the proportion of fast genes (**Supplementary** Figure 8), there was a significant relationship between all three parameters and phyla designations (**Supplementary** Figure 9). This result is in line with Pagel’s λ, with the exception of basal gene turnover rate, indicating that inter-phyla differences between parameters are larger than inter-species differences detected by Pagel’s λ. Differences in the proportion of fast genes between phyla aligned with known biological characteristics: *Chlamydiaota*, which we predicted to have comparably low proportions of fast genes and to which belong obligate intracellular *Chlamydia* species, are thought to undergo little exchange of genetic material (Marti et al. 2022), while *Bacteriodota* species, which we predicted to have comparably high predicted proportions of fast genes and to which belong commensal *Bacteroides* species, have broad pangenomes with large varieties of metabolism genes (Shin et al. 2024). Only core mutation rate was significantly correlated with both the basal gene turnover rate and the proportion of fast genes (**Supplementary** Figure 10), agreeing with previous observations that core and accessory diversity are positively correlated (Figure 1), and suggesting that genes turn over in the two accessory compartments independently of each other. Overall, there is evidence that core mutation rates and the proportion of fast genes in pangenomes evolve according to the phylogeny, where more closely related species have more similar dynamics.

The observation that more closely related species have more similar pangenome dynamics may be due to ecological similarity between species (Dewar et al. 2024). To determine the relationship between species lifestyle and PopPUNK-mod parameter estimates, we regressed each parameter against generalism score, a measure of a species’ ability to colonise multiple distinct habitats (Podlesny et al. 2025) using data from 85 species for which this score was available (**Supplementary** Figures 11 **& 12**), and accounting for population structure using the phylogeny as a random effect, as conducted previously (Dewar et al. 2024). We did not identify a significant relationship between generalism score and any of the fitted parameters from PopPUNK-mod, with the phylogeny accounting for 62% of observed variation in generalism score based on R^2^ values (**Supplementary Table 3**). Overall, our analysis indicates variability in accessory gene turnover primarily driven by rapidly exchanged genes, with variability correlating with phylogenetic relationships between species, while not being exclusively correlated with ecological characteristics.

### Fine-grained performance analysis of PopPUNK-mod

Due to the infeasibility of conducting a detailed investigation of PopPUNK-mod performance on 408 species, we first analysed performance on simulated data (**Supplementary Results**). We found PopPUNK-mod was able to accurately recapitulate basal gene turnover rate, core mutation rate and proportion of fast genes, with these parameters being robust to fixed parameter misspecification (**Supplementary** Figure 13), addressing the issue of sensitivity to gene prediction and clusters observed when modelling using gene frequency distributions (Tonkin-Hill, Gladstone, et al. 2023).

We then conducted a fine-grained performance analysis of PopPUNK-mod on a subset of 22 species for which hundreds to thousands of genomes are available from global sequencing projects (**Supplementary Table 1**). Results highlighted that species varied in their dynamics as described by each model parameter (Figure 4). Parameter ranges were largely consistent between species with the same lifestyles. As observed in Figure 3, basal gene turnover rate was consistent across all species, with the exception of *M. tuberculosis*, which exhibited low overall gene turnover, indicative of the reproductive isolation of the species and minimal accessory genome diversity (Gagneux 2018; Marin et al. 2025). Core mutation rate was lower in single-host pathogens, correlating with lowered growth rates and resulting accruement of small mutations (Bobay and Ochman 2018). *H. pylori* had the highest estimated core mutation rate, agreeing with previous observations, which is postulated to be due to high rates of homologous recombination (Duchêne et al. 2016; Kennemann et al. 2011).

**Figure 4:**
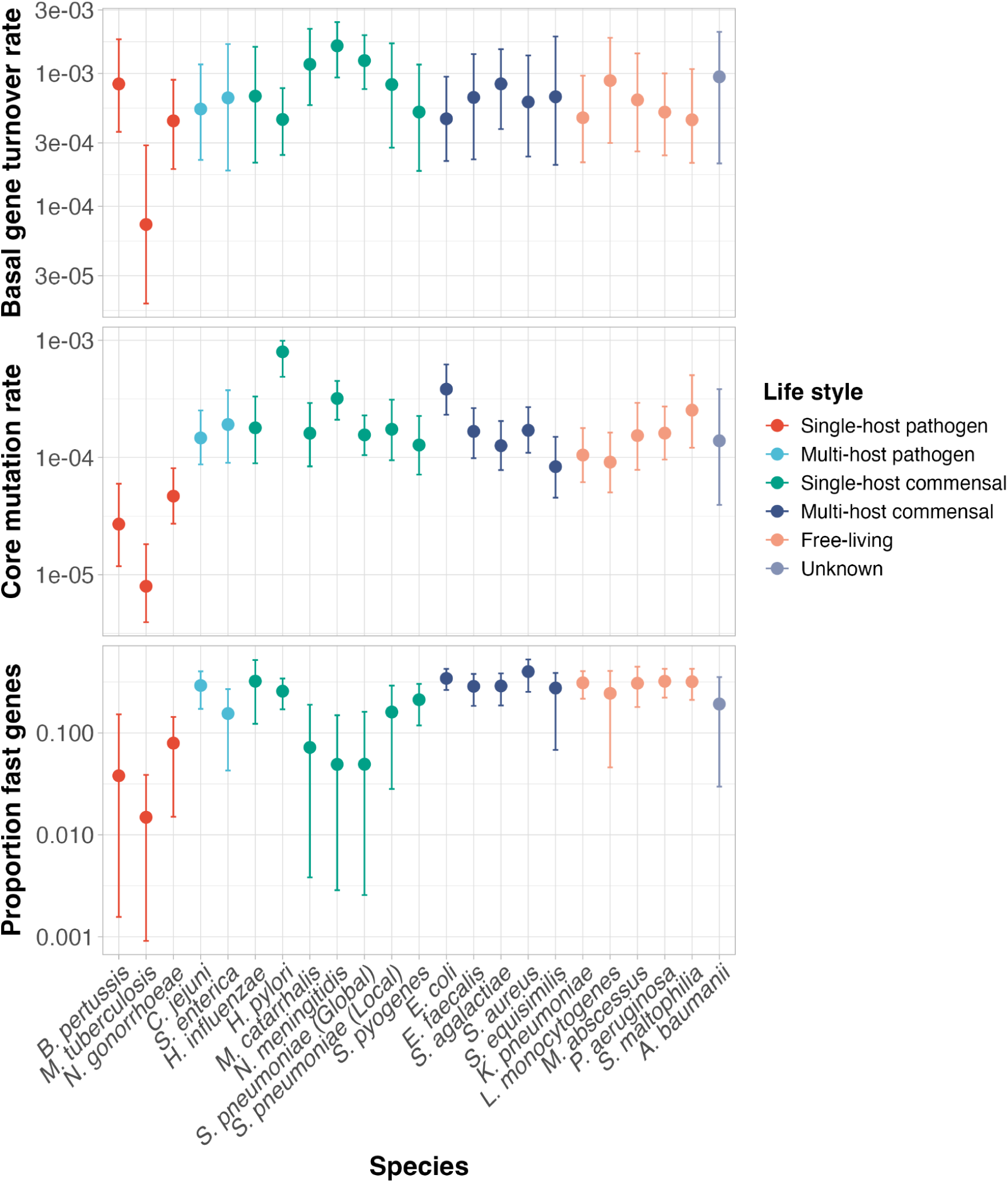
PopPUNK-mod estimated parameter posterior distributions of bacterial genomic data. Points highlight the median of posterior distribution, error bars indicate the 95% credible intervals centred on the median. Basal gene turnover rate and core mutation rate are measured in mutations per site per genome per generation. Proportion of fast genes is measured as the proportion of total accessory genes that are exchanged instantaneously in each generation. All simulations were performed with 1000 individuals and 500 generations. Fixed parameters for model fitting, including core genome size, pangenome size, and average gene frequency are shown in **Supplementary Table 4**. Points are coloured by species lifestyle assignments in **Supplementary Table 1**. Data for each species is available in **Supplementary File 2**. MCMC traces are available in **Supplementary File 3**.

The proportion of fast genes in the accessory genome varied over two orders of magnitude between species. The proportion of fast genes was consistent between species with the same lifestyle, with the exception of some single-host commensals; *M. catarrhalis*, *N. meningitidis* and *S. pneumoniae,* had lower ranges, similar to single-host pathogens. These species also had slightly higher basal gene turnover rates compared to other species, and are naturally competent (Schoen et al. 2009; Santoro et al. 2019; Luke et al. 2004), suggesting that gene turnover is more uniform across the pangenomes of these species due to high rates of recombination.

To understand the relationship between inferred parameters and measures of population diversity, we compared predicted proportions of fast genes to population and pangenome statistics from alternative work on bacterial pangenome dynamics (Bobay and Ochman 2018; Andreani et al. 2017). We observed a significant positive correlation between the proportion of fast genes with effective population size, and a negative correlation with dN/dS (**Supplementary** Figure 14). These correlations suggest that in species with large amounts of standing neutral diversity, their respective pangenomes contain large numbers of highly mobile, potentially deleterious genetic elements. No correlation was present between the proportion of fast genes and measures of gene mobility or gene frequency distributions (**Supplementary** Figures 14 **& 15**).

To analyse model fit performance, we visualised the simulated pairwise distance distributions generated from the fitted parameter ranges for each species, overlaying them onto the observed distances for each species (**Supplementary** Figure 16). For some species, we observed poor fitting of intercept values, corresponding to the proportion of fast genes, due to lack of point density around the origin. Taking *E. coli* as an example of a poor intercept fit, we found fixing the intercept at the value predicted from regression analysis improved recapitulation of the observed distance distributions (**Supplementary** Figures 17 **& 18**). Therefore, combining regression and PopPUNK-mod could provide a means of improving fits for species with lowered origin point density, although this is outside the scope of current work. Additionally, we investigated the effect of sampling bias on our results. As with the regression models, we fitted PopPUNK-mod to the local *S. pneumoniae* dataset from Maela, comparing parameter estimates to the global dataset (Figure 4). We found that credible intervals overlapped between the local and global *S. pneumoniae* datasets, indicating that our approach is more robust to sampling bias compared to use of a regression model.

Overall, we observe stable baseline rates of gene exchange across species, with the main between-species differences appearing in the proportion of rapidly exchanged genes, in line with results in Figure 3.

## Discussion

The observation that pangenomes are maintained over long times in many bacterial species has led to debate over whether the forces driving formation and maintenance of pangenomes are dominated by neutral or selective dynamics (McInerney et al. 2017; Vos and Eyre-Walker 2017; Shapiro 2017). However, previous attempts to model these differences have relied on single species, single data points (e.g. genome length) or population genetics arguments to distinguish these possibilities. These approaches are insufficient to quantitatively determine pangenome evolutionary mechanisms and forces at play within bacteria at large. By combining scalable genomic methods, with a new molecular simulator, we are able to show that interspecies pangenome diversity is driven predominantly by variation in the number of rapidly exchanged genes, whilst the basal rate of gene exchange is conserved.

Based on our analysis of 408 species, we found that core mutation rate and the proportion of fast genes are correlated with the phylogenetic relationship between species, which cannot be distinguished from ecological effects. Lineage-specific bursts of small variant mutagenesis and MGE activity are linked to adaptive evolution in bacterial species, allowing species to explore a wider range of evolutionary trajectories (Consuegra et al. 2021; Arredondo-Alonso et al. 2025; Remigi et al. 2014). Therefore, association of rapidly exchanged genes with specific strains may indicate instances of speciation through diversifying selection. Based on our smaller analysis of 22 pathogenic bacteria, our relative estimates of core and accessory mutation rates broadly agree with known ecological characteristics of the species studied; reproductively isolated pathogens had notably lower mutation rates than multi-host and free living species, which have remarkably similar basal gene turnover rates. These results agree with findings from Collins and Higgs (2012), who show that the rate of gain and loss of slowly exchanged accessory genes varies by one order of magnitude between different species or genera, while the rates in the rapidly exchanged accessory genes vary by two orders of magnitude. However, the identity and function of the rapidly exchanged genes remains unknown, and further work will be required to explore the mechanisms driving differences in gene turnover between species.

Using phylogenetic regression, we found that variation in core mutation rate and the proportion of fast genes does not explain variation in bacterial lifestyle, which is instead largely explained by the phylogeny. This result seemingly disagrees with Dewar et al. (2024), who show pangenome fluidity is strongly associated with bacterial lifestyle, even when accounting for population structure. However, our result agrees with recent comprehensive analysis showing “community conservatism”, whereby closely related species occupy similar niches, is found across bacterial phyla and environments (Malfertheiner et al. 2026). Using Pagel’s λ, we show that a phylogenetic signal does exist for core mutation rate and the proportion of fast genes, meaning that we cannot disentagle the effect of species relatedness when assessing their relationship with bacterial generalism. However, as in Dewar et al. (2024), we found a significant positive correlation between gene mobility and effective population size, which is consistent with previous studies (Bobay and Ochman 2018; Andreani et al. 2017). This overall agreement indicates that our approach is consistently capturing the same underlying pangenome dynamics as observed previously, and is in agreement with evidence of an interaction between species relatedness and ecological niche. However, using our mechanistic model, we are better able to differentiate the contribution of differentially exchanged genes, which is not possible using measures such as pangenome fluidity or effective population size.

We cannot yet say whether differences in gene exchange rates are due to neutral or adaptive processes. The proportion of rapidly exchanged genes correlated with measures of standing diversity, namely effective population size and dN/dS, however, this correlation can have directly contradictory explanations (Shapiro 2017; McInerney et al. 2017). One explanation is that rapidly exchanged genes are largely deleterious and so are quickly removed from a population, while more slowly exchanged genes convey lineage-specific benefits and so are maintained by selection (Collins and Higgs 2012). An alternative explanation is that these compartments may be formed of genes that are all neutrally evolving, with the differences in rates being explained by differences in their own mobility (N’Guessan et al. 2021). However, our results do support the theory that accessory genes are not equal in the forces that govern their frequencies, and that these forces differ by species. Furthermore, lack of correlation between basal gene turnover and the proportion of fast genes suggests exchange of genes in the slow and rapid compartments occurs independently, further supporting the existence of multiple sets of genes with different biological and evolutionary characteristics. As we do not currently fit to time resolved phylogenies, further work will be required to determine the precise timescale of these gene exchanges. Comparing absolute timescales between species will shed light on how pangenome structure is impacted by the time since emergence of a species, and the degree to which genetic drift may play a role in gene frequencies in species which have emerged more or less recently.

Due to issues with parameter identifiability between positive and negative selection, selection parameters could not be reliably fitted using PopPUNK-mod, and so the contribution of neutral or adaptive processes to differences in rates of gene gain and loss in the accessory genome could not be elucidated here. Additionally, within-species differences in the direction of selection, such as one lineage undergoing positive selection while another undergoes negative selection, is not currently captured by our model. Alternative data sources, such as distributions of lineage-specific fitness effects for all genes gathered from experimental mutagenesis experiments or detection of selection in sequence data (Rocha 2018; Eyre-Walker and Keightley 2007; Douglas and Shapiro 2024), will be required to help differentiate between the impact of positive and negative selection on observed pangenome structure. Furthermore, although we show our model is robust to sampling bias, use of unbiased carriage sampling datasets where possible is crucial to ensure capture of all variation in pangenome dynamics present in a species. As we assumed parity in some underlying model parameters for all species, future modelling-based work to determine selective forces acting on bacterial genomes should also take into account variation in species emergence date, differences in population size, differences in gene gain and loss rates, and explicitly model sampling bias.

We developed Pansim in order to capture both neutral and adaptive pangenome dynamics. Pansim represents the first bacterial pangenome simulator which captures both neutral and adaptive processes in the accessory genome, modelling selection effects across the entire accessory genome instead of selective sweeps of single alleles. As a stand-alone tool, Pansim provides a means of modelling the effects of complex interactions between many different pangenome dynamics on observed population diversity, and can be used to generate simulated populations for workflow benchmarking or model training. Furthermore, PopPUNK-mod provides a highly flexible fitting framework for Pansim, enabling fitting of several parameters at once. The novel density-based summary statistic developed here allows Pansim to be fit to pairwise distance data, which can easily be combined with alternative data sources to generate summary statistics that are more informative of selective pangenome dynamics in future iterations.

By combining a mechanistic model of pangenome evolution, with evidence from over 600,000 genomes drawn from over 400 taxonomically diverse bacterial species, we have shown that interspecies differences in pangenome diversity are likely driven by variation in the number of rapidly exchanged genes. These differences are better explained by phylogeny than ecology, and are also inconsistent with all accessory genes being maintained under the same forces, either selective or neutral, as has previously been proposed (Vos and Eyre-Walker 2017; McInerney et al. 2017). Our work instead favours evidence from rare gene pseudogenisation that pangenome dynamics are instead governed by a complex interaction between individual gene mobility and selection pressures that differ according to accessory gene function (Douglas and Shapiro 2024). Further work is required to elucidate whether forces acting upon pangenomes are largely neutral or adaptive, aiding us in our understanding of why bacteria have pangenomes.

## Methods

### Genome assembly dataset collation

For small dataset model fitting, genome assemblies from 22 bacterial species were retrieved from public repositories (**Supplementary Table 1, Supplementary File 1**). Datasets were analysed for each species separately using PopPUNK v2.6.0 (Lees et al. 2019). Initially, datasets were pruned of low quality genomes using the default built-in PopPUNK quality control filters (‘poppunk --create-db --qc-filter prun’). These filters removed genomes with lengths greater than five standard deviations away from the average genome length, greater than 10% ambiguous bases in their assemblies, genomes which have > 5% of their pairwise distances at zero core and accessory values, which are likely duplicate genomes. Genomes gathered from Blackwell et al. (2021) were subjected to an additional requirement of ≥90% of reads being assigned to a single taxon at the species level by Kraken 2 (Wood et al. 2019). All PopPUNK databases were created with default k-mer ranges (k=13-29 with step of 4) and a sketch size of 10,000. There were three exceptions to this parameterisation; *Mycobacterium tuberculosis*, where a sketch size of 50,000 was used, *Salmonella enterica*, where a sketch size of 40,000 was used, and *Acinetobacter baumanii*, where a k-mer range of 17-41 (step of 4) and sketch size of 100,000 was used. Species lifestyle assignments were inferred from literature searches (**Supplementary Table 1**): single- and multi-host species were defined as those able to colonise one or multiple host species respectively; pathogens were defined as obligate pathogenic species; commensals were defined as species that do not always cause disease during host colonisation; free-living species were defined as species able to grow in non-host environments e.g. soil or water; unknown labels were assigned to species with no known natural reservoir.

For *Mycobacterium tuberculosis*, *Streptococcus pneumoniae*, *Escherichia coli* and *Listeria monocytogenes*, core genome phylogenies were generated from PopPUNK core distances (‘poppunk_visualise --microreact’) and visualised with Microreact (Argimón et al. 2016).

For the additional bacterial species analysed, genomes from AllTheBacteria (Hunt et al. 2024) (accessed August 2024) were parsed based on species assignment, with only genomes annotated as “high quality” being used for further analysis. At most 5000 genomes per species were used for PopPUNK analysis, with species with >5000 genomes available being randomly downsampled to 5000 genomes. Species with less than 50 genomes in AlltheBacteria were not included in this study.

Each species dataset was then analysed using an automated quality control pipeline. First, each species dataset was sketched using PopPUNK v2.7.6, using a sketch size of 100,000 and k-mer size range 17bp-41bp. Core and accessory distances calculated by PopPUNK were extracted and ordered respectively, with the 70^th^ percentile being identified for each distance category. Above this distance, a quality control cutoff was determined as the distance that is at least 20% greater than the distance one percentile before it, with all genomes that produce distances higher than this threshold for either the core or accessory distance distribution being removed. The proportion of zero distances (a perfect match between two genomes) was increased to 10% per genome to account for the large number of highly similar genomes in certain species. Code for automated quality control is available on GitHub (https://github.com/anleip/PopPUNK_automation). This resulted in a final dataset of 408 bacterial species for model fitting.

### Phylogenetic analysis and regression

One hundred and four universal CheckM marker genes (Parks et al. 2015) were extracted from a random representative genome selected from each of the 408 bacterial species identified from the AllTheBacteria dataset. Marker gene Hidden Markov Model (HMM) profiles for each marker gene were downloaded from Uniprot (Bateman et al. 2021). Profiles were used to extract marker gene sequences from each genome with PyHMMER v0.11.1 (Larralde and Zeller 2023) using a custom script (https://github.com/gtonkinhill/markers_atb). Gene sequences were concatenated per genome, and aligned using IQtree v3.0.1 (Nguyen et al. 2015) in fast mode with using the LG amino acid substitution matrix (Le and Gascuel 2008) with invariant sites and gamma distributed rate heterogeneity with four rates (‘iqtree--prefix [prefix] -s [alignment] -fast -st AA -m LG+I+G4’). Taxonomic assignment was conducted using GTDB-Tk v2.4.1 (Chaumeil et al. 2022) on representative genome sequences.

The tree was then visualised using phytools v2.5-2 (Revell 2012). Median values of parameters estimated by PopPUNK-mod were added to respective tips as continuous traits. Pagel’s λ (Pagel 1999) was calculated for each trait, which identifies whether a phylogenetic signal is present for associated continuous traits, whereby trait values are more similar between more closely related taxa than between distantly related taxa. Values of λ = 0 and λ > 0 indicate absence or presence of phylogenetic signal for a trait respectively. A p-value is provided for each trait, indicating whether λ significantly deviates from 0.

To analyse the relationship between PopPUNK-mod fitted parameters and bacterial lifestyle, generalism scores were gathered from metaTraits (Podlesny et al. 2025) using the SPIRE database trait catalogue (Schmidt et al. 2024). A total of 85 species possessed a generalism score (**Supplementary File 1**). A linear regression model was built using MCMCglmm v2.36 (Hadfield 2010) by regressing generalism score against each PopPUNK-mod fitted parameter individually, using the phylogeny as a fixed effect (generalism score ∼ parameter, random = ∼Species) as used in Dewar et al. (2024). We report the *p*-value (reported as “pMCMC” by MCMCglmm) of the intercept and coefficient of each linear regression, as well as the R^2^ of each component, made up of the PopPUNK-mod parameter and the phylogeny.

### Non-linear regression analysis

Non-linear regression was performed on core vs. accessory pairwise distances generated by PopPUNK. **Equation 1**, which describes an asymptotic curve, was fitted to pairwise distances using the ‘curve_fit’ function in Scipy (v1.14.1) (Virtanen et al. 2020)

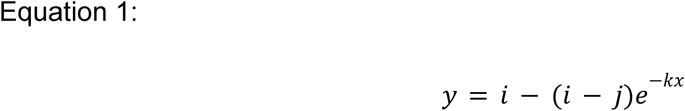

Where:

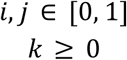

In **Equation 1**, *y* is the accessory distance, *x* is the core distance, *i* is the accessory distance plateau (asymptote), *j* is the y-intercept for the accessory distance, and *k* is the rate ratio describing how quickly the plateau is approached.

### Population dynamics simulations

Simulations were conducted using Pansim v0.1.1, which enables simulation of neutral and selective pangenome dynamics. Pansim implements a Wright-Fisher process (Ewens 2004), where a population is partitioned into X individual cells and proceeds forwards in time with non-overlapping generations to generation N (Figure 2A). Each individual cell contains a core genome, which accrues single nucleotide polymorphisms (SNPs), and an accessory genome, where genes can be gained and lost, in each generation using tau-leaping (Gillespie 2001). In both the core and accessory genomes, changes occur at random in each generation. The total size of the accessory genome across all individuals makes up the pangenome, and is finite in size. Under neutrality, the number of offspring produced by each individual cell in the next generation is uniformly distributed.

Pansim enables sharing of alleles between members of the same generation, either through homologous recombination (HR, pink arrows) in the core genome, or horizontal gene transfer (HGT, purple) arrows in the accessory genome (Figure 2B). The rates of HR and HGT are independent of one another and are independent of the relatedness between individual cells. Pansim also splits gene gain/loss rates across two compartments, slow and fast genes, the size ratio of which can be controlled.

Pansim simulates competition between members of the same generation by calculating pairwise accessory genome distances, and weighting the probability of each cell generating progeny in the next generation by its respective average pairwise distance to all other members of the population, selecting for the most diverged members of the population (Figure 2C).

Pansim models gene fitness distributions as a double exponential distribution about 0, where each gene is assigned a selection coefficient, s, in the range −1 <= s < ∞, which weights the probability of an individual generating progeny by the product of all gene selection coefficients in the accessory genome (Figure 2D).

The assumptions underlying Pansim are as follows. Firstly, all model fitting to observed data used a fixed population size of 1000 and number of generations of 500. These parameters were chosen based on stabilisation of core and accessory distances in simulations above these numbers (**Supplementary Results**), with larger numbers resulting in longer runtime. However, the bacterial species used in model fitting likely do not share these population parameters, with certain species having smaller or larger effective population sizes (Bobay and Ochman 2018), as well as emergence dates much earlier or later than others. Changing population size or the number generation will impact rate parameter estimates which are measured in sites per genome per generation. Furthermore, as we do not use a dated phylogeny during model fitting, generation time is unknown. Therefore, results should be interpreted as relative values between bacterial species, rather than precise rates of mutation.

Secondly, for simplicity of fitting, Pansim assumes gene gain and loss rates are equal, although similar models differentiate these processes (Collins and Higgs 2012; Gamblin et al. 2025). Bacteria exhibit gene deletional bias, with gene loss rate being higher than gene gain rate (Bobay and Ochman 2017). However, we expect this assumption to have limited impact on results, as a gene gain or loss event appears identically when comparing two genomes in a pairwise manner, as conducted when using PopPUNK.

Finally, Pansim does not model sampling bias; however, the datasets used for model fitting are largely pathogen data which will be overrepresentative of disease associated lineages (Blackwell et al. 2021). However, this issue is likely negated by fitted to pairwise distances, as an undersampled lineage will still produce M*N datapoints, where M is the number of genomes belonging to a lineage and N is the total number of genomes in the dataset. If this lineage is very different from other lineages in the population, it will produce a strong signal at high core and accessory distance, even with limited numbers of genomes. We show that sampling bias has limited impact on parameter estimates with comparing results from global and local datasets for *S. pneumoniae* (Figure 4).

Baseline parameter values and the respective units for Pansim simulations are described in **Supplementary Table 2**. Parameter values used in simulation comparisons are provided in **Supplementary Table 3**.

### Pangenome Analysis

Pangenome analysis was performed using WTBCluster v0.1.0. WTBCluster is a Snakemake workflow (Köster and Rahmann 2012), employing Pyrodigal v3.6.3 (Larralde 2022) to predict coding sequences and MMSeqs2 Linclust v16.747c6 (Steinegger and Söding 2017, 2018) for coding sequence clustering, before tokenising genomes into integers, with each integer representing a single gene cluster. Clustering was performed at 50% identity and 50% coverage (coverage mode [‘--cov-mod’]: 0, cluster mode [‘--cluster-mod’]: 2, identity mode [‘--seq-id-mod’]: 2, alignment mode [‘--alignment-mod’]: 3). All coding sequences were clustered in a single step for each species (‘mmseqs2_num_batches’: 1). Core genome size, pangenome size and average gene frequency per genome were then calculated from tokenised genomes using a custom script (‘analyse_pangenome.py’).

### Summary statistics for two-dimensional distance distributions

Pairwise core vs. accessory distance distributions were compared using an approach combining Kernel Density Estimation (KDE) and Jensen-Shannon Distance (JSD) calculation, both implemented individually in Scipy (Virtanen et al. 2020) (Figure 2E). First, the two distance distributions were linearly scaled, so that the value range of the combined distributions was normalised to between 0-1 using the MinMaxScaler in Scikit-Learn (Pedregosa et al. 2011). A KDE was then generated for each 2-dimensional distribution using a bandwidth of 0.03, the Epanechnikov kernel, and the ball-tree algorithm, generating distances in Euclidean space. A square matrix of size 100×100 was then generated, and the values for each KDE distribution were binned to discretise each KDE distribution. To prevent regions of very high density dominating, the matrices were smoothed by raising all values to the power of *x*, where *x* < 1.0 (default *x*=0.25). The matrices were then normalised to between 0-1 and linearised to form a 1-dimensional vector, before their JSD was calculated:

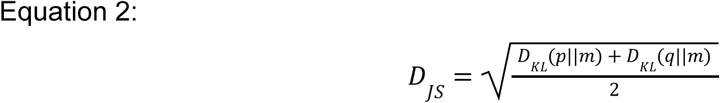

where *D_JS_*is the JSD, *p* and *q* are the vectors being compared, *m* is the pointwise mean of *p* and *q,* and *D_KL_* is the Kullback-Leibler divergence:

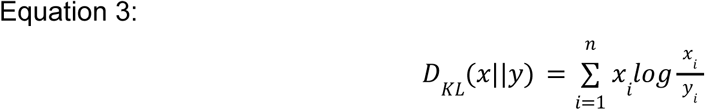

where *x_i_* and *y_i_* are the densities at bin *i* between two vectors *x* and *y* of length *n*.

### Model fitting using Approximate Bayesian Computation

Model fitting was performed using PopPUNK-mod v0.2.2. Pairwise distances were extracted from PopPUNK fits for individual species using a custom script (‘poppunk_extract_distances.py’) during fitting to observed data, or generated by Pansim during fitting to simulated data. Distances were downsampled to 100,000 entries to reduce computational workload. BOLFI Approximate Bayesian Computation (ABC) (Gutmann and Corander 2015; Lintusaari et al. 2018) within PopPUNK-mod, was then used to fit Pansim simulations to downsampled pairwise distances. PopPUNK-mod can be used to fit any combination of parameters available in Pansim. During fitting, the observed data (target distribution) is compared to simulated data generated by Pansim across a range of parameterisations using the KDE-JSD approach described above, as well as inclusion of relative proportions of core, intermediate and rare genes in each pangenome. Core genes were identified as clusters of orthologous genes with frequency ≥95% of all genomes in a population, rare genes as ≥1% and <5% frequency, and intermediate genes as ≥5% and < 95% frequency. Distances between simulated (*u*) and observed (*v*) data are calculated using a euclidean distance given by:

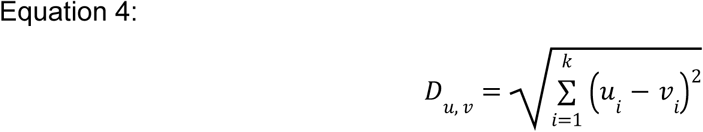

where parameters *i* to *k* denote the JSD, and proportions of core, intermediate and rare genes. Based on the euclidean distances, BOLFI-ABC generates a discrepancy distribution (Figure 2F), fitting a gaussian process which is explored using MCMC. In this work, a Gaussian process was fitted using an RBF kernel with covariance for each parameter set at 10% of the maximum parameter value. MCMC was conducted using the Metropolis-Hastings algorithm for 100,000 samples with 50,000 sample burn-in.

Parameter posterior distributions were generated for all fitted parameters. For fitting to simulated data, ground-truth values were compared to 95% posterior credible intervals. If the ground-truth value sat within the 95% posterior credible intervals, this was deemed a true positive, and a false negative if not. Precision was determined by determining the range of the credible intervals, with a smaller range indicating higher precision.

## Data availability

Pansim v0.1.1 is available on Github (https://github.com/bacpop/Pansim/releases/tag/v0.1.1) and on Zenodo (doi: 10.5281/zenodo.17976329). PopPUNK-mod v0.2.2 is available on Github (https://github.com/samhorsfield96/PopPUNK-mod/releases/tag/v0.2.2) and on Zenodo (doi: 10.5281/zenodo.18402068). WTBCluster v0.1.0 is available on Github (https://github.com/samhorsfield96/WTBcluster/releases/tag/v0.1.0). Pairwise genome distances for all species described in Figures 3 **& 4** are available on Zenodo (doi: 10.5281/zenodo.18129934).

## Conflict of interest

None declared.

## Funding

This work was supported by the European Molecular Biology Laboratory, European Bioinformatics Institute. S.T.H. was funded by the MRC Centre for Global Infectious Disease Analysis (studentship grant ref.: MR/S502388/1), jointly funded by the UK Medical Research Council (MRC) and the UK Foreign, Commonwealth and Development Office (FCDO), under the MRC/FCDO Concordat agreement and is also part of the EDCTP2 program supported by the European Union. For the purpose of open access, the authors have applied a Creative Commons Attribution (CC BY) license to any Author Accepted Manuscript version arising from this submission.

## Author Contributions

Conceptualization: S.T.H., N.J.C., and J.A.L. Methodology: All authors. Software: S.T.H. Validation: S.T.H. Formal analysis: S.T.H. Investigation: S.T.H. Resources: S.T.H., N.J.C. and J.A.L. Data curation: S.T.H. Writing - original draft: S.T.H. Writing - review and editing: All authors. Visualization: S.T.H. Supervision: N.J.C. and J.A.L. Funding acquisition: N.J.C. and J.A.L.

## Notes

https://github.com/bacpop/Pansim/releases/tag/v0.1.1

https://github.com/samhorsfield96/PopPUNK-mod/releases/tag/v0.2.2

https://github.com/samhorsfield96/WTBcluster/releases/tag/v0.1.0

https://zenodo.org/records/18129934

### Competing Interest Statement

The authors have declared no competing interest.

