## Supplementary Material for "Rapid gene exchange explains differences in bacterial pangenome structure"

### Supplementary Figures

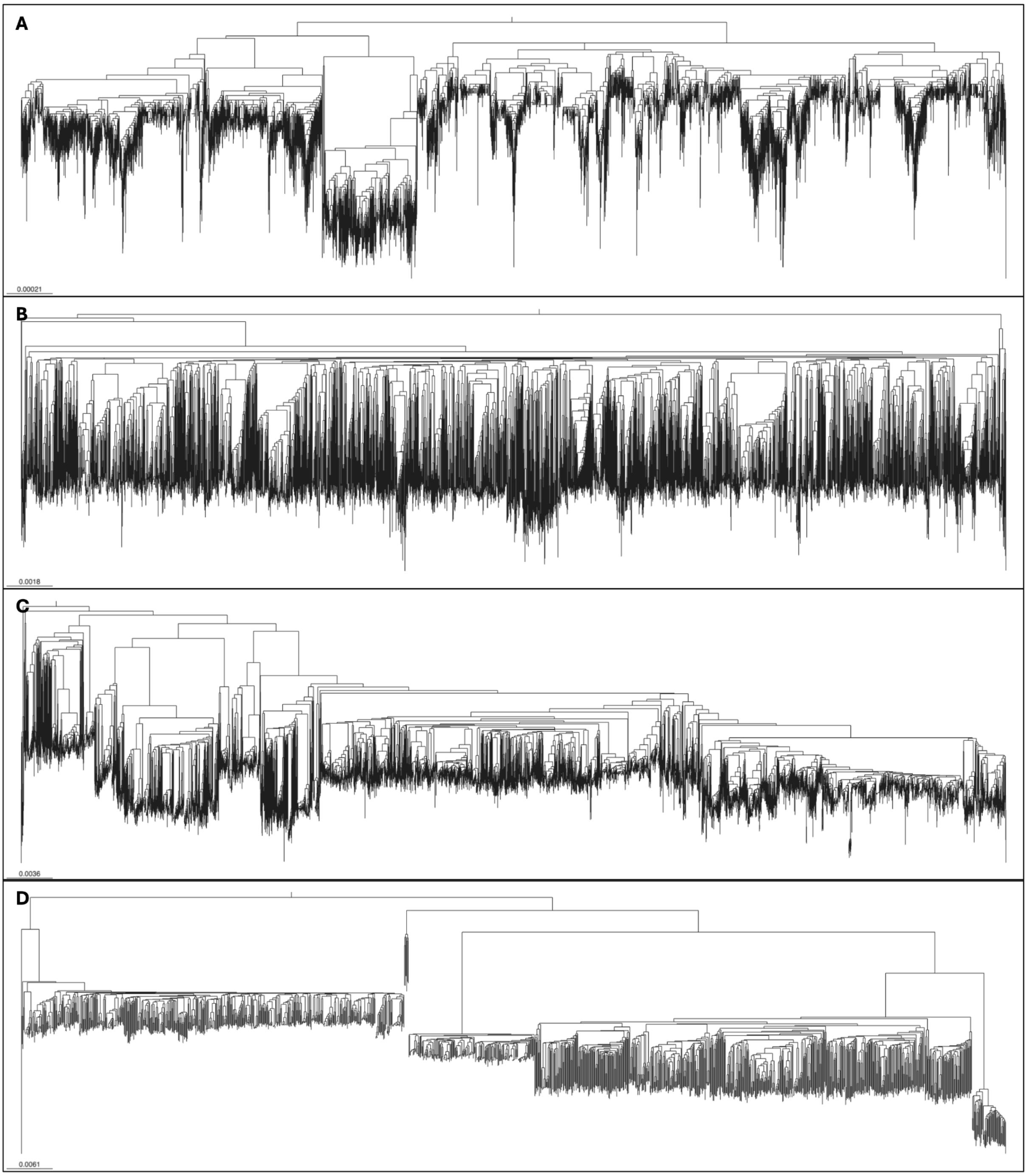


Supplementary Figure 1: Neighbour joining tree of PopPUNK core distances from (**A**) *Mycobacterium tuberculosis*, (**B**) *Streptococcus pneumoniae*, (**C**) *Escherichia coli* and (**D**) *Listeria monocytogenes*.

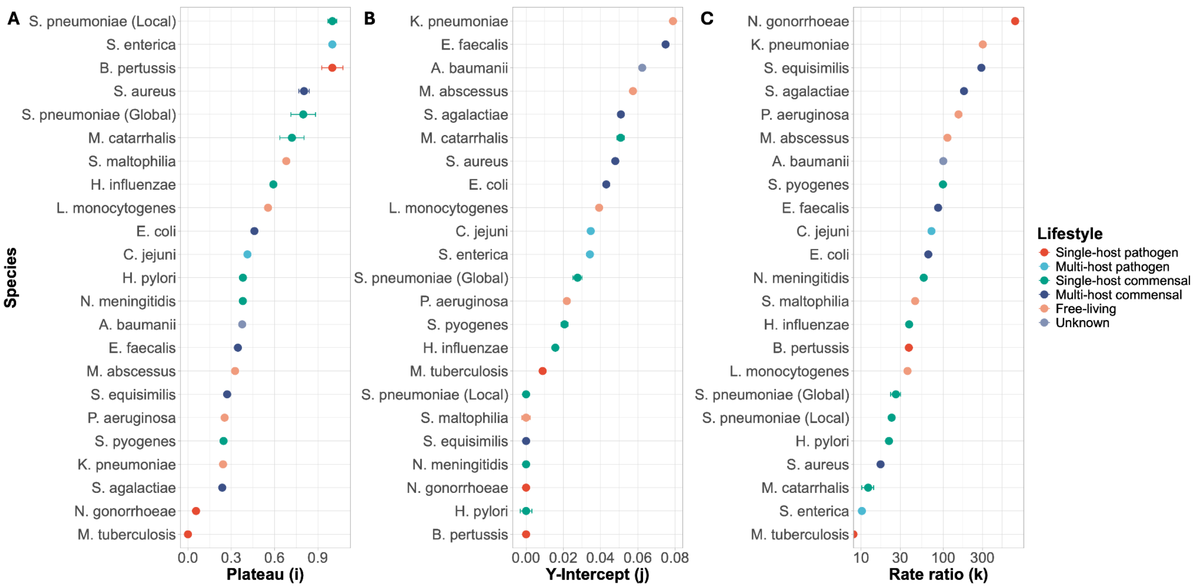


Supplementary Figure 2: Comparison of parameters values from negative exponential fits to PopPUNK pairwise core vs. accessory distances for 22 bacterial species. Panels highlight values for (**A**) plateau (*i*), (**B**) y-intercept (*j*) and (**C**) rate ratio (*k*). 95% confidence intervals are represented as error bars. Points are coloured by species lifestyle.

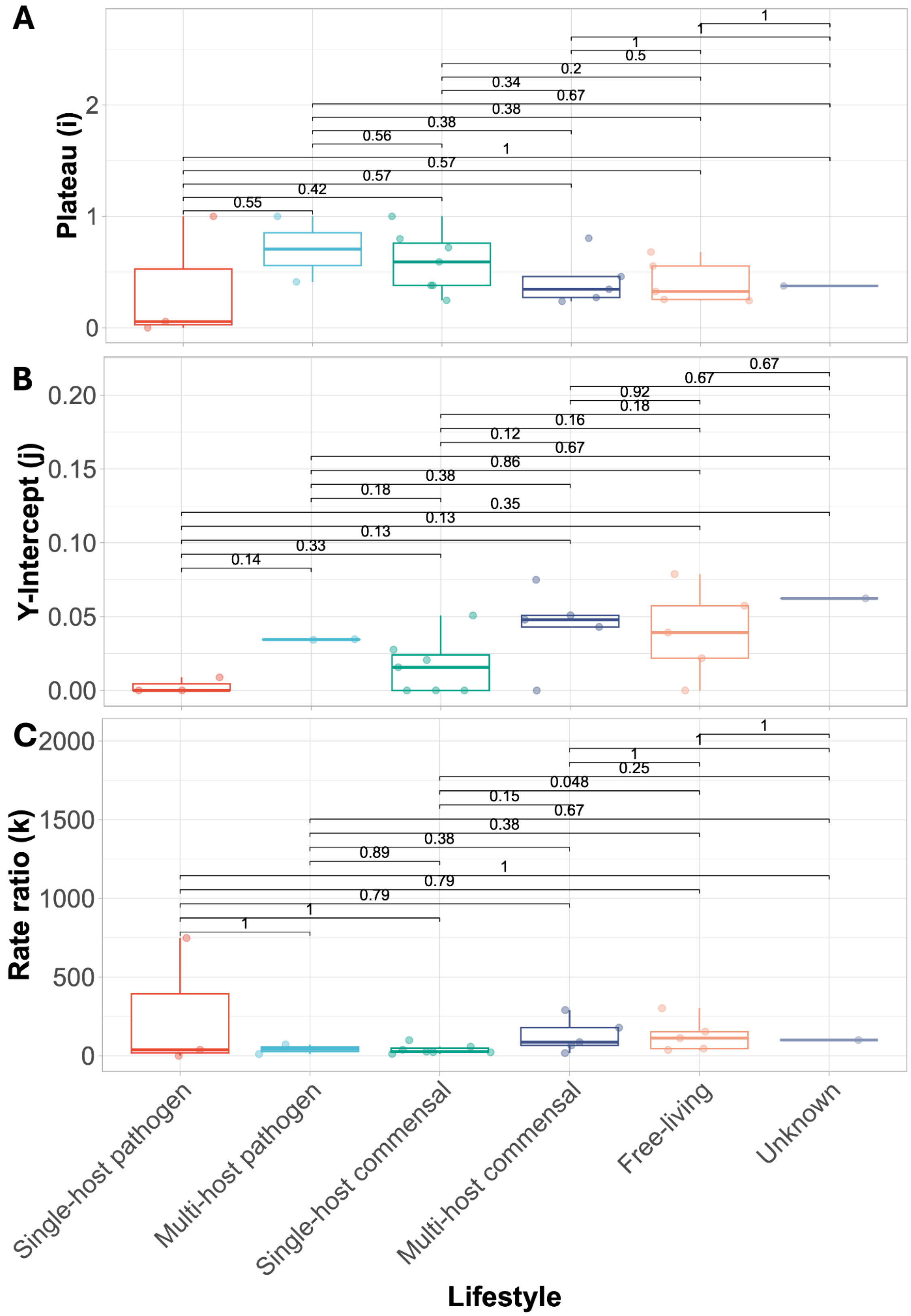


Supplementary Figure 3: Statistical comparison of parameters values from negative exponential fits to PopPUNK pairwise core vs. accessory distances for 22 bacterial species by lifestyle. Panels highlight values for (**A**) plateau (*i*), (**B**) y-intercept (*j*) and (**C**) rate ratio (*k*). Statistical comparisons were conducted using unpaired Wilcoxon tests, with p-values indicated above pairwise comparisons. *p*-values were corrected for multiple comparisons using the Bonferroni correction method.

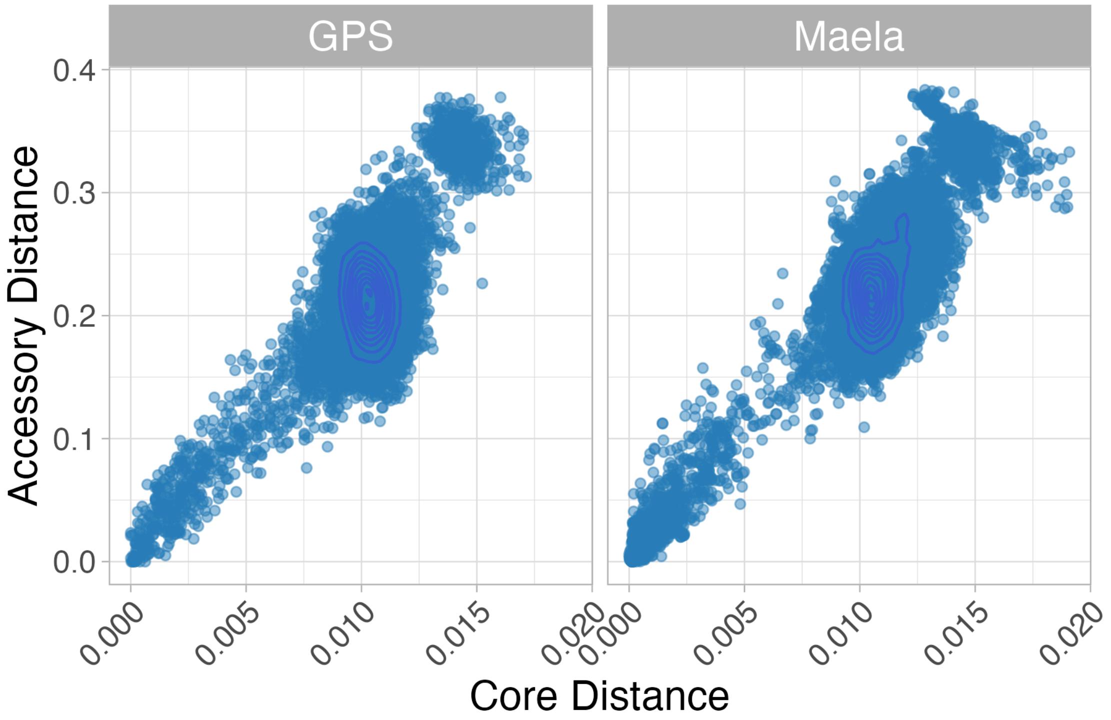


Supplementary Figure 4: PopPUNK distance distributions of *S. pneumoniae* from the Global Pneumococcal Sequencing (GPS) project and local sampling in the Maela refugee camp. Each point is a single pairwise comparison. Contours describe point density.

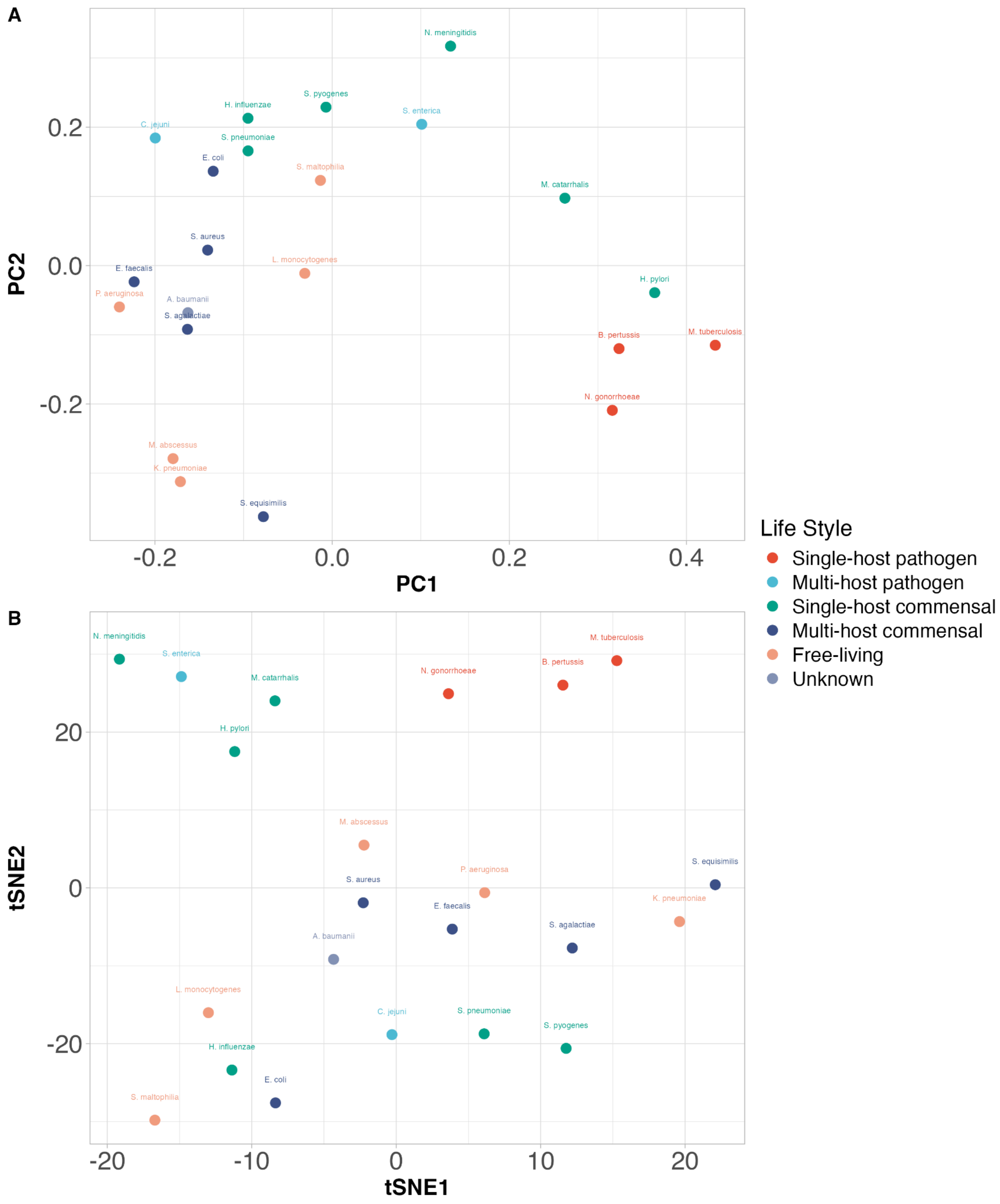


Supplementary Figure 5: PCA (A) and tSNE (B) of pairwise Jensen-Shannon distances between PopPUNK core vs. accessory distance distributions of bacterial species. Points are coloured by species lifestyle. Each point is labelled by its respective species.

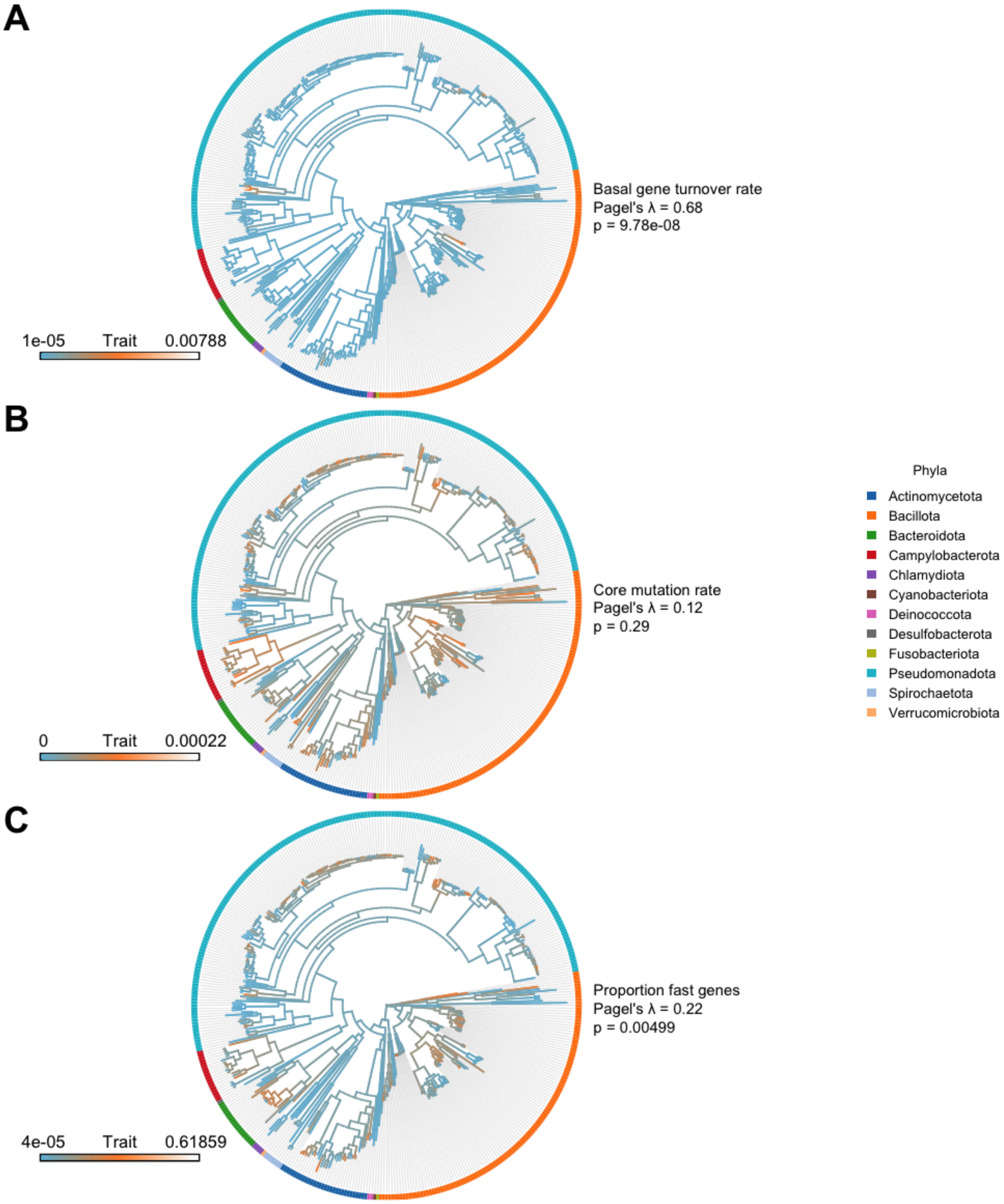


Supplementary Figure 6: Phylogenetic signal analysis of lower credible interval parameter values. Model parameters used were (**A**) basal gene turnover rate, (**B**) core mutation rate and (**C**) proportion of fast genes. Parameter units detailed in **Supplementary Table 2**. Branches are coloured according to parameter value from blue (low) to orange/white (high), denoted by trait scale in bottom left of each plot. Pagel's $\lambda$ was used to calculate the degree of correlation between each parameter and the underlying phylogeny, and is displayed per parameter along with respective p-values calculated using phytools. Phyla designations were assigned from GTDB. Data for each species is available in **Supplementary File 1**.

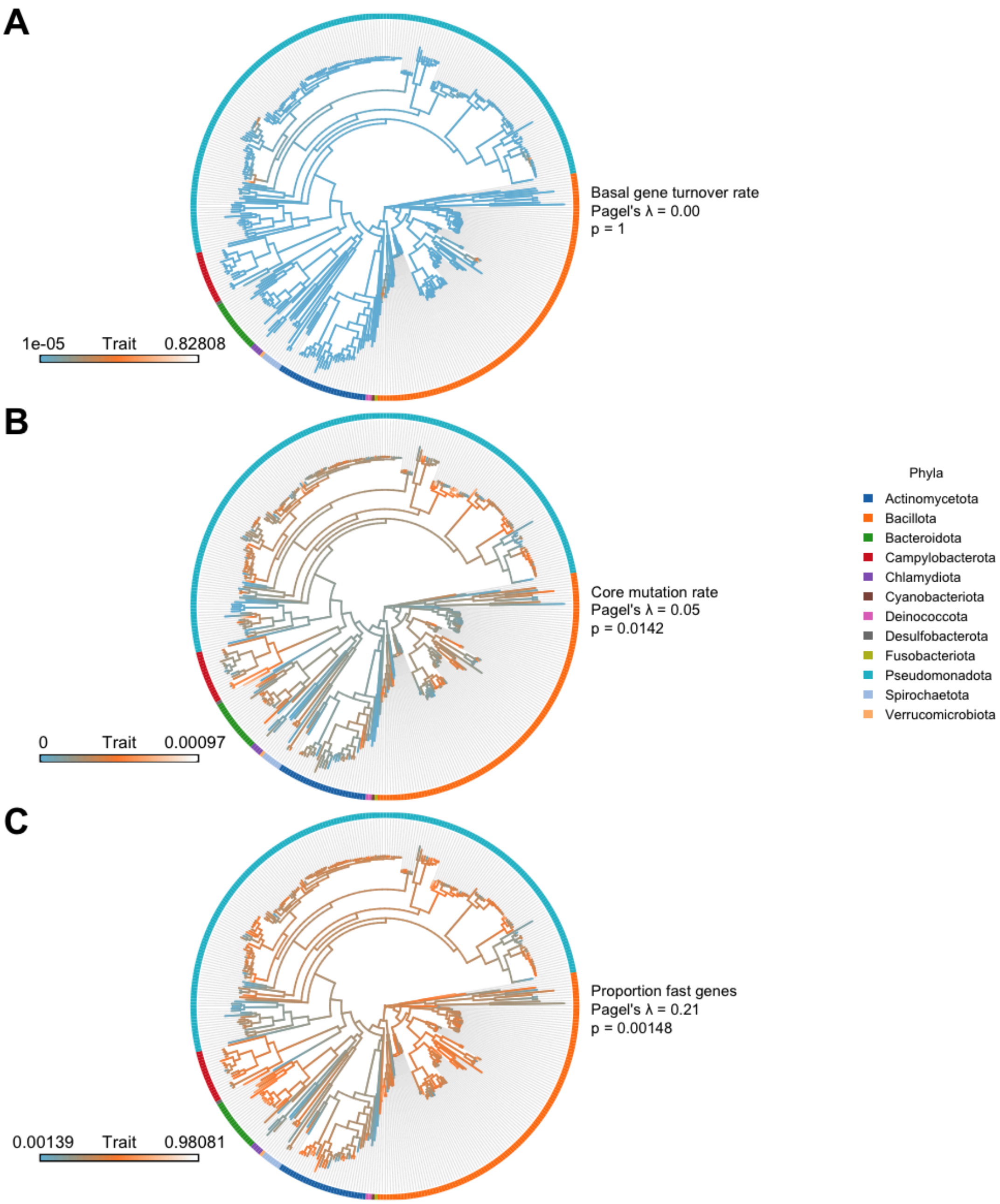


Supplementary Figure 7: Phylogenetic signal analysis of upper credible interval parameter values. Model parameters used were (**A**) basal gene turnover rate, (**B**) core mutation rate and (**C**) proportion of fast genes. Parameter units detailed in **Supplementary Table 2**. Branches are coloured according to parameter value from blue (low) to orange/white (high), denoted by trait scale in bottom left of each plot. Pagel's $\lambda$ was used to calculate the degree of correlation between each parameter and the underlying phylogeny, and is displayed per parameter along with respective p-values calculated using phytools. Phyla designations were assigned from GTDB. Data for each species is available in **Supplementary File 1**.

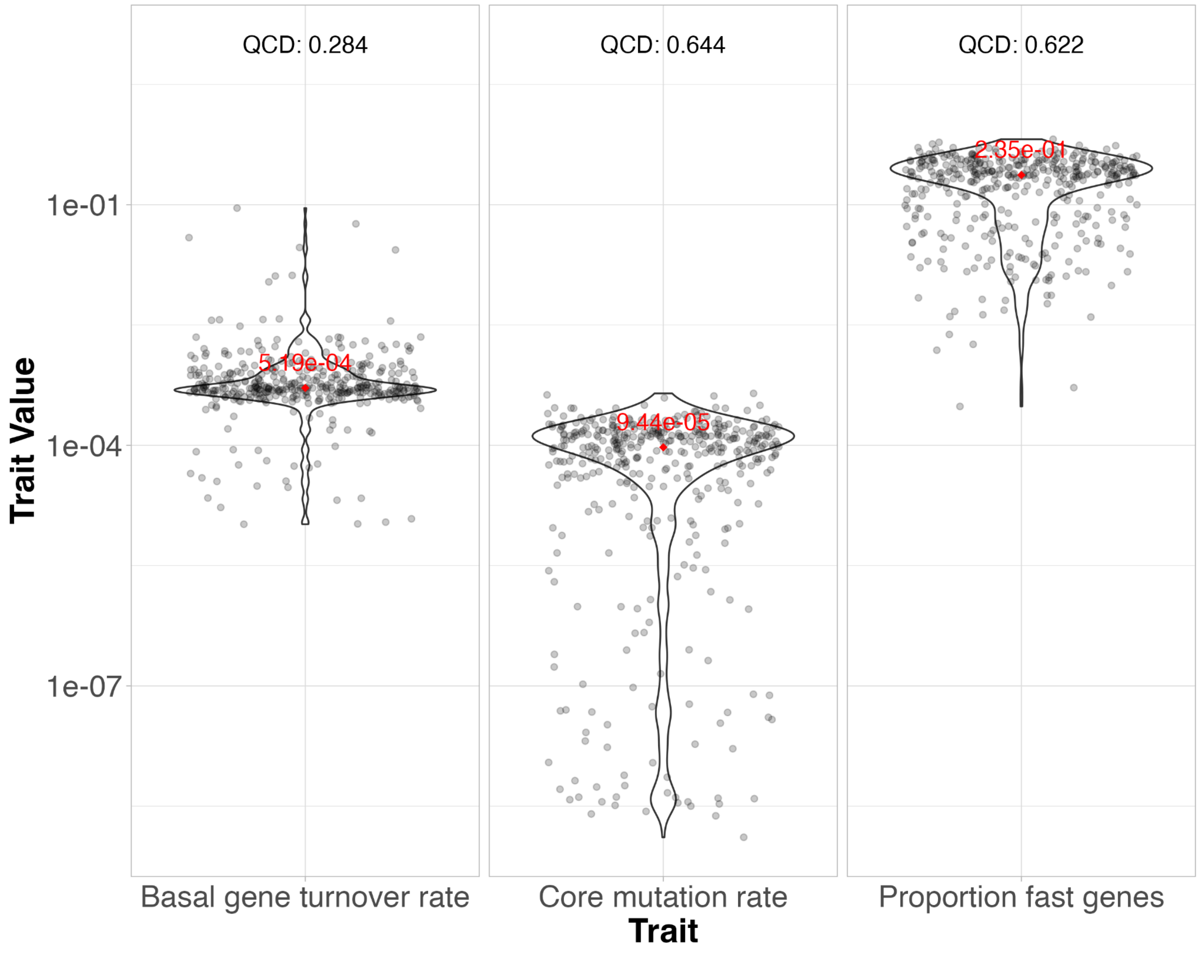


Supplementary Figure 8: Parameter estimate distributions for 408 species from AllTheBacteria. Medians of each distribution are annotated in red. The Quartile Coefficient of Variation (QCD) is annotated at the top of each facet, calculated as (Q3 - Q1) / (Q3 + Q1), where Q1 and Q3 are the 1$^{st}$ and 3$^{rd}$ quartiles respectively. Units for each trait are available in **Supplementary Table 2**.

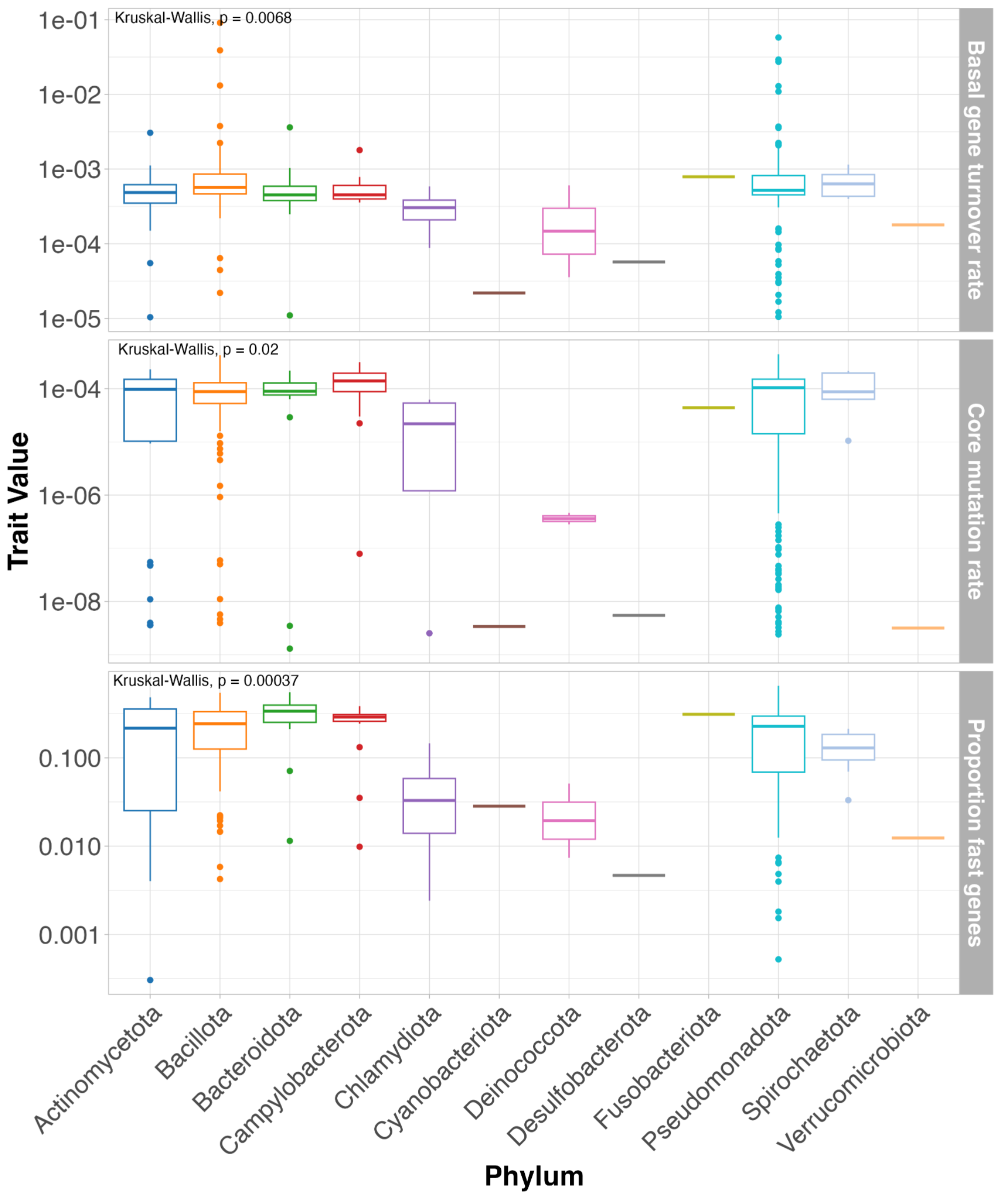


Supplementary Figure 9: Comparison of PopPUNK-mod median parameter estimates across phyla from 408 species from AllTheBacteria. Statistical tests carried out using the Kruskal-Wallis test.

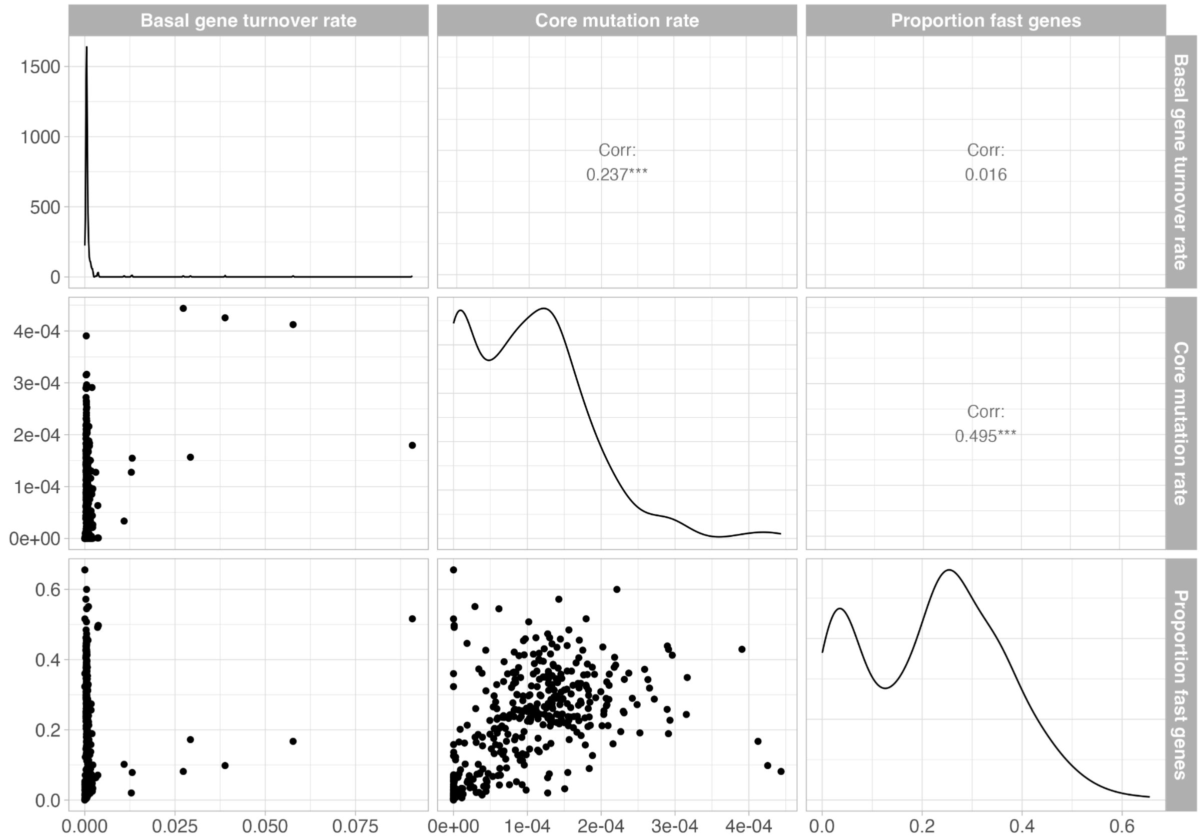


Supplementary Figure 10: Correlation of median posterior estimates of 408 species from AllTheBacteria. Scatter plots shown between each parameter value pair. Density plots show distribution of each parameter value. Pearson correlation coefficient shown for each pairwise comparison with *p*-value (*** denotes p < 0.001). Units for each trait are available in **Supplementary Table 2**.

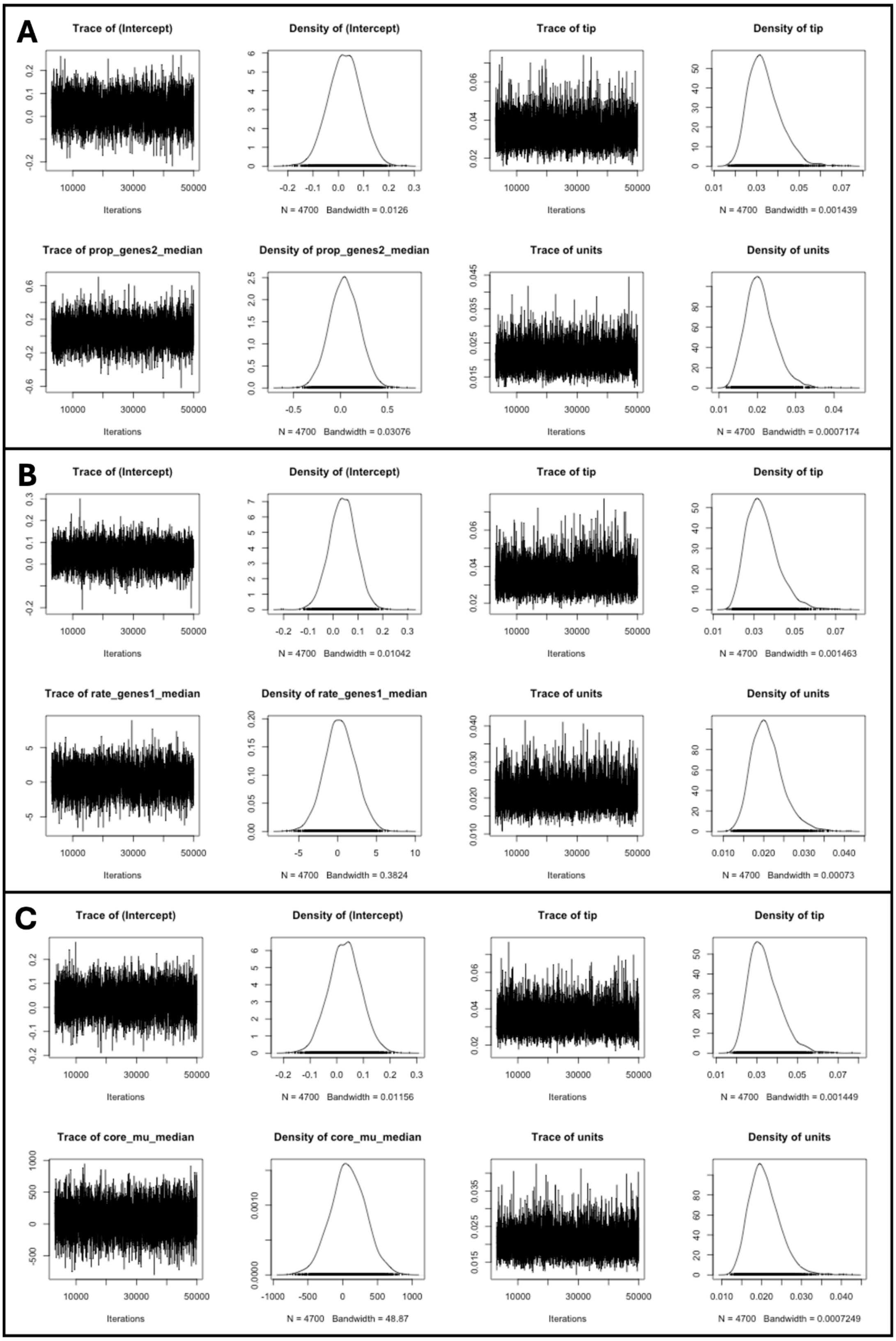


Supplementary Figure 11: Markov Chain Monte Carlo (MCMC) traces and posterior distributions from linear models of generalism score vs. PopPUNK-mod fitted parameters using MCMCglmm. Panels describe PopPUNK-mod parameter used in regression: (**A**) proportion of fast genes, (**B**) basal gene turnover rate, (**C**) core mutation rate. Plots in each panel: top left, intercept fit; bottom left, coefficient fit; top right, phylogeny as random effect; bottom right; error term as random effect.

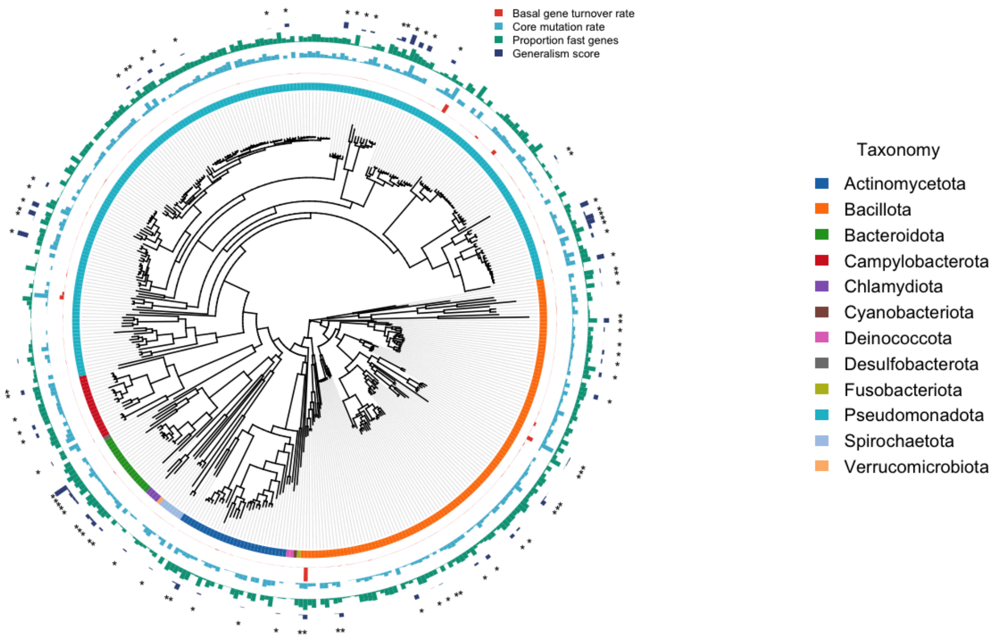


Supplementary Figure 12: Phylogenetic tree of 408 species from AllTheBacteria annotated with parameter estimates from PopPUNK-mod and generalism score from metaTraits. Ring annotation moving from inside to outside: phylum assignment, median basal gene turnover rate, median core mutation rate, median proportion of fast genes, generalism score. The 85 species with generalism scores are annotated with asterisks on the outermost layer.

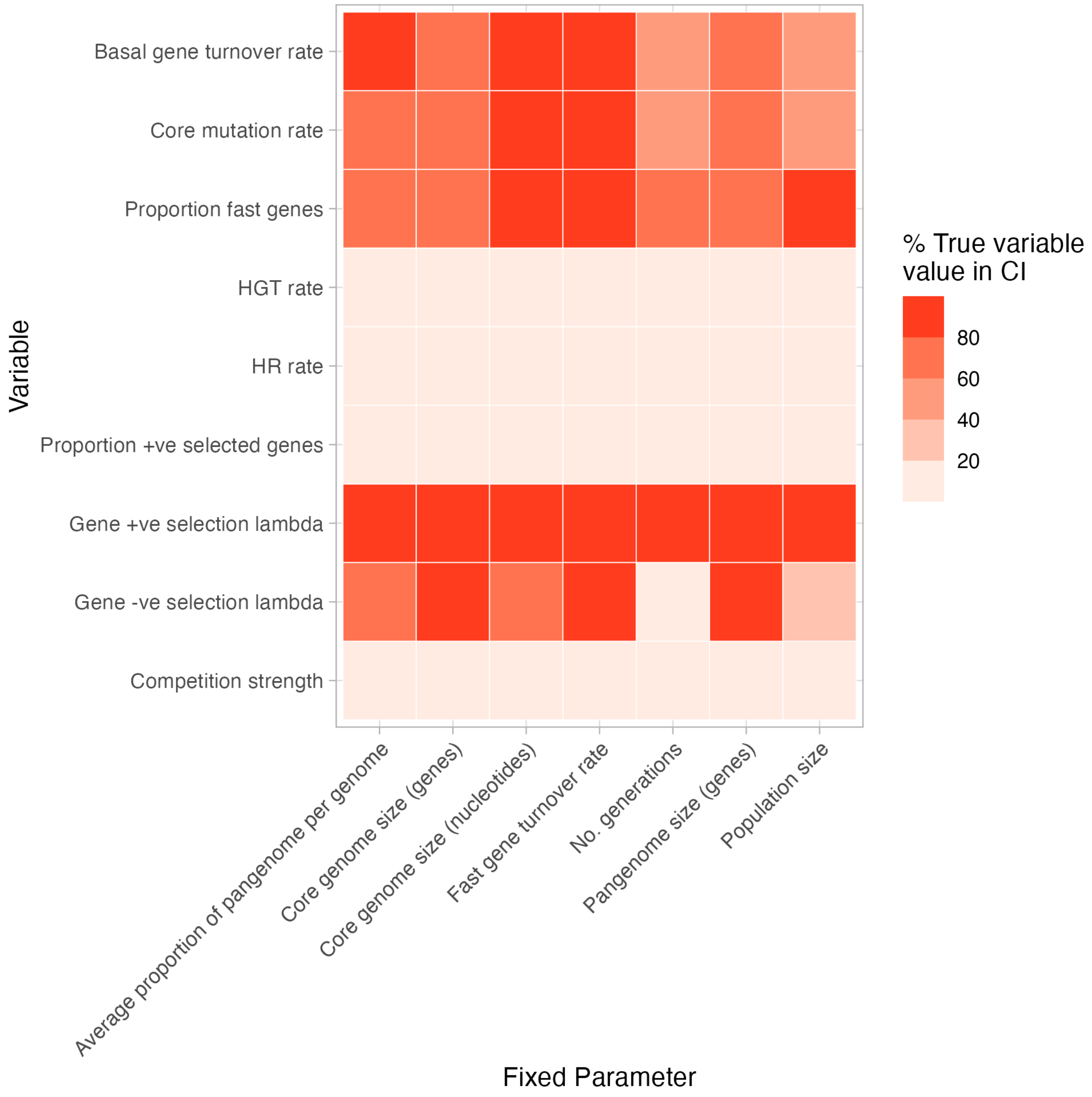


Supplementary Figure 13: Sensitivity analysis of fixed parameters on estimation of ground truth fitted parameters. All ground truth variable parameters were set to defaults in **Supplementary Table 2**.

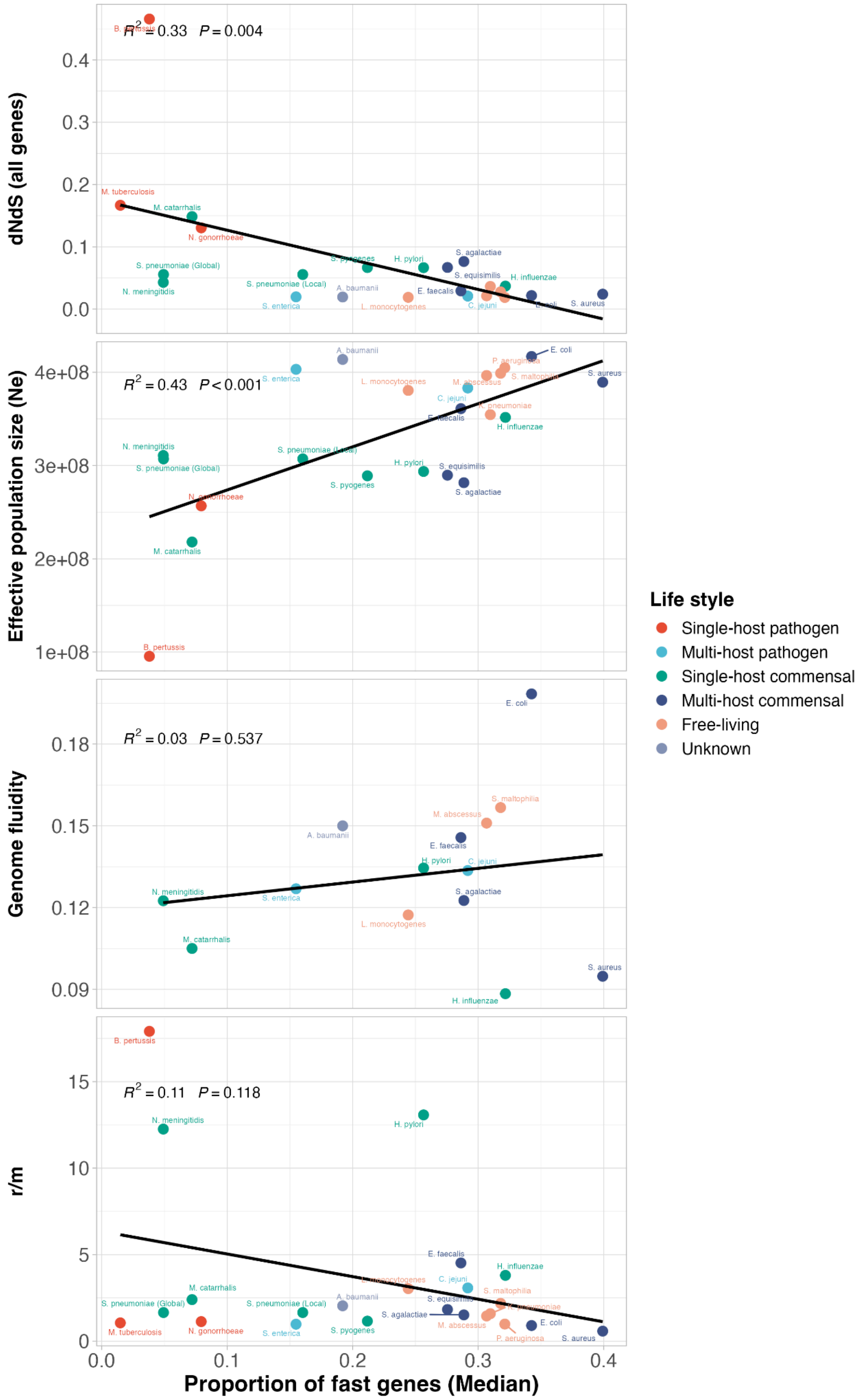


Supplementary Figure 14: Correlation of proportion of fast genes median value estimated by PopPUNK with population statistics. First panel; dN/dS of all genes in pangenome from Bobay and Ochman (2018). Second panel; effective population size from Bobay and Ochman (2018). Third panel; Genome fluidity from Andreani et al. (2017). Fourth panel panel; r/m, number of SNPs resulting from recombination over background mutations from Bobay and Ochman (2018). $R^2$ and coefficient p-values from linear regression.

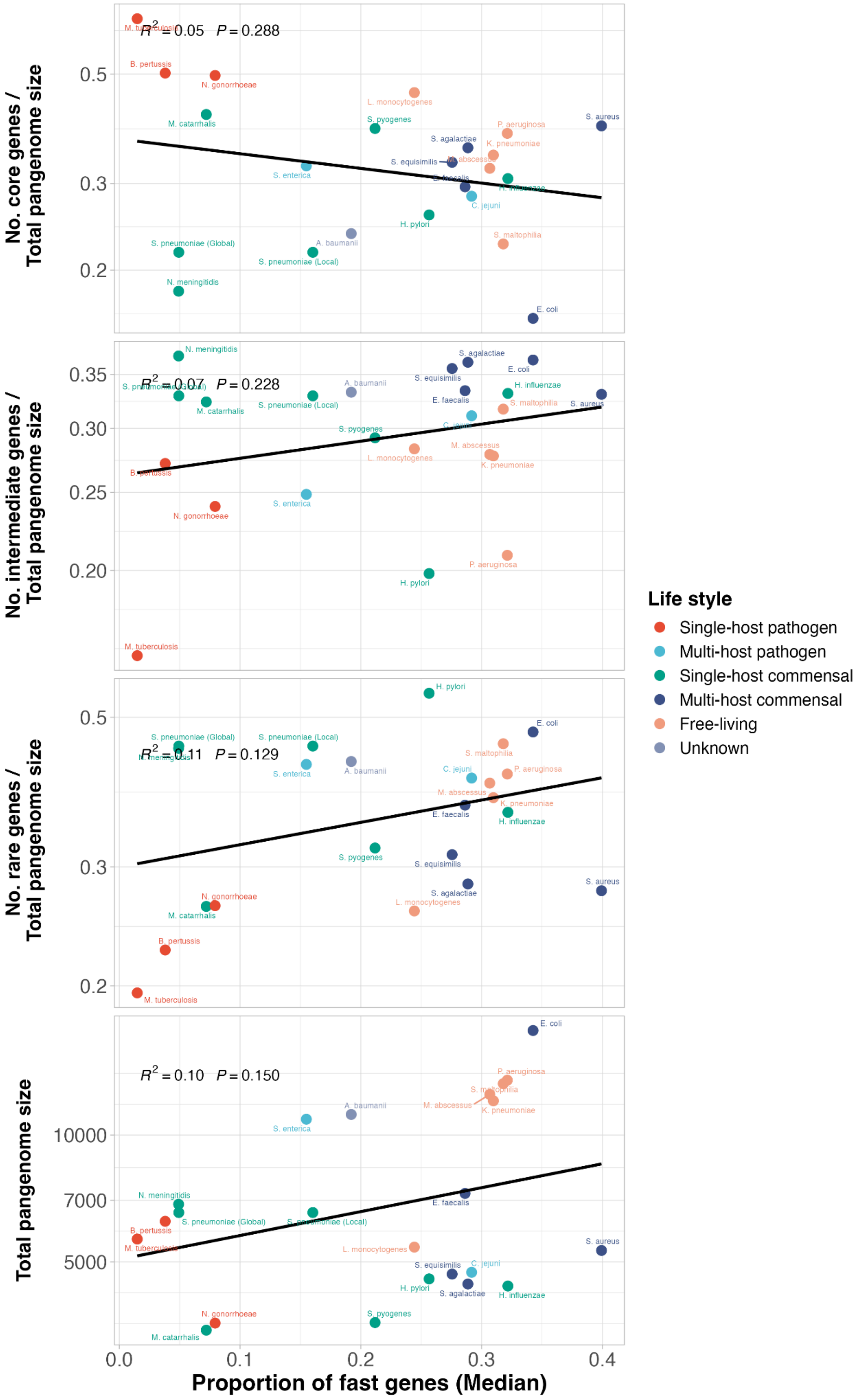


Supplementary Figure 15: Correlation of proportion of fast genes median value estimated by PopPUNK with pangenome statistics. First panel; fraction of core genes in pangenome. Second panel; fraction of intermediate frequency genes in pangenome. Third panel; fraction of rare genes in pangenome. Fourth panel panel; total genes in pangenome. $R^2$ and coefficient p-values from linear regression.

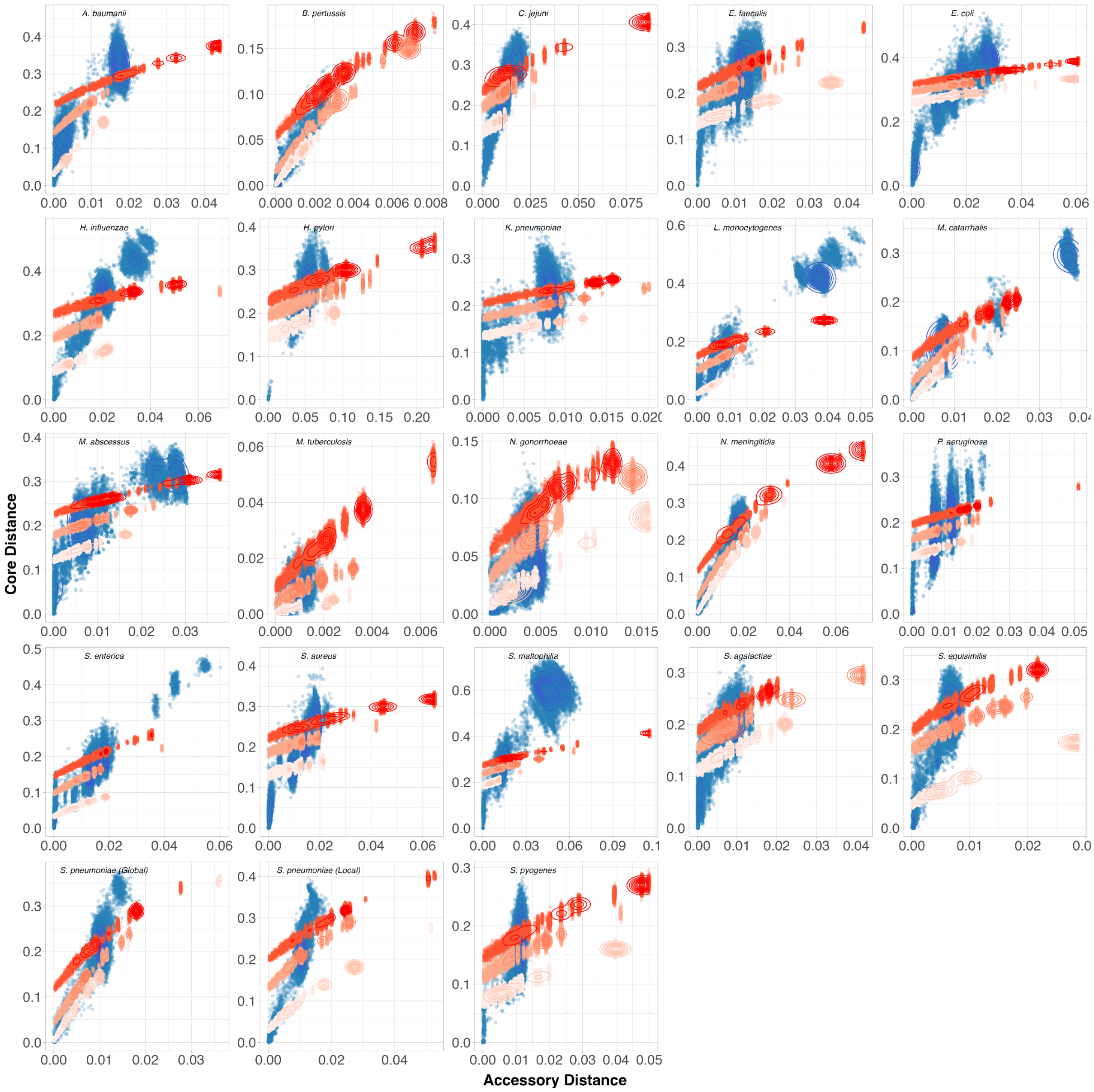


Supplementary Figure 16: Real pairwise genome distances (blue) and Pansim distances (red) based on median parameters estimated from PopPUNK-mod in **Figure 4**. Moving from light, medium and dark red represents Pansim run with lower credible interval, median and upper credible interval parameter values respectively. Each data point represents a pairwise comparison between two genomes.

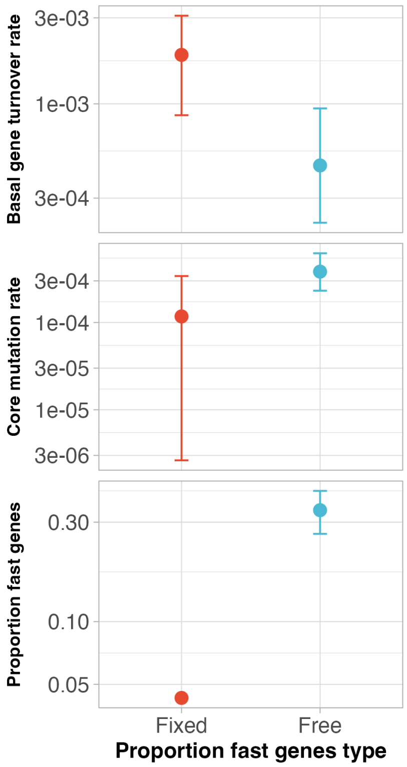


Supplementary Figure 17: Comparison of fitted parameters with and without fixing proportion of fast genes to parameter estimated using non-linear regression (value = 0.043, all values in **Supplementary File 4**).

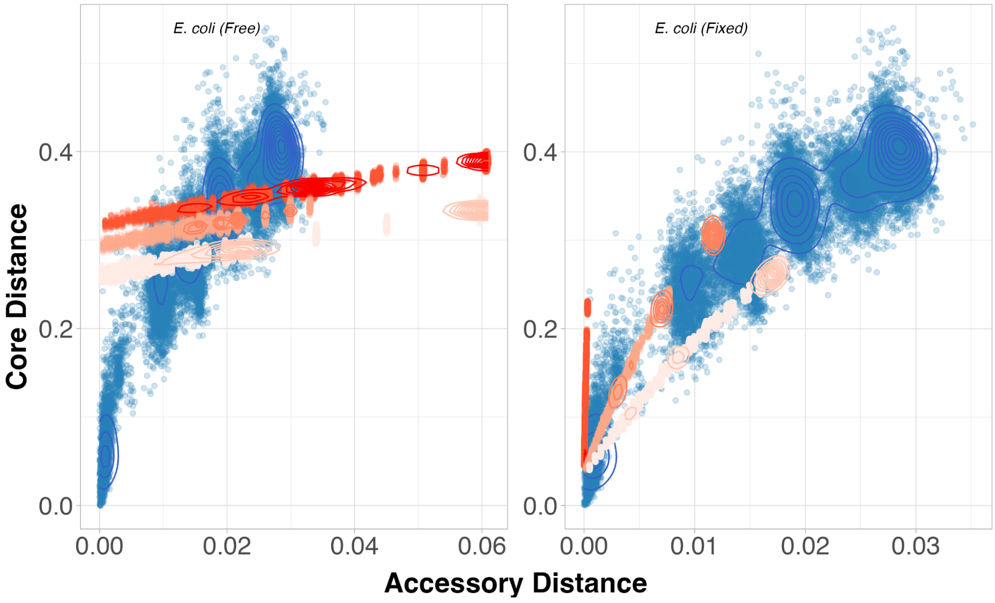


Supplementary Figure 18: Comparison of Pansim pairwise distance distributions with and without fixing proportion of fast genes to parameter estimated using non-linear regression. Blue points represent distances from observed data, red represents Pansim distances. Moving from light, medium and dark red represents Pansim run with lower credible interval, median and upper credible interval parameter values respectively.

### Supplementary Tables

Supplementary Table 1: Number, source of genomes and lifestyle categorisation from 22 bacterial species used in analysis.

| Species/Dataset | No. Genomes | Source | Lifestyle | Lifestyle Citation |
|---|---|---|---|---|
| *Acinetobacter baumannii* | 8263 | (Sayers et al. 2021) | Unknown | (Antunes et al. 2014) |
| *Bordetella pertussis* | 339 | (Blackwell et al. 2021) | Single-host pathogen | (Preston 2005) |
| *Campylobacter jejuni* | 40303 | (Timme et al. 2019) | Multi-host pathogen | (Humphrey et al. 2014) |
| *Enterococcus faecalis* | 2026 | (Pöntinen et al. 2021) | Multi-host commensal | (Pöntinen et al. 2021; Hammerum 2012) |
| *Escherichia coli* | 30138 | (Horesh et al. 2021; Timme et al. 2019) | Multi-host commensal | (Tenaillon et al. 2010) |
| *Haemophilus influenzae* | 89 | (Blackwell et al. 2021) | Single-host commensal | (Erwin and Smith 2007) |
| *Helicobacter pylori* | 2201 | (Sayers et al. 2021) | Single-host commensal | (Reshetnyak et al. 2021; Mladenova-Hristova et al. 2017) |
| *Klebsiella pneumoniae* | 1361 | (David et al. 2019) | Free-living | (Rocha et al. 2022) |
| *Listeria monocytogenes* | 40474 | (Timme et al. 2019) | Free-living | (Nowak et al. 2024) |
| *Moraxella catarrhalis* | 209 | (Sayers et al. 2021) | Single-host commensal | (Karalus and Campagnari 2000) |
| *Mycobacterium abscessus* | 861 | (Ruis et al. 2021) | Free-living | (Lagune et al. 2024) |
| *Mycobacterium tuberculosis* | 13170 | (Blackwell et al. 2021) | Single-host pathogen | (Mohammadnabi et al. 2024) |
| *Neisseria gonorrhoeae* | 8801 | (Blackwell et al. 2021) | Single-host pathogen | (Lenz and Dillard 2018) |
| *Neisseria meningitidis* | 33379 | (Jolley et al. 2018) | Single-host commensal | (Ladhani et al. 2020; Mikucki et al. 2022) |
| *Pseudomonas aeruginosa* | 6082 | (Blackwell et al. 2021) | Free-living | (Folkesson et al. 2012) |
| *Salmonella enterica* subsp. *enterica* | 48116 | (Timme et al. 2019) | Multi-host pathogen | (Winfield and Groisman 2003) |
| *Staphylococcus aureus* | 42117 | (Petit and Read 2018) | Multi-host commensal | (Krismer et al. 2017; Haag et al. 2019) |
| *Stenotrophomonas maltophilia* | 1210 | (Gröschel et al. 2020) | Free-living | (Denet et al. 2018) |
| *Streptococcus* | 18029 | (Jolley et al. 2018) | Multi-host | (Da Cunha et al. 2014; |

| | | | | |
|---|---|---|---|---|
| *agalactiae* | | | commensal | Ren et al. 2024) |
| *Streptococcus dysgalactiae* subsp. *equisimilis* | 501 | (Sayers et al. 2021) | Multi-host commensal | (Haidan et al. 2000; Porcellato et al. 2021) |
| *Streptococcus pneumoniae* (Global) | 42157 | (Gladstone et al. 2019) | Single-host commensal | (Brooks and Mias 2018) |
| *Streptococcus pneumoniae* (Local) | 3069 | (Chewapreecha et al. 2014) | Single-host commensal | (Brooks and Mias 2018) |
| *Streptococcus pyogenes* | 2084 | (Davies et al. 2019) | Single-host commensal | (Armitage et al. 2024; Bessen 2009) |
| AllTheBacteria | 271939 | (Hunt et al. 2024) | N/A | N/A |

Supplementary Table 2: Baseline parameter values for Pansim simulations.

| Parameter | Argument | Value | Units |
|---|---|---|---|
| Population size | pop_size | 1000 | Individuals |
| Core nucleotides | core_size | 1,000,000 | Nucleotides |
| Pangenome size | pan_genes | 5000 | Genes |
| Core genome size | core_genes | 1000 | Genes |
| Average gene frequency | avg_gene_freq | 0.25 | Proportion |
| Number of generations | n_gen | 100 | Generations |
| Maximum pairwise distances | max_distances | 100000 | Distances |
| Core mutation rate | core_mu | $5x10^{-5}$ | Mutations per site per genome per generation |
| Homologous recombination rate (core genome) | HR_rate | 0.0 | Exchanges per core mutations |
| Horizontal gene transfer rate (accessory genome) | HGT_rate | 0.0 | Exchanges per core mutations |
| Basal gene turnover rate | rate_genes1 | 0.001 | Changes per site per genome per generation |
| Fast gene turnover rate | rate_genes2 | 10.0 | Changes per site per genome per generation |
| Proportion of fast genes | prop_genes2 | 0.0 | Proportion |
| Proportion of positively selected genes | prop_positive | NA (no selection) | Proportion |
| Positively-selected gene lambda | pos_lambda | NA (no selection) | N/A |
| Negatively-selected gene lambda | neg_lambda | NA (no selection) | N/A |
| Random number seed | seed | 0 | N/A |
| Genome size penalty | genome_size_penalty | 0.99 | N/A |
| Competition strength | competition_strength | 0.0 | N/A |

Supplementary Table 3: Results from MCMCglmm analysis of generalism score vs. PopPUNK-mod fitted parameters.

| | Coefficient | | | | | | Intercept | | | | | | $R^2$ | |
|---|---|---|---|---|---|---|---|---|---|---|---|---|---|---|
| **Parameter** | **posterior mean** | **lower 95% CI** | **upper 95% CI** | **effective sample size** | **p-value** | **Significance (p<0.05)** | **posterior mean** | **lower 95% CI** | **upper 95% CI** | **effective sample size** | **p-value** | **Significance (p<0.05)** | **Fixed effect** | **Random effect (phylogeny)** |
| **Proportion of fast genes** | 0.04454 | -0.27966 | 0.3346 | 5198 | 0.7817 | ns | 0.02649 | -0.09852 | 0.1492 | 5086 | 0.6817 | ns | 0.0007 | 0.6183 |
| **Basal gene turnover rate** | 0.328 | -3.63821 | 4.1382 | 4700 | 0.8774 | ns | 0.0387 | -0.07038 | 0.1378 | 4930 | 0.4634 | ns | 0.0002 | 0.6193 |
| **Core mutation rate** | 77.79862 | -402.59769 | 592.9574 | 4700 | 0.754 | ns | 0.02969 | -0.08466 | 0.1492 | 4700 | 0.6102 | ns | 0.0007 | 0.6191 |

Supplementary Table 4: Pangenome statistics for Pansim simulated populations. Core and rare thresholds defined at ≥ 95% and < 5% respectively.

| Changed parameter | Basal gene turnover rate | Proportion fast genes | HGT rate | HR rate | Competition strength | Proportion +ve selected genes | Gene +ve selection lambda | Gene -ve selection lambda | Core genome size | Intermediate genome size | Rare genome size | Average genome size | Stddev genome size |
|---|---|---|---|---|---|---|---|---|---|---|---|---|---|
| **Baseline** | 0.001 | 0 | 0 | 0 | 0 | NA | NA | NA | 1729 | 1716 | 1543 | 2522 | 10 |
| **Basal gene turnover rate** | 0.01 | 0 | 0 | 0 | 0 | NA | NA | NA | 1000 | 4000 | 0 | 2721 | 23 |
| | 0.1 | 0 | 0 | 0 | 0 | NA | NA | NA | 1000 | 4000 | 0 | 2954 | 33 |
| | 1 | 0 | 0 | 0 | 0 | NA | NA | NA | 1000 | 4000 | 0 | 2999 | 31 |
| **Proportion fast genes** | 0.001 | 0.1 | 0 | 0 | 0 | NA | NA | NA | 1679 | 1791 | 1520 | 2565 | 15 |
| | 0.001 | 0.5 | 0 | 0 | 0 | NA | NA | NA | 1375 | 2763 | 854 | 2738 | 24 |
| | 0.001 | 1 | 0 | 0 | 0 | NA | NA | NA | 1000 | 4000 | 0 | 2999 | 30 |
| **HGT rate** | 0.001 | 0 | 0.1 | 0 | 0 | NA | NA | NA | 1962 | 1257 | 1775 | 2551 | 12 |
| | 0.001 | 0 | 1 | 0 | 0 | NA | NA | NA | 2572 | 1078 | 1341 | 2860 | 13 |
| | 0.001 | 0 | 10 | 0 | 0 | NA | NA | NA | 4984 | 16 | 0 | 4966 | 6 |
| **HR rate** | 0.001 | 0 | 0 | 100 | 0 | NA | NA | NA | 1910 | 1218 | 1863 | 2505 | 12 |
| | 0.001 | 0 | 0 | 1000 | 0 | NA | NA | NA | 1889 | 1302 | 1797 | 2515 | 10 |
| **Competition strength** | 0.001 | 0 | 0 | 0 | 1 | NA | NA | NA | 1563 | 2073 | 1359 | 2526 | 12 |
| | 0.001 | 0 | 0 | 0 | 10 | NA | NA | NA | 1183 | 3244 | 572 | 2561 | 18 |
| **Proportion +ve selected genes** | 0.001 | 0 | 0 | 0 | 0 | 0 | 10 | 10 | 2323 | 158 | 2250 | 2412 | 7 |
| | 0.001 | 0 | 0 | 0 | 0 | 0.1 | 10 | 10 | 2374 | 170 | 2152 | 2460 | 6 |
| | 0.001 | 0 | 0 | 0 | 0 | 1 | 10 | 10 | 2700 | 331 | 1838 | 2820 | 6 |
| **Gene +ve selection lambda** | 0.001 | 0 | 0 | 0 | 0 | 0.5 | 0.1 | 100 | 2999 | 34 | 1350 | 3006 | 3 |
| | 0.001 | 0 | 0 | 0 | 0 | 0.5 | 1 | 100 | 2872 | 17 | 1583 | 2880 | 3 |
| | 0.001 | 0 | 0 | 0 | 0 | 0.5 | 10 | 100 | 2580 | 289 | 2041 | 2707 | 7 |
| **Gene -ve selection lambda** | 0.001 | 0 | 0 | 0 | 0 | 0.5 | 100 | 0.1 | 2294 | 38 | 1922 | 2320 | 3 |
| | 0.001 | 0 | 0 | 0 | 0 | 0.5 | 100 | 1 | 2349 | 94 | 1966 | 2400 | 3 |
| | 0.001 | 0 | 0 | 0 | 0 | 0.5 | 100 | 10 | 2370 | 228 | 2227 | 2496 | 8 |

Supplementary Table 5: Pangenome statistics for bacterial datasets

| **Species** | **Pangenome size (1%≤X≤100%)** | **Core genome size (X≥95%)** | **Intermediate genome size (5%≤X≤95%)** | **Rare genome size (1%≤X≤5%)** | **Average gene frequency** |
|---|---|---|---|---|---|
| *Acinetobacter baumannii* | 11195 | 2657 | 3724 | 4814 | 0.32260457 |
| *Bordetalla pertussis* | 6245 | 3138 | 1695 | 1412 | 0.594793208 |
| *Campylobacter jejuni* | 4727 | 1336 | 1470 | 1921 | 0.367283477 |
| *Enterococcus faecalis* | 7273 | 2148 | 2430 | 2695 | 0.380483305 |
| *Escherichia coli* | 17708 | 2828 | 6458 | 8422 | 0.262687147 |
| *Haemophilus influenzae* | 4386 | 1346 | 1454 | 1586 | 0.397435538 |
| *Helicobacter pylori* | 4560 | 1181 | 904 | 2475 | 0.318723569 |
| *Klebsiella pneumoniae* | 12062 | 4132 | 3345 | 4585 | 0.415136074 |
| *Listeria monocytogenes* | 5424 | 2488 | 1535 | 1401 | 0.532129994 |
| *Morexalla catarrhalis* | 3446 | 1427 | 1115 | 904 | 0.503010026 |
| *Mycobacterium abscessus* | 12488 | 4021 | 3478 | 4989 | 0.39502853 |
| *Mycobacterium tuberculosis* | 5668 | 3672 | 889 | 1107 | 0.705114214 |
| *Neisseria gonorrhoeae* | 3582 | 1780 | 860 | 942 | 0.590216657 |
| *Neisseria meningitidis* | 6856 | 1243 | 2529 | 3084 | 0.289385431 |
| *Pseudomonas aeruginosa* | 13498 | 5117 | 2819 | 5562 | 0.438012155 |
| *Salmonella enterica* | 10909 | 3553 | 2711 | 4645 | 0.401603371 |
| *Staphylococcus aureus* | 5326 | 2091 | 1761 | 1474 | 0.480415607 |
| *Stenotrophomonas maltophilia* | 13248 | 2996 | 4199 | 6053 | 0.316597172 |

| | | | | | |
|---|---|---|---|---|---|
| *Streptococcus agalactiae* | 4435 | 1572 | 1607 | 1256 | 0.454352142 |
| *Streptococcus dysgalactiae* subspecies *equisimilis* | 4681 | 1550 | 1666 | 1465 | 0.428283334 |
| *Streptococcus pneumoniae* | 6552 | 1425 | 2157 | 2970 | 0.309641112 |
| *Streptococcus pyogenes* | 3592 | 1393 | 1049 | 1150 | 0.464457857 |

### Supplementary Results

#### Effects of parameter alteration on pairwise distance distributions using Pansim

To understand how pangenome dynamics impact distributions of core and accessory distance distributions, we ran Pansim with a range of parameters corresponding to neutral (**Supplementary Figure 19**) or selection (**Supplementary Figure 20**) conditions. Increasing the basal gene turnover rate (the minimum rate at which genes are gained and lost, in changes per site per genome per generation) increased the rate at which accessory genome divergence saturation was reached relative to diversification of the core genome (**Supplementary Figure 19A**). Increasing the proportion of fast genes increased the y-intercept (**Supplementary Figure 19B**), emulating the observation in real data of rapid gene exchange before accruement of core genome diversity (**Supplementary Figure 2**). Furthermore, the highest parameter values in **Supplementary Figures 19A** and **19B** are equivalent, indicating that once the basal gene turnover rate is high enough, genes in this compartment cannot be distinguished from those in the fast turnover gene compartment. Increasing HGT rate in the accessory genome slightly increased the rate of accessory genome divergence saturation (**Supplementary Figure 19C**), similar to the effect observed in **Supplementary Figure 19A**, explained by genes gained in diverged lineages being shared. However, at 10 HGT exchanges per core SNP, the accessory genome was homogenised, resulting in no difference between individuals. Increasing HR rate had the opposite effect of increasing HGT rate, resulting in greater core genome variation at lower accessory genome distances (**Supplementary Figure 19D**), explained by core SNPs in diverged lineages being shared through horizontal transfer.

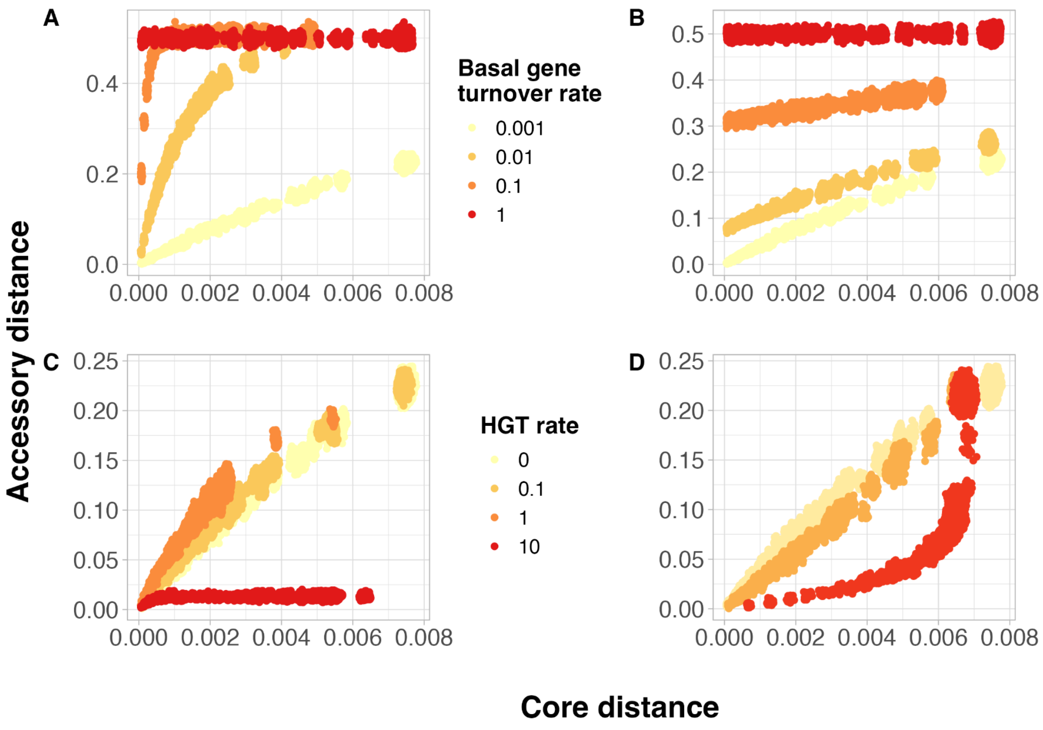


Supplementary Figure 19: Effect of changing neutral parameters on pairwise distance distributions in Pansim. (**A**) Changing basal gene turnover rate: the minimum rate at which genes are gained and lost, in changes per site per genome per generation. (**B**) Changing the proportion of fast genes in the pangenome, which have constant rate 10.0 changes per site per genome per generation, and so are saturated. (**C**) Changing HGT rate in accessory genome, in exchanges per core SNP (in per site per genome per generation). (**D**) Changing HR rate in core genome, in exchanges per core SNP (in per site per genome per generation). All simulations were run with a core genome mutation rate of $5x10^{-5}$ changes per site per genome per generation. Each point is a single pairwise comparison between two genomes. Values for all parameters not shown are available in **Supplementary Table 2**.

Alteration of selection parameters primarily affected the scale of core and accessory distance (**Supplementary Figure 20**). Notably, altering the strength of either positive or negative selection had visually identical impacts on the core vs accessory distance distributions, with increasing selection strength leading to an overall reduction in diversity due to favouring of specific highly fit genotypes. Increasing the proportion of positively selected genes gradually reduced the maximum core and accessory distance observed (**Supplementary Figure 20A**). Reducing the lambda of the double exponential distributions for either positively- (**Supplementary Figure 20B**) or negatively-selected (**Supplementary Figure 20C**) genes had near-identical impacts, reducing the maximum core and accessory distance observed. There was an opposite impact on core genome and average genome size when changing the lambda for either positively- or negative-selected genes, with lambda reduction favouring an increase in both statistics for the positively-selected genes, but a decrease in both statistics for negatively-selected genes (**Supplementary Table 4**). Increasing competition strength had the opposite effect, favouring more diverse genotypes (**Supplementary Figure 21**), further evidenced by an increase in intermediate genome size (**Supplementary Table 4**). Overall, Pansim is a flexible simulator that translates changes in pangenome dynamics parameters to pairwise distance and gene frequency distributions.

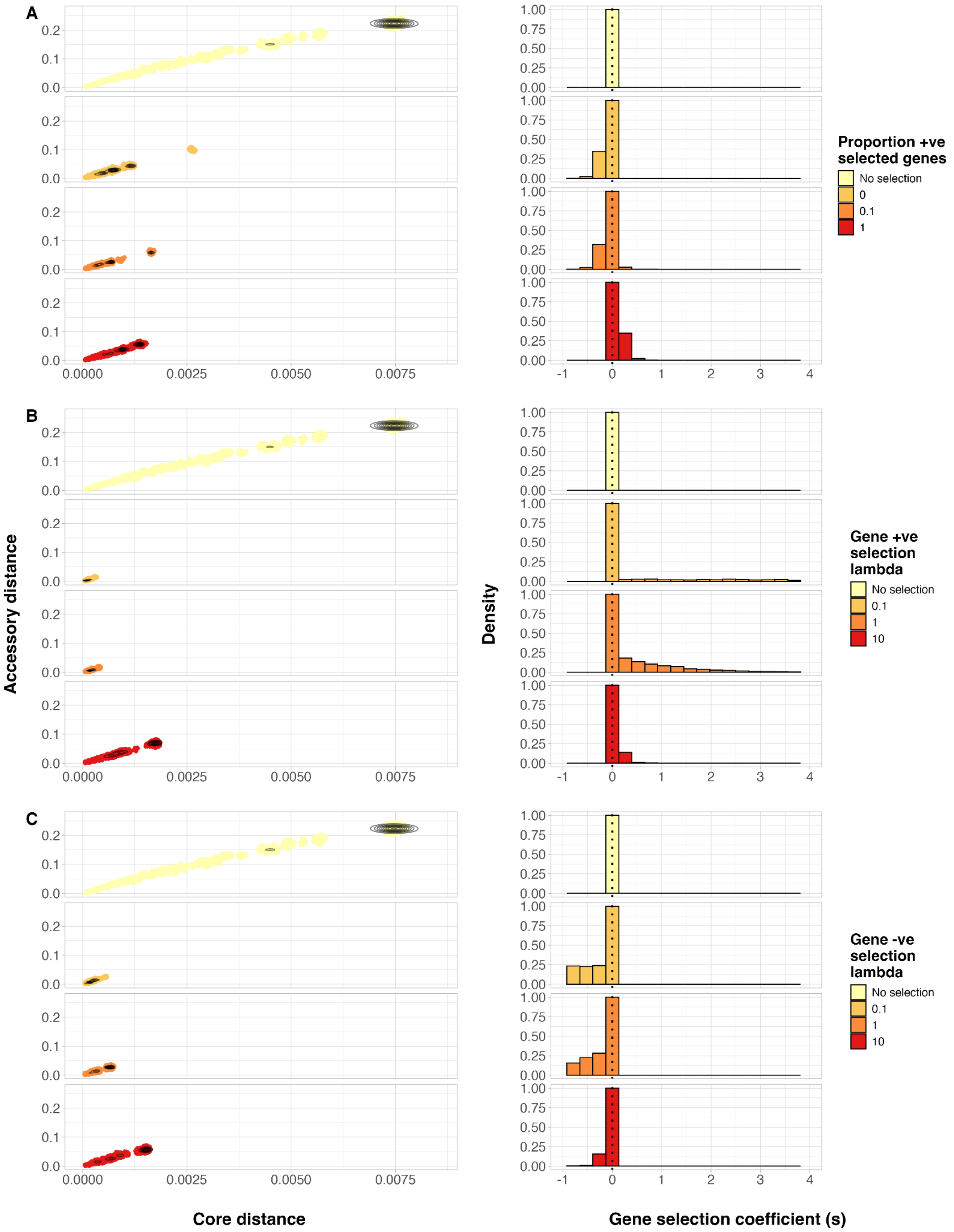


Supplementary Figure 20: Effect of changing selection parameters on pairwise distance distributions in Pansim. (**A**) Changing the proportion of positively selected genes in the pangenome. (**B**) Changing the lambda of the exponential distribution for the positively selected genes (larger lambda favours smaller absolute *s*). (**C**) Changing the lambda of the exponential distribution for the negatively selected genes (larger lambda favours smaller absolute *s*). Left column: pairwise distance distributions, with each point representing a single pairwise comparison between genomes, and contours indicating point density. Right column: selection coefficient distributions for pairwise distance distributions in the left column. Contours indicate respective point density. Each point is a single pairwise comparison between two genomes. Values for all parameters not shown are available in **Supplementary Table 2**.

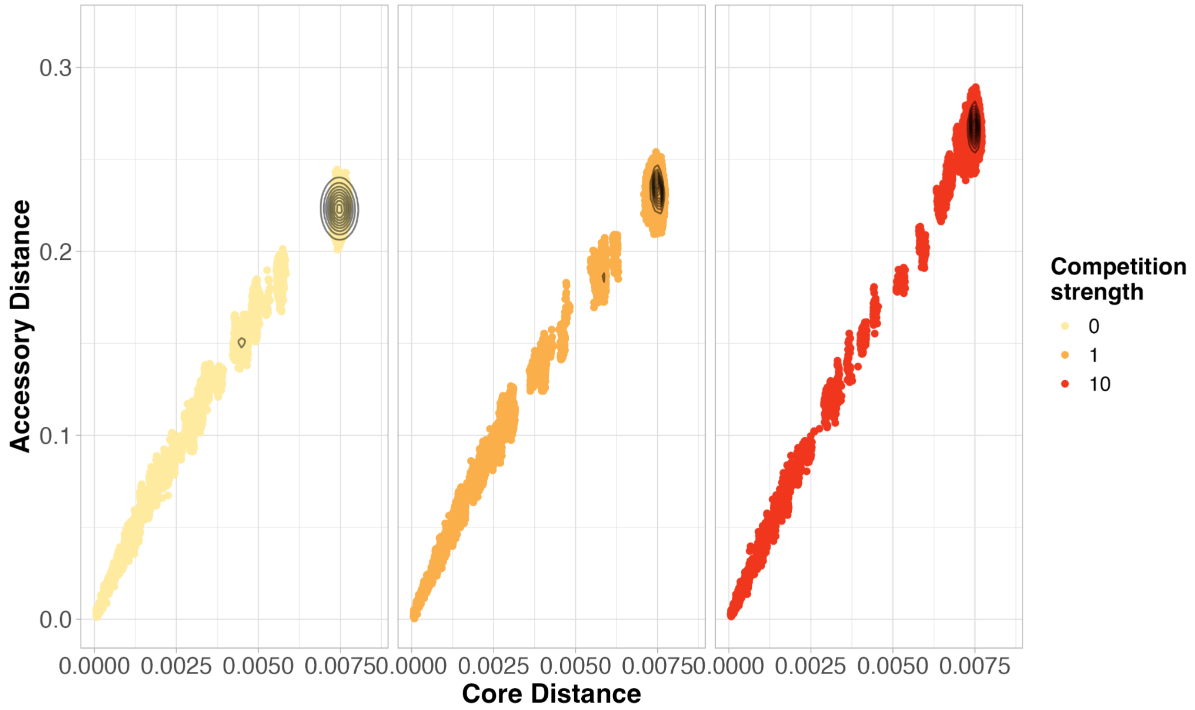


Supplementary Figure 21: Effect of increasing competition strength on pairwise distance distributions in Pansim. Each point represents a single pairwise comparison between genomes. Contours represent point density.

#### Testing sensitivity of population diversity to fixed population parameters

To determine the sensitivity of population diversity to fixed population parameters, we compared the average and standard deviation of core and accessory pairwise distance when varying the number of generations and population size. After 100 generations, both core and accessory distances reach a steady state (**Supplementary Figure 22**), the onset of which was not affected by population size. However, when altering fitted parameters, variation in the pairwise distances increases, with simulations with 500 generations or more showing similar average trajectories, independent of population size (**Supplementary Figure 23**). Therefore, pairwise distance measures are robust to variation and therefore misspecification in fixed population parameters when simulating with 500 generations or more.

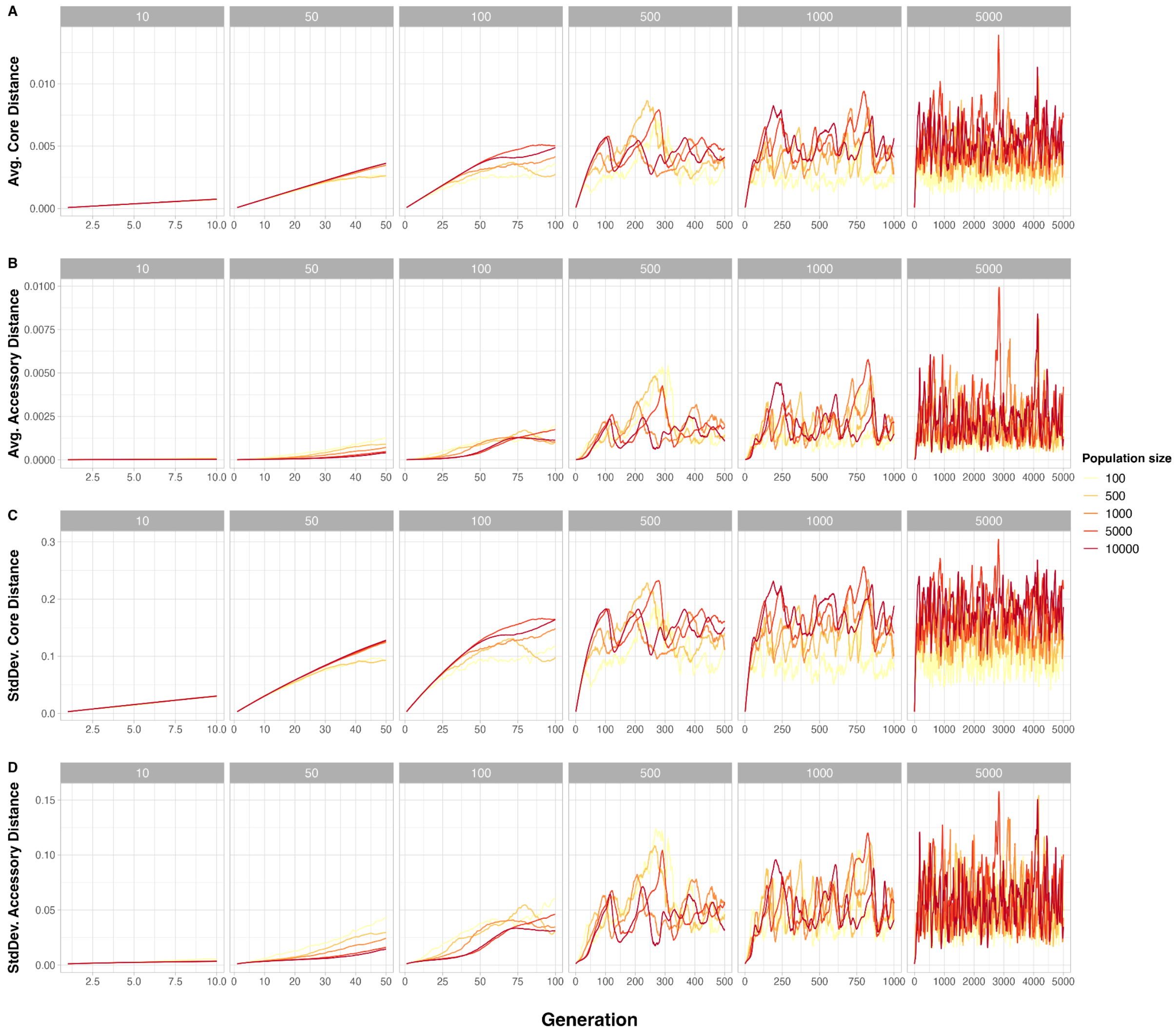


Supplementary Figure 22: Effect of number of generations and population size on population diversity in Pansim without varying fitted parameters. (**A**) Average core distance, (**B**) average accessory distance, (**C**) standard deviation of core distances and (**D**) standard deviation of accessory distances. Facets describe the total number of generations used in the simulation, colours describe the number of individuals in the population. Simulations were run with parameters described in **Supplementary Table 2** with the exception of population size and number of generations.

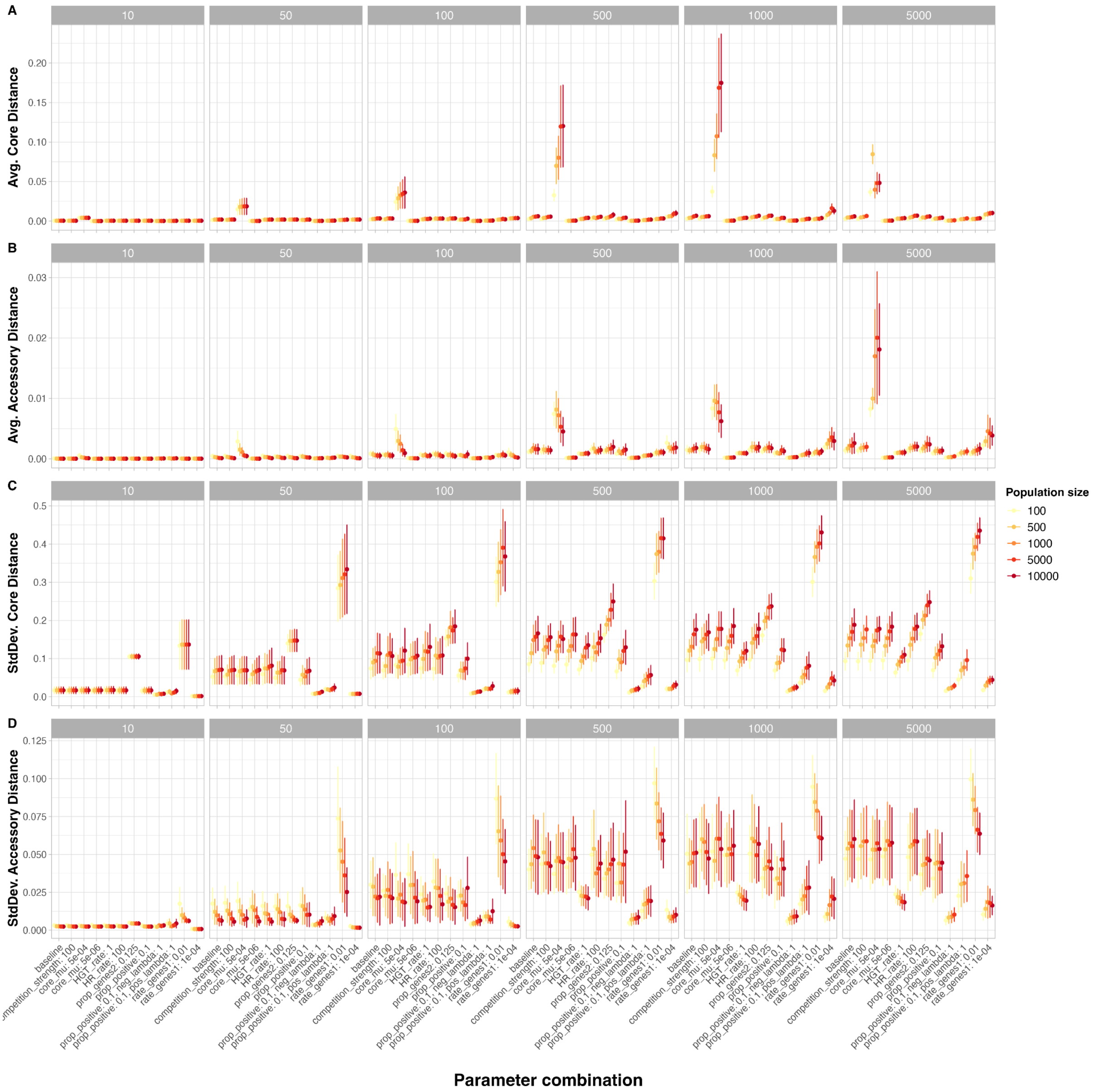


Supplementary Figure 23: Effect of number of generations and population size on population diversity in Pansim with variation when varying fitted parameters. (**A**) Average core distance, (**B**) average accessory distance, (**C**) standard deviation of core distances and (**D**) standard deviation of accessory distances. Points show the average value across all generations, error bars show the standard deviation of the value across all generations. Facets describe the total number of generations used in the simulation, colours describe the number of individuals in the population. Simulations were run with parameters described in **Supplementary Table 2** with the exception of population size and number of generations for the baseline approach, with altered fitted parameters shown on the x axis label.

### Testing ground truth parameter recapitulation with Approximate Bayesian Computation

To ensure reliable fitting, we first tested the ability of PopPUNK-mod to recapitulate known parameters by fitting to simulated data generated by Pansim. As a benchmark of the baseline performance of PopPUNK-mod, the core mutation rate and basal gene turnover rate in the accessory genome were jointly fitted (**Supplementary Figure 24**). PopPUNK-mod was able to recapitulate basal gene turnover rate parameters across 4 orders of magnitude ($10^{-4}$-$10^{-1}$ changes per site per genome per generation), with the ground-truth values sitting within the 95% credible intervals, although it was unable to recapitulate the lowest parameter value tested ($10^{-5}$ changes per site per genome per generation). For the core mutation rate, PopPUNK-mod was also able to recapitulate parameter values across 4 orders of magnitude ($10^{-7}$-$10^{-4}$ changes per site per genome per generation). PopPUNK-mod was not able to recapitulate values for $10^{-6}$ and $10^{-3}$ changes per site per genome per generation, although median parameter estimates were within one order of magnitude of ground-truth values. These results highlight that PopPUNK-mod is able to accurately detect changes in core and accessory genome mutation rate over a wide range of parameter values.

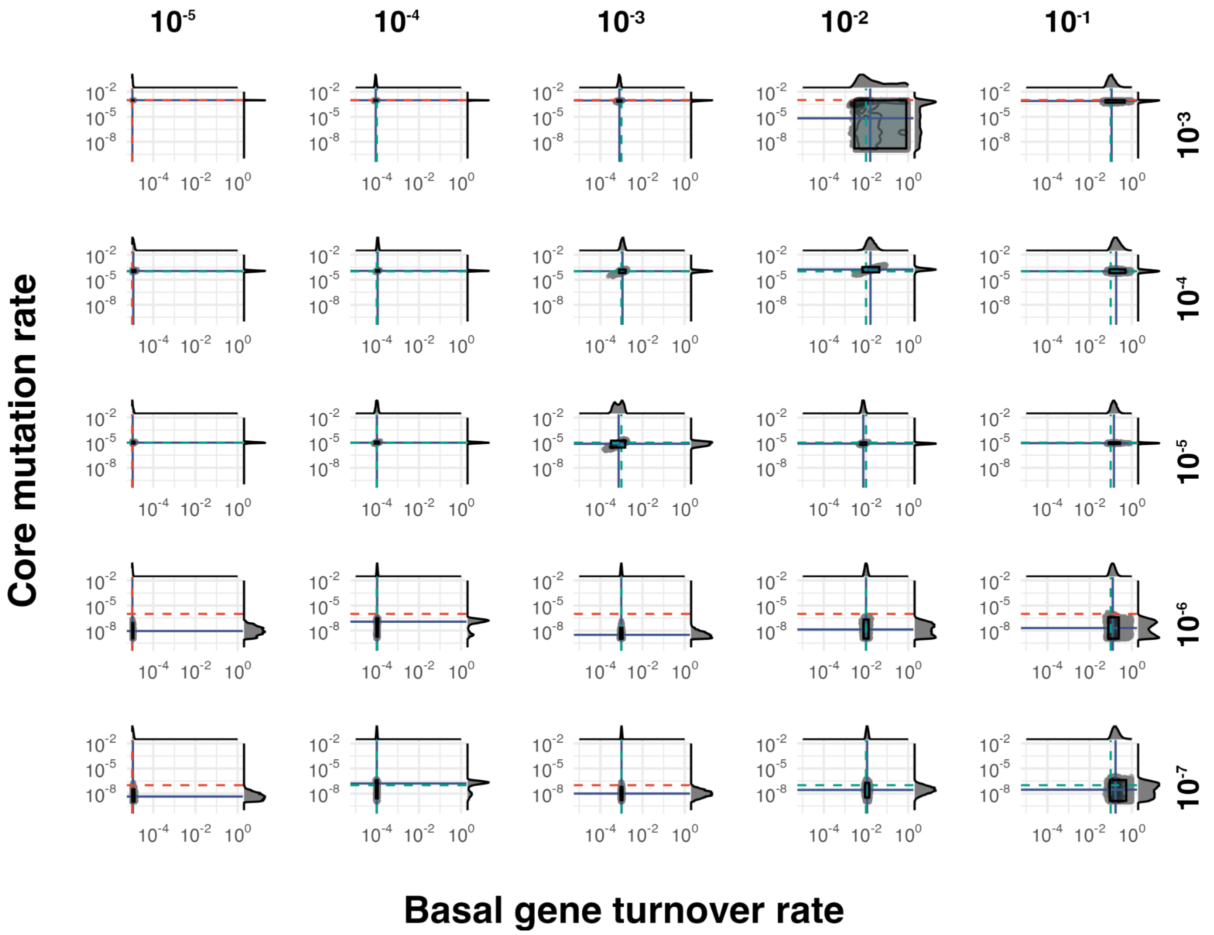


Supplementary Figure 24: Benchmarking of ABC fitting to ground-truth simulations with varying core and accessory mutation rates. Each panel represents a different simulation with specific core mutation rate (rows, units in mutations per site per genome per generation) or basal gene turnover rate (columns, units in mutations per site per genome per generation). The ground truth values for each parameter for the simulation is denoted above and to the right of each column or row for core mutation rate and basal gene turnover rate respectively. Each point represents a single MCMC sample, with contours overlaid to represent density. Marginal distributions on the top and right of each panel represent point density. Blue solid lines represent the median values for the MCMC sampled parameter values. Blue boxes represent the 95% credible intervals for both parameters. Dashed lines represent the ground-truth values for each simulation; green indicates this value sits inside the 95% credible intervals of the predicted parameter values, red if outside. Both axes are on a logarithmic scale.

We then tested the ability of PopPUNK-mod to recapitulate additional pangenome dynamics parameters from Pansim runs (**Supplementary Figure 25**). For neutral parameters (**Supplementary Figure 25 A-C**), PopPUNK-mod was able to recapitulate accessory and core mutation rates, as well as the additional parameter included. The model was able to accurately capture the proportion of fast genes in the simulation, performing best of all tested parameters in terms of the size of the credible intervals and capturing of the ground truth parameter in the credible intervals. Credible intervals HGT and HR were notably large, spanning 6 and 3 orders of magnitude respectively. For selection parameters (**Supplementary Figure 25 D-G**), PopPUNK-mod was able to recapitulate parameters for the proportion of positively selected genes and the positively-selected gene lambda, whilst ground-truth values for the negatively-selected gene lambda and competition strength fell outside of the predicted parameters 95% credible intervals. Ground-truth values of basal gene turnover rate and core genome mutation rate also fell outside of the credible intervals for the proportion of positively selected genes and the negatively-selected gene lambda. These results indicate that prediction of selection parameters using PopPUNK-mod, while possible, is difficult likely due to the similar effects on pairwise distance distributions caused by positive and negative selection (**Supplementary Figure 20**).

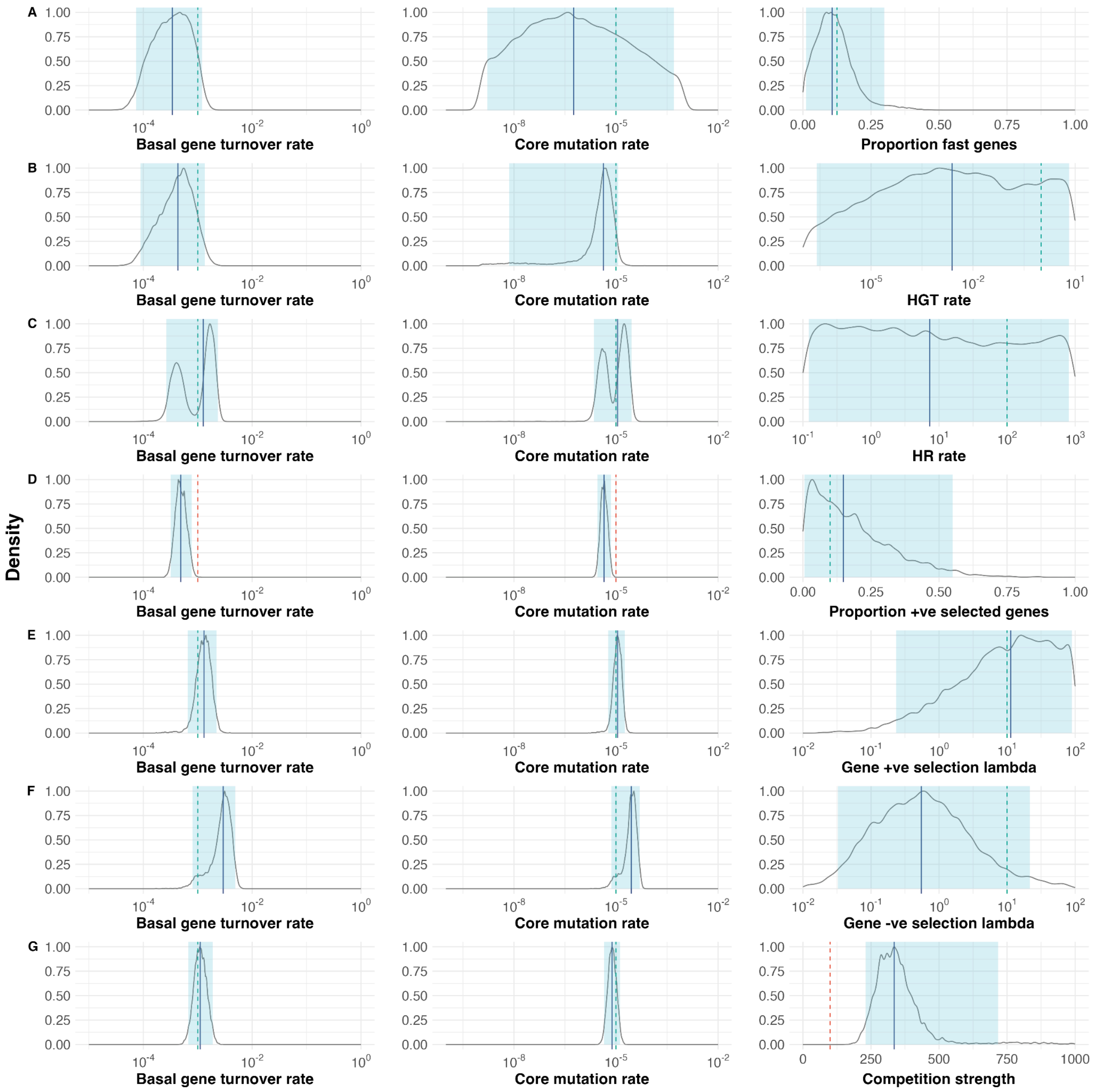


Supplementary Figure 25: PopPUNK-mod fitting with additional parameters. Rows show results from individual simulations fitting basal gene turnover rate (left column) and core mutation rate (middle column) and a third parameter (right column): (**A**) proportion of fast genes, (**B**) HGT rate, (**C**) HR rate, (**D**) proportion of positively selected genes, (**E**) positively-selected gene lambda, (**E**) negatively-selected gene lambda, (**G**) competition strength. Density plots show MCMC samples for each fitted parameter. Regions inside blue shading represent 95% credible intervals, the blue solid bar represents the median parameter estimate. Dashed lines represent the ground-truth values for each simulation; green indicates this value sits inside the 95% credible intervals of the predicted parameter values, red if outside. Axis for basal gene turnover rate and core mutation rate are on a logarithmic scale. Scales vary between logarithmic and non-logarithmic for parameter shown in the right column.

To determine the sensitivity of PopPUNK-mod estimates to fixed population parameters, we conducted a sensitivity analysis of parameter estimate accuracy, fitting PopPUNK-mod to Pansim simulations where population parameters were mismatched between the ground-truth simulations and those provided to PopPUNK-mod (**Supplementary Figure 13**). Results highlighted that sensitivity to fixed parameters varies by the fitted parameter chosen. Lambdas for positive and negative gene selection appear to be insensitive to fixed parameter perturbation, however, as in reality these parameters must be fitted jointly with the proportion of +ve selection genes, these parameters cannot be fitted robustly.

To determine whether accuracy of parameter inference could be improved if parameter numbers were reduced, we removed inference of core mutation rate, and added fitting of selection parameters (**Supplementary Figure 26**) and competition (**Supplementary Figure 27 & 28**). Selection parameters were still non-identifiable with large credible intervals, even when reducing the number of parameters. However, fits of competition strength were accurate even when including the proportion of fast genes as an additional parameter, including the parameter in a small credible interval. Reducing the competition strength parameter prior range from 0-1000 (**Supplementary Figure 27**) to 0-100 (**Supplementary Figure 28**) did not visibly impact results. However, when fitting competition strength to observed data, fixing core mutation rate at the median shown in **Figure 4**, we observed inconsistent parameter estimates for basal gene turnover rate and the proportion of fast genes, as well as competition strength credible intervals which spanned a majority of prior parameter range for many species (**Supplementary Figure 29**). Therefore, these parameterisations were deemed non-identifiable.

Overall, we conclude that the three most robust parameters to changes in all fixed parameters are basal gene turnover rate, core mutation rate and proportion of fast genes. Unfortunately, selection parameters are not identifiable when using the current summary statistics.

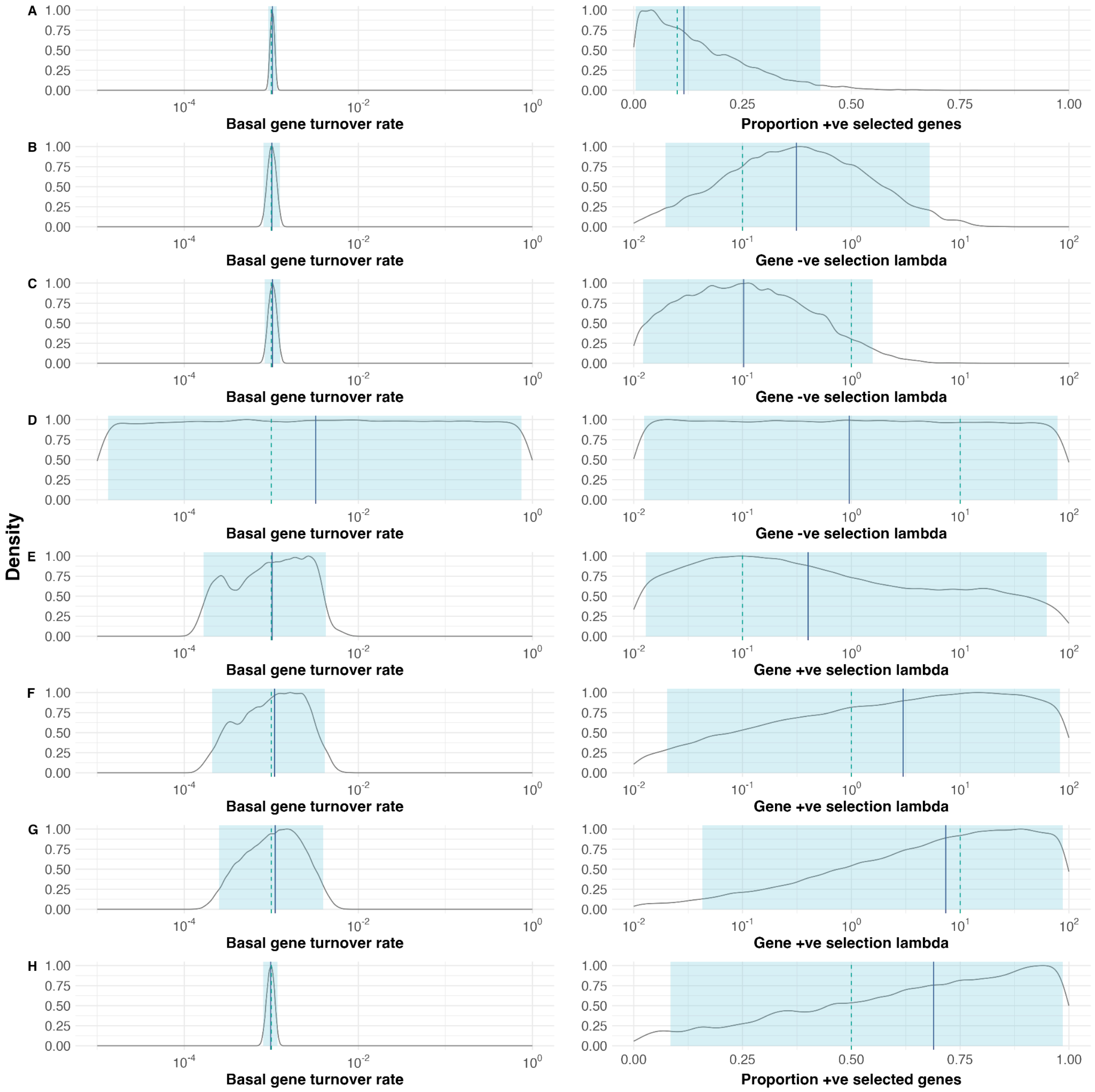


Supplementary Figure 26: PopPUNK-mod performance on simulated runs fitting only basal gene turnover rate and single selection parameter. (**A**) Proportion of positively selected genes = 0.1, negative lambda = 10.0, positive lambda = 10.0, fitting proportion of positively selected genes. (**B**) Proportion of positively selected genes = 0.1, negative lambda = 0.1, positive lambda = 10.0, fitting negative lambda. (**C**) Proportion of positively selected genes = 0.1, negative lambda = 1.0, positive lambda = 10.0, fitting negative lambda. (**D**) Proportion of positively selected genes = 0.1, negative lambda = 10.0, positive lambda = 10.0, fitting negative lambda. (**E**) Proportion of positively selected genes = 0.1, negative lambda = 10.0, positive lambda = 0.1, fitting positive lambda. (**F**) Proportion of positively selected genes = 0.1, negative lambda = 10.0, positive lambda = 1.0, fitting positive lambda. (**G**) Proportion of positively selected genes = 0.1, negative lambda = 10.0, positive lambda = 10.0, fitting positive lambda. (**H**) Proportion of positively selected genes = 0.5, negative lambda = 10.0, positive lambda = 10.0, fitting proportion of positively selected genes. Basal gene turnover rate was 0.001 gene exchange per site per individual per generation for all simulations. The dotted line denoted the real parameter value (red if outside 95% credible intervals, green if inside); the solid blue line is the posterior distribution median, the blue shaded area denotes the central 95% credible interval.

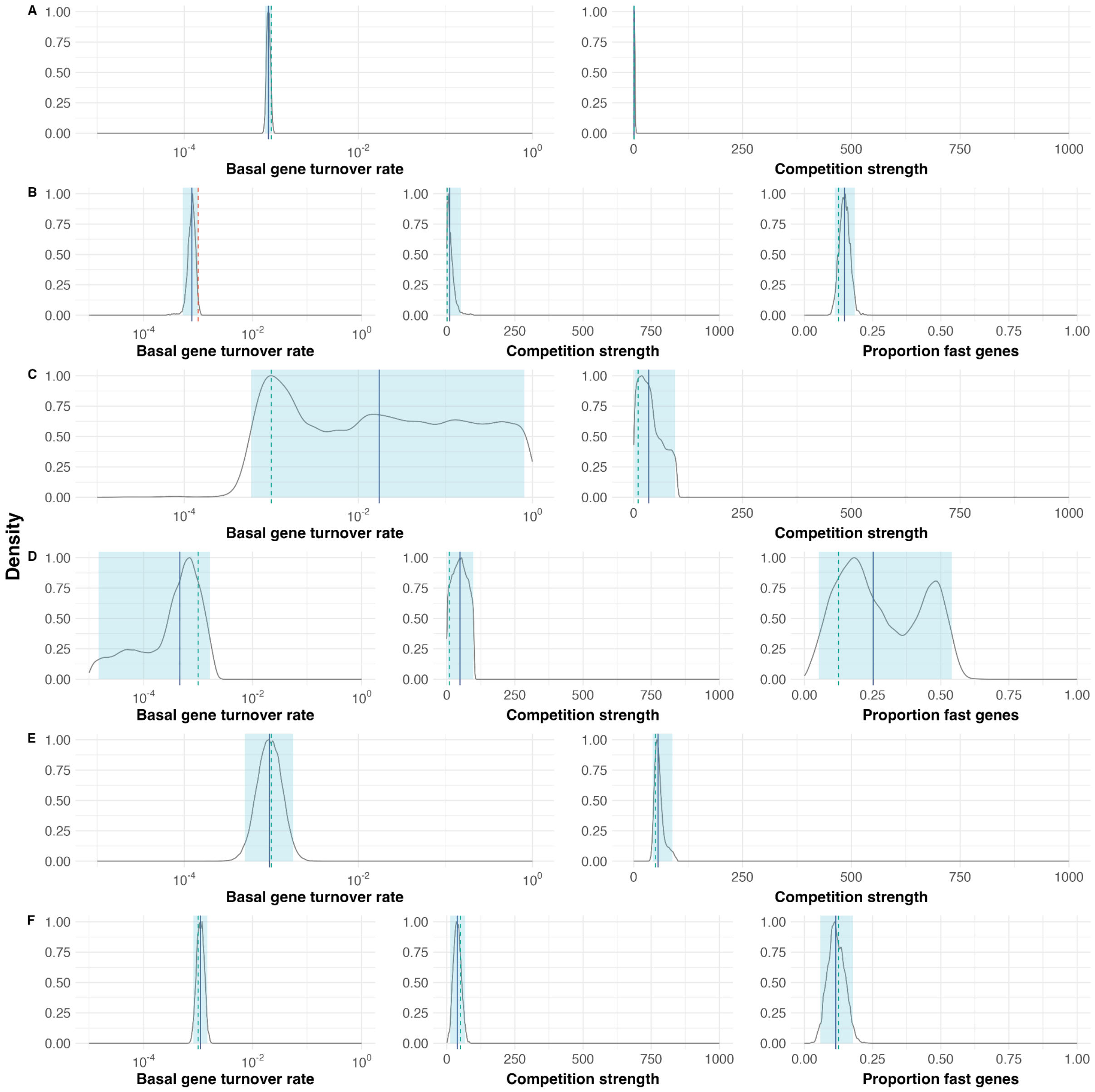


Supplementary Figure 27: PopPUNK-mod performance on simulated runs fitting only basal gene turnover rate with competition strength and proportion of fast genes, with uniform competition strength prior between 0-1000. (**A**) Competition strength = 1, fitting competition strength. (**B**) Competition strength = 1, proportion of fast genes = 0.125, fitting competition strength. (**C**) Competition strength = 10, fitting competition strength. (**D**) Competition strength = 10, proportion of fast genes = 0.125, fitting competition strength. (**E**) Competition strength = 50, fitting competition strength. (**F**) Competition strength = 50, proportion of fast genes = 0.125, fitting competition strength. Basal gene turnover rate was 0.001 gene exchange per site per individual per generation for all simulations. The dotted line denoted the real parameter value (red if outside 95% credible intervals, green if inside); the solid blue line is the posterior distribution median, the blue shaded area denotes the central 95% credible interval.

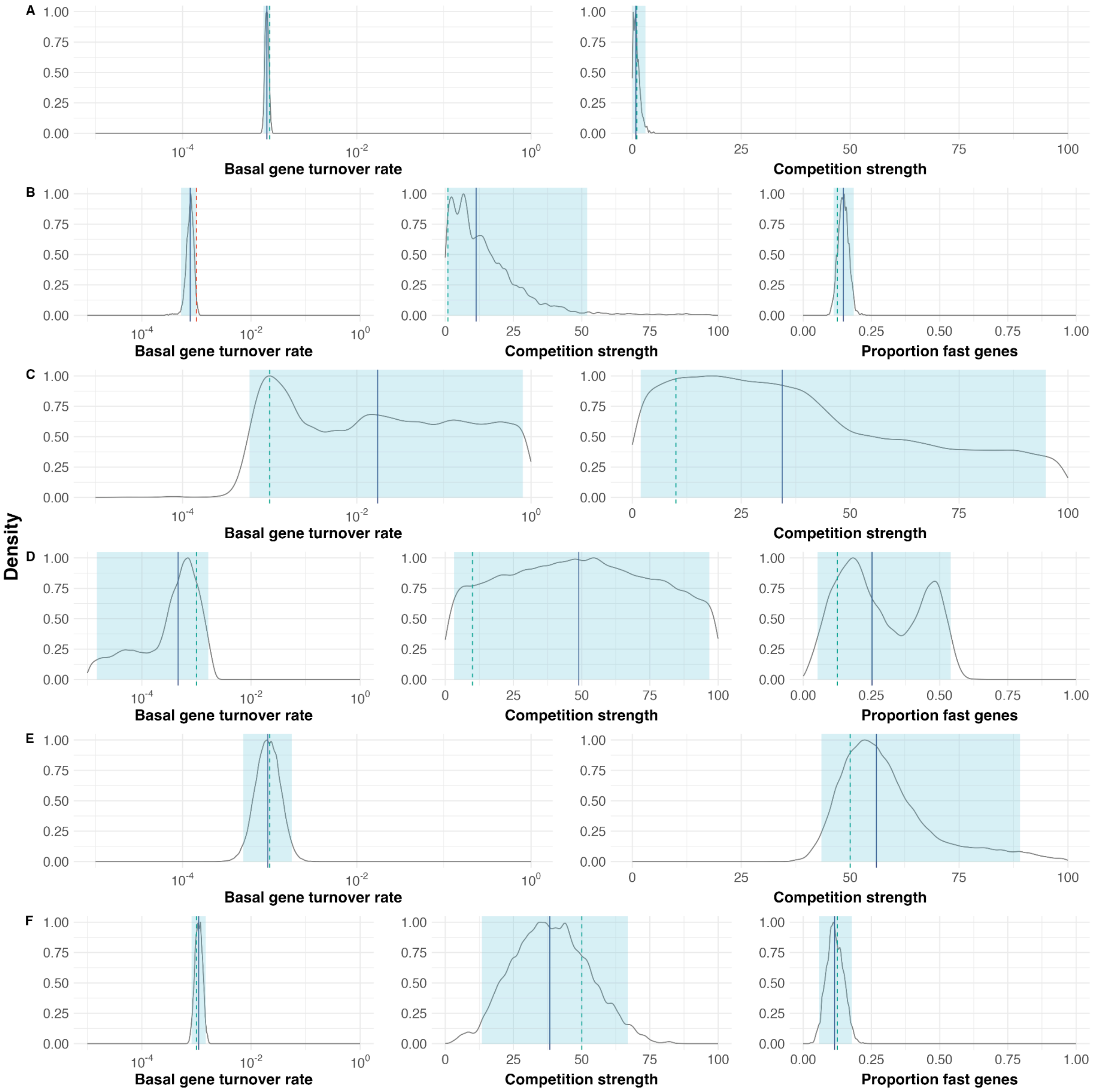


Supplementary Figure 28: PopPUNK-mod performance on simulated runs fitting only basal gene turnover rate with competition strength and proportion of fast genes, with uniform competition strength prior between 0-100. (**A**) Competition strength = 1, fitting competition strength. (**B**) Competition strength = 1, proportion of fast genes = 0.125, fitting competition strength. (**C**) Competition strength = 10, fitting competition strength. (**D**) Competition strength = 10, proportion of fast genes = 0.125, fitting competition strength. (**E**) Competition strength = 50, fitting competition strength. (**F**) Competition strength = 50, proportion of fast genes = 0.125, fitting competition strength. Basal gene turnover rate was 0.001 gene exchange per site per individual per generation for all simulations. The dotted line denoted the real parameter value (red if outside 95% credible intervals, green if inside); the solid blue line is the posterior distribution median, the blue shaded area denotes the central 95% credible interval.

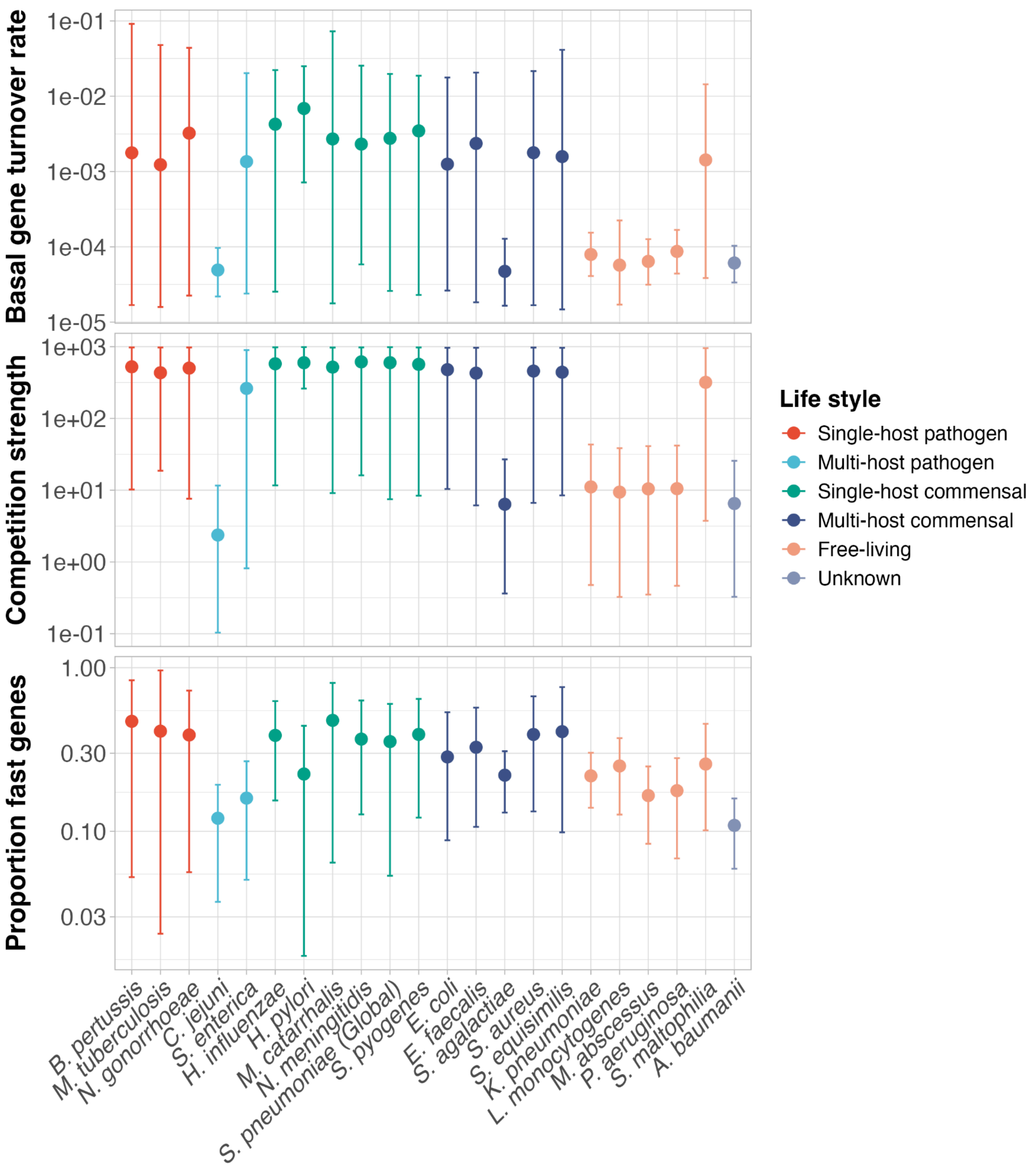


Supplementary Figure 29: PopPUNK-mod estimated parameter posterior distributions of observed bacterial genomic data. Points highlight the median of posterior distribution, error bars indicate the 95% credible intervals centred on the median. Basal gene turnover rate is measured in mutations per site per genome per generation. Competition strength is a multiplier to the average pairwise distance of an individual to all other individuals in a population, which is taken as the individual's overall fitness. Proportion of fast genes is measured as the proportion of total accessory genes that are exchanged instantaneously in each generation. All simulations were performed with 1000 individuals and 500 generations. Points are coloured by species lifestyle assignments. Data for each species is available in **Supplementary File 5**.

Brooks, Lavida R. K., and George I. Mias. 2018. “Streptococcus Pneumoniae’s Virulence and Host Immunity: Aging, Diagnostics, and Prevention.” *Frontiers in Immunology* 9 (June): 1366.

Chewapreecha, Claire, Simon R. Harris, Nicholas J. Croucher, et al. 2014. “Dense Genomic Sampling Identifies Highways of Pneumococcal Recombination.” *Nature Genetics* 46 (3): 305–309.

Da Cunha, Violette, Mark R. Davies, Pierre-Emmanuel Douarre, et al. 2014. “Streptococcus Agalactiae Clones Infecting Humans Were Selected and Fixed through the Extensive Use of Tetracycline.” *Nature Communications* 5 (1): 4544.

David, Sophia, Sandra Reuter, Simon R. Harris, et al. 2019. “Epidemic of Carbapenem-Resistant Klebsiella Pneumoniae in Europe Is Driven by Nosocomial Spread.” *Nature Microbiology* 4 (11): 1919–1929.

Davies, Mark R., Liam McIntyre, Ankur Mutreja, et al. 2019. “Atlas of Group A Streptococcal Vaccine Candidates Compiled Using Large-Scale Comparative Genomics.” *Nature Genetics* 51 (6): 1035–1043.

Denet, Elodie, Valentin Vasselon, Béatrice Burdin, Sylvie Nazaret, and Sabine Favre-Bonté. 2018. “Survival and Growth of Stenotrophomonas Maltophilia in Free-Living Amoebae (FLA) and Bacterial Virulence Properties.” *PloS One* 13 (2): e0192308.

Erwin, Alice L., and Arnold L. Smith. 2007. “Nontypeable Haemophilus Influenzae: Understanding Virulence and Commensal Behavior.” *Trends in Microbiology* 15 (8): 355–362.

Folkesson, Anders, Lars Jelsbak, Lei Yang, et al. 2012. “Adaptation of Pseudomonas Aeruginosa to the Cystic Fibrosis Airway: An Evolutionary Perspective.” *Nature Reviews. Microbiology* 10 (12): 841–851.

Gladstone, Rebecca A., Stephanie W. Lo, John A. Lees, et al. 2019. “International Genomic Definition of Pneumococcal Lineages, to Contextualise Disease, Antibiotic Resistance and Vaccine Impact.” *EBioMedicine* 43: 338–346.

Gröschel, Matthias I., Conor J. Meehan, Ivan Barilar, et al. 2020. “The Phylogenetic Landscape and Nosocomial Spread of the Multidrug-Resistant Opportunist Stenotrophomonas Maltophilia.”

*Nature Communications* 11 (1): 1–12.

Haag, Andreas F., J. Ross Fitzgerald, and José R. Penadés. 2019. “Staphylococcus Aureus in Animals.” *Microbiology Spectrum* 7 (3). https://doi.org/10.1128/microbiolspec.GPP3-0060-2019.

Haidan, A., S. R. Talay, M. Rohde, K. S. Sriprakash, B. J. Currie, and G. S. Chhatwal. 2000. “Pharyngeal Carriage of Group C and Group G Streptococci and Acute Rheumatic Fever in an Aboriginal Population.” *Lancet* 356 (9236): 1167–1169.

Hammerum, A. M. 2012. “Enterococci of Animal Origin and Their Significance for Public Health.” *Clinical Microbiology and Infection: The Official Publication of the European Society of Clinical Microbiology and Infectious Diseases* 18 (7): 619–625.

Horesh, Gal, Grace A. Blackwell, Gerry Tonkin-Hill, Jukka Corander, Eva Heinz, and Nicholas R. Thomson. 2021. “A Comprehensive and High-Quality Collection of Escherichia Coli Genomes and Their Genes.” *Microbial Genomics* 7 (2). https://doi.org/10.1099/mgen.0.000499.

Humphrey, Suzanne, Gemma Chaloner, Kirsty Kemmett, et al. 2014. “Campylobacter Jejuni Is Not Merely a Commensal in Commercial Broiler Chickens and Affects Bird Welfare.” *mBio* 5 (4): e01364–14.

Hunt, Martin, Leandro Lima, Wei Shen, John Lees, and Zamin Iqbal. 2024. “AllTheBacteria - All Bacterial Genomes Assembled, Available and Searchable.” *bioRxiv*, 2024.03.08.584059–2024.03.08.584059.

Jolley, Keith A., James E. Bray, and Martin C. J. Maiden. 2018. “Open-Access Bacterial Population Genomics: BIGSdb Software, the PubMLST.org Website and Their Applications.” *Wellcome Open Research* 3. https://doi.org/10.12688/wellcomeopenres.14826.1.

Karalus, R., and A. Campagnari. 2000. “Moraxella Catarrhalis: A Review of an Important Human Mucosal Pathogen.” *Microbes and Infection* 2 (5): 547–559.

Krismer, Bernhard, Christopher Weidenmaier, Alexander Zipperer, and Andreas Peschel. 2017. “The Commensal Lifestyle of Staphylococcus Aureus and Its Interactions with the Nasal Microbiota.” *Nature Reviews. Microbiology* 15 (11): 675–687.

Ladhani, Shamez N., Jay Lucidarme, Sydel R. Parikh, Helen Campbell, Ray Borrow, and Mary E. Ramsay. 2020. “Meningococcal Disease and Sexual Transmission: Urogenital and Anorectal Infections and Invasive Disease due to Neisseria Meningitidis.” *Lancet* 395 (10240): 1865–1877.

Lagune, Marion, Laurent Kremer, and Jean-Louis Herrmann. 2024. “Mycobacterium Abscessus, a Complex of Three Fast-Growing Subspecies Sharing Virulence Traits with Slow-Growing Mycobacteria.” *Clinical Microbiology and Infection: The Official Publication of the European Society of Clinical Microbiology and Infectious Diseases* 30 (6): 726–731.

Lenz, Jonathan D., and Joseph P. Dillard. 2018. “Pathogenesis of Neisseria Gonorrhoeae and the Host Defense in Ascending Infections of Human Fallopian Tube.” *Frontiers in Immunology* 9 (November): 2710.

Mikucki, August, Nicolie R. McCluskey, and Charlene M. Kahler. 2022. “The Host-Pathogen Interactions and Epicellular Lifestyle of Neisseria Meningitidis.” *Frontiers in Cellular and Infection Microbiology* 12. https://doi.org/10.3389/FCIMB.2022.862935.

Mladenova-Hristova, Irena, Olga Grekova, and Ami Patel. 2017. “Zoonotic Potential of Helicobacter Spp.” *Wei Mian Yu Gan Ran Za Zhi [Journal of Microbiology, Immunology, and Infection]* 50 (3): 265–269.

Mohammadnabi, Noura, Jebreil Shamseddin, Mobina Emadi, et al. 2024. “Mycobacterium Tuberculosis: The Mechanism of Pathogenicity, Immune Responses, and Diagnostic Challenges.” *Journal of Clinical Laboratory Analysis* 38 (23): e25122.

Nowak, Maciej, Zbigniew Paluszak, Natalia Wiktorczyk-Kapischke, et al. 2024. “Characterization of

Listeria Monocytogenes Strains Isolated from Soil under Organic Carrot Farming." *Frontiers in Microbiology* 15: 1530446.

Petit, Robert A., and Timothy D. Read. 2018. "Staphylococcus Aureus Viewed from the Perspective of 40,000+ Genomes." *PeerJ* 6 (7). https://doi.org/10.7717/PEERJ.5261.

Pöntinen, Anna K., Janetta Top, Sergio Arredondo-Alonso, et al. 2021. "Apparent Nosocomial Adaptation of Enterococcus Faecalis Predates the Modern Hospital Era." *Nature Communications* 12 (1): 1–13.

Porcellato, Davide, Marit Smistad, Siv Borghild Skeie, Hannah Joan Jørgensen, Lars Austbø, and Oddvar Oppegaard. 2021. "Whole Genome Sequencing Reveals Possible Host Species Adaptation of Streptococcus Dysgalactiae." *Scientific Reports* 11 (1). https://doi.org/10.1038/S41598-021-96710-Z.

Preston, Andrew. 2005. "Bordetella Pertussis: The Intersection of Genomics and Pathobiology." *Journal de l'Association Medicale Canadienne [Canadian Medical Association Journal]* 173 (1): 55–62.

Ren, Yunxiao, Carmen Li, Dulmini Nanayakkara Sapugahawatte, et al. 2024. "Predicting Hosts and Cross-Species Transmission of Streptococcus Agalactiae by Interpretable Machine Learning." *Computers in Biology and Medicine* 171 (108185): 108185.

Reshetnyak, Vasiliy Ivanovich, Alexandr Igorevich Burmistrov, and Igor Veniaminovich Maev. 2021. "Helicobacter Pylori: Commensal, Symbiont or Pathogen?" *World Journal of Gastroenterology* 27 (7): 545–560.

Rocha, Jaqueline, Isabel Henriques, Margarita Gomila, and Célia M. Manaia. 2022. "Common and Distinctive Genomic Features of Klebsiella Pneumoniae Thriving in the Natural Environment or in Clinical Settings." *Scientific Reports* 12 (1): 10441–10441.

Ruis, Christopher, Josephine M. Bryant, Scott C. Bell, et al. 2021. "Dissemination of Mycobacterium Abscessus via Global Transmission Networks." *Nature Microbiology* 6 (10): 1279–1288.

Sayers, Eric W., Mark Cavanaugh, Karen Clark, et al. 2021. "GenBank." *Nucleic Acids Research* 49 (D1): D92–D96.

Tenaillon, Olivier, David Skurnik, Bertrand Picard, and Erick Denamur. 2010. "The Population Genetics of Commensal Escherichia Coli." *Nature Reviews. Microbiology* 8 (3): 207–217.

Timme, Ruth E., Maria Sanchez Leon, and Marc W. Allard. 2019. "Utilizing the Public GenomeTrakr Database for Foodborne Pathogen Traceback." *Methods in Molecular Biology* 1918: 201–212.

Winfield, Mollie D., and Eduardo A. Groisman. 2003. "Role of Nonhost Environments in the Lifestyles of Salmonella and Escherichia Coli." *Applied and Environmental Microbiology* 69 (7): 3687–3694.
