## Supplementary File 3 for "Rapid gene exchange explains differences in bacterial pangenome structure"

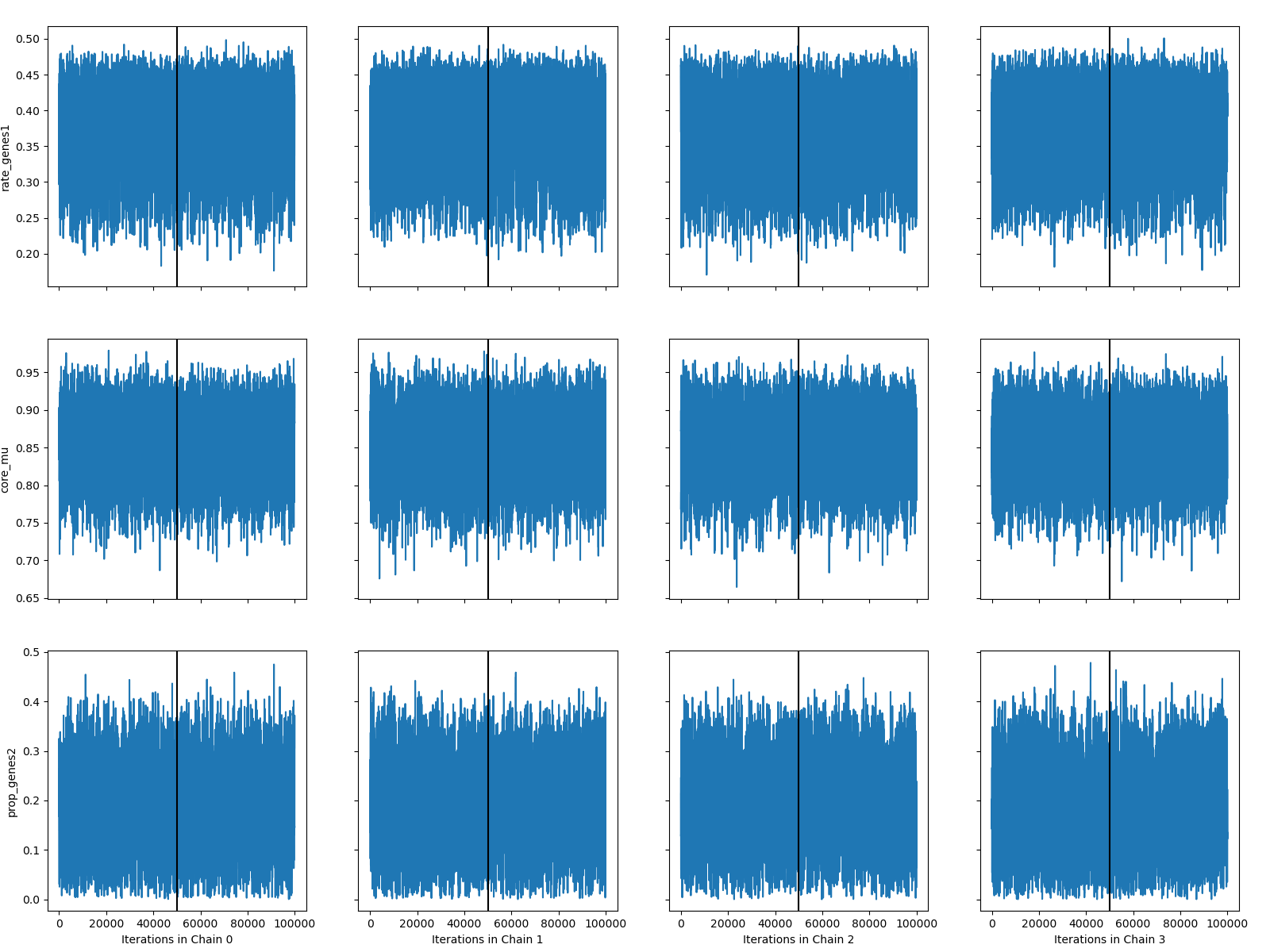

*Acinetobacter baumannii*

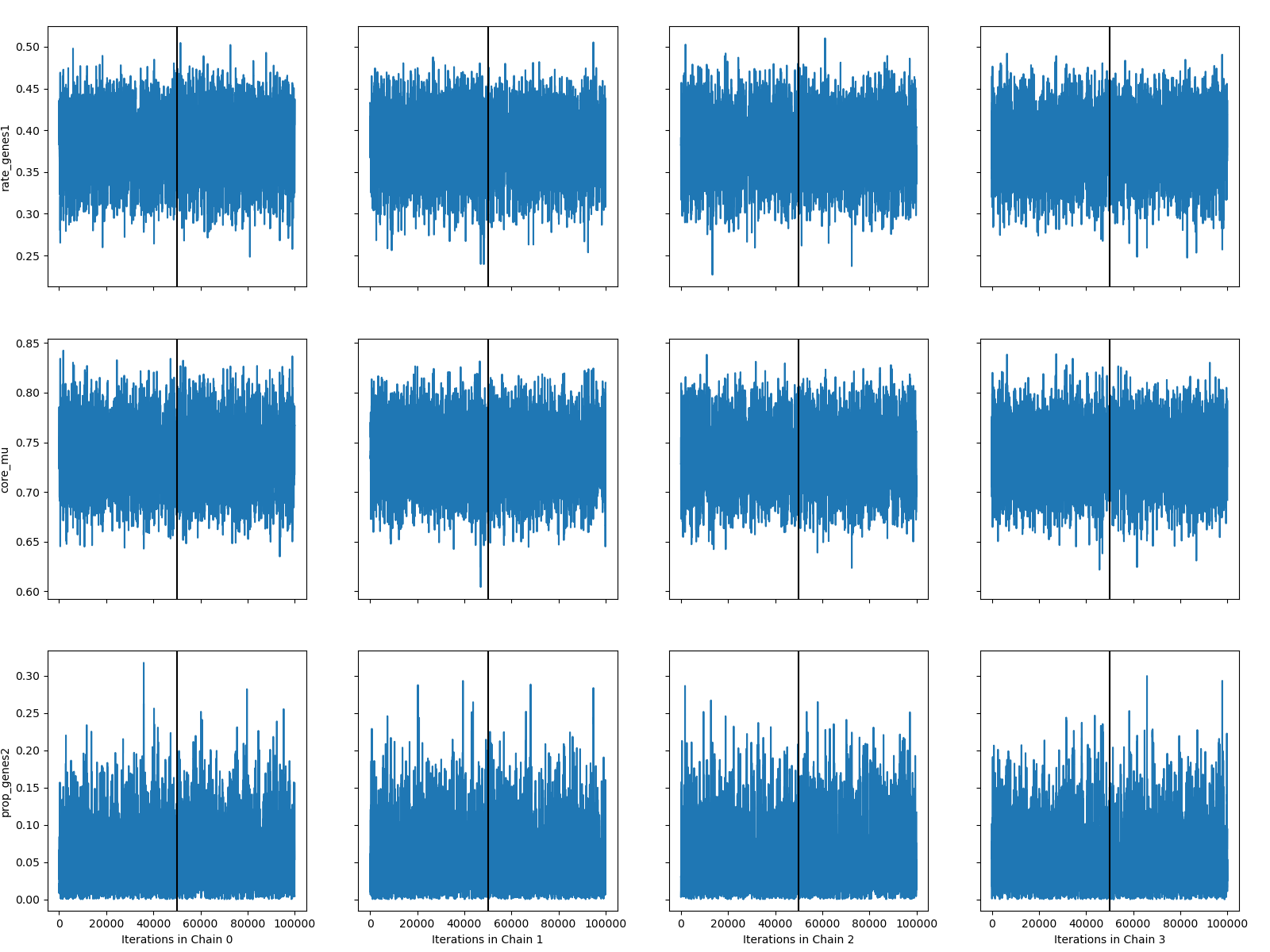

*Bordetella pertussis*

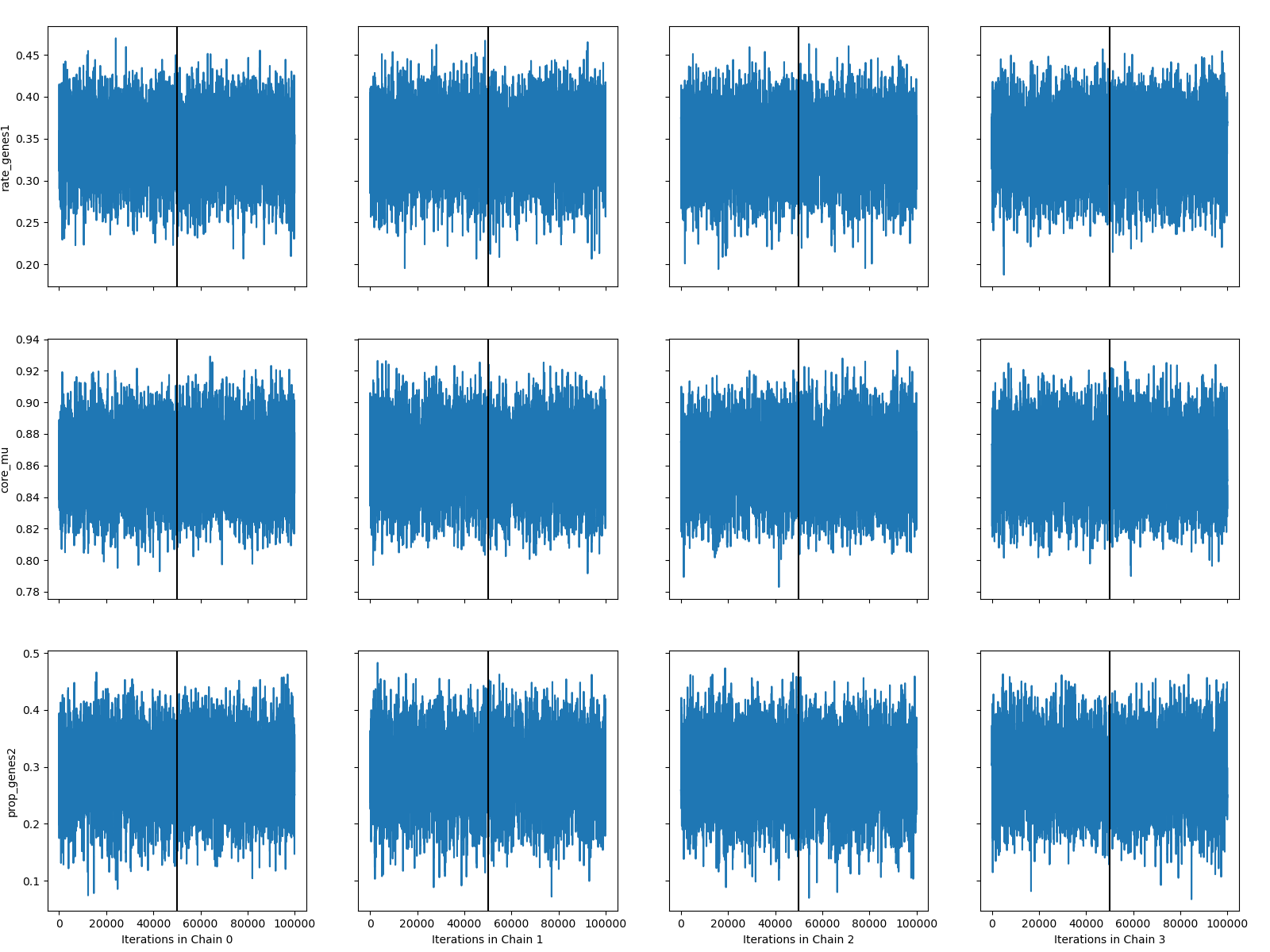

*Campylobacter jejuni*

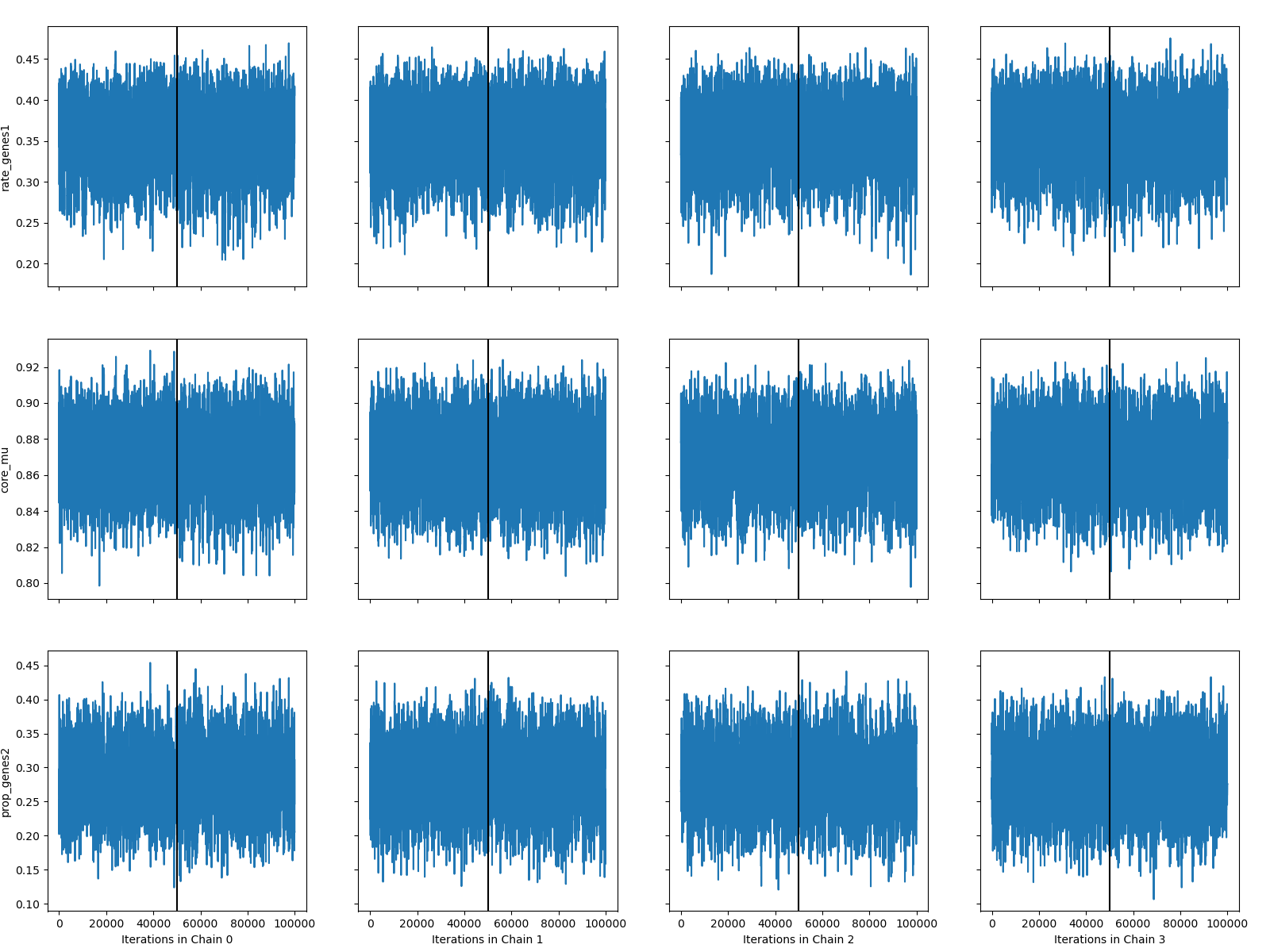

*Enterococcus faecalis*

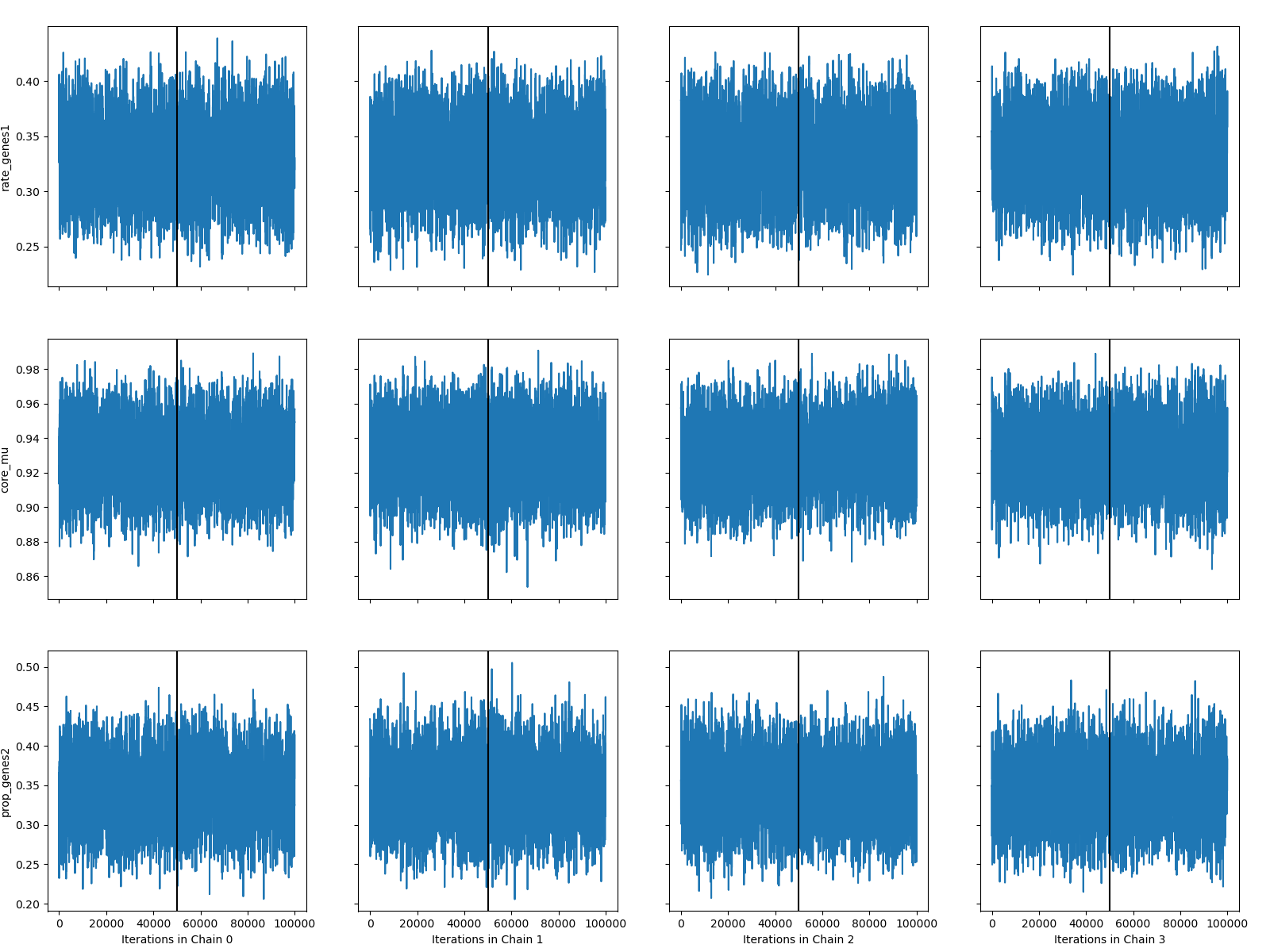

*Escherichia coli*

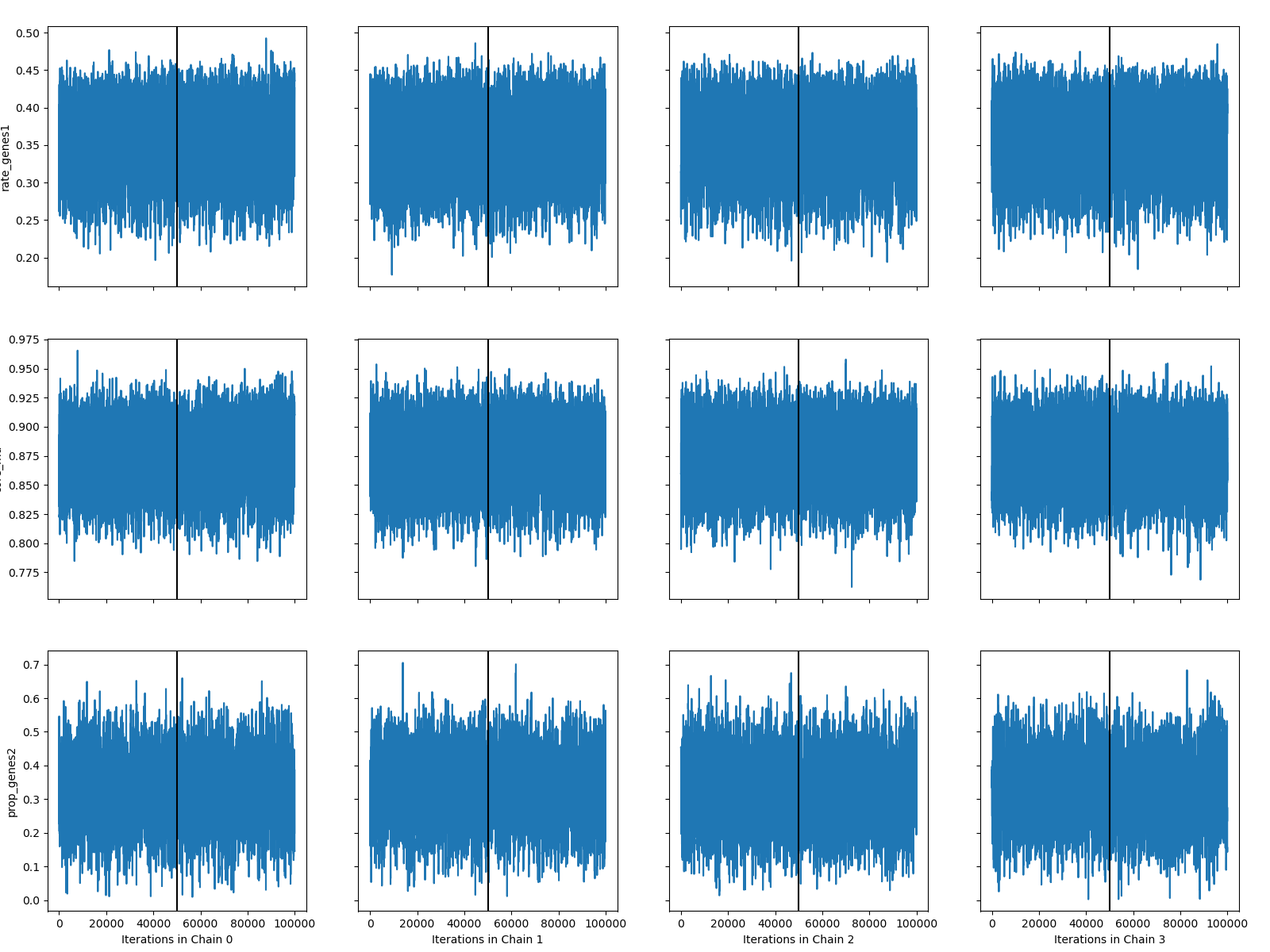

*Haemophilus influenzae*

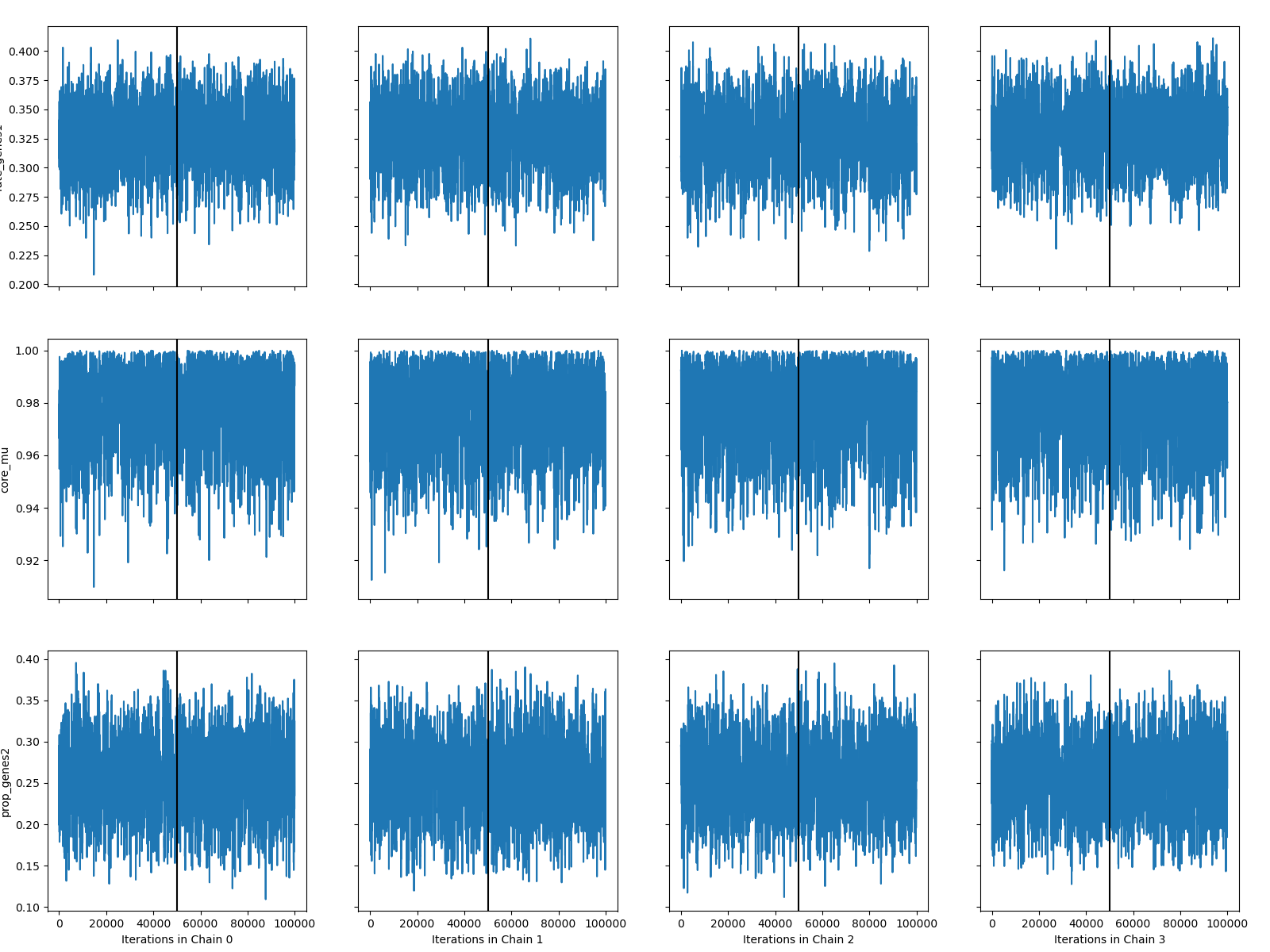

*Helicobacter pylori*

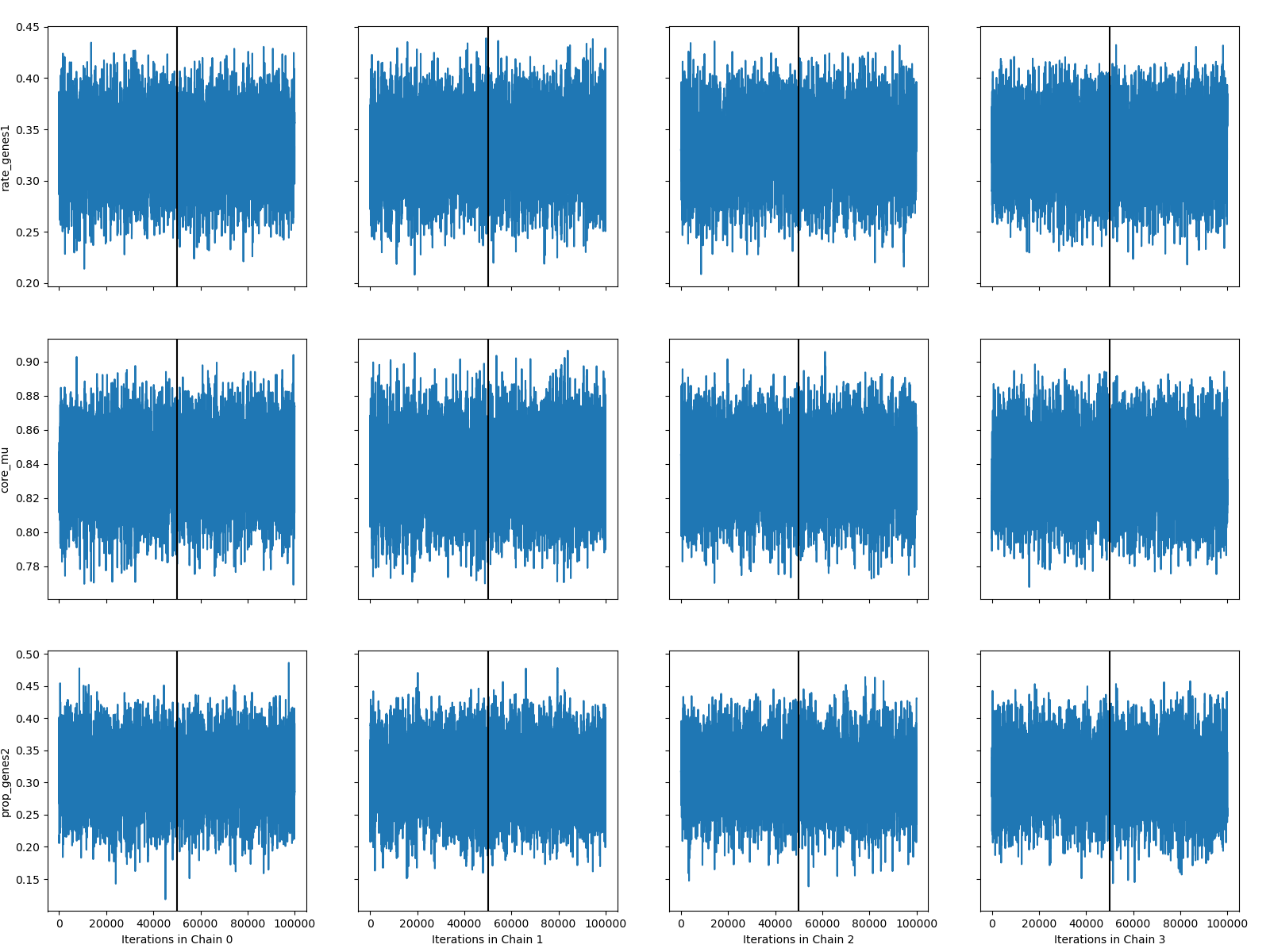

*Klebsiella pneumoniae*

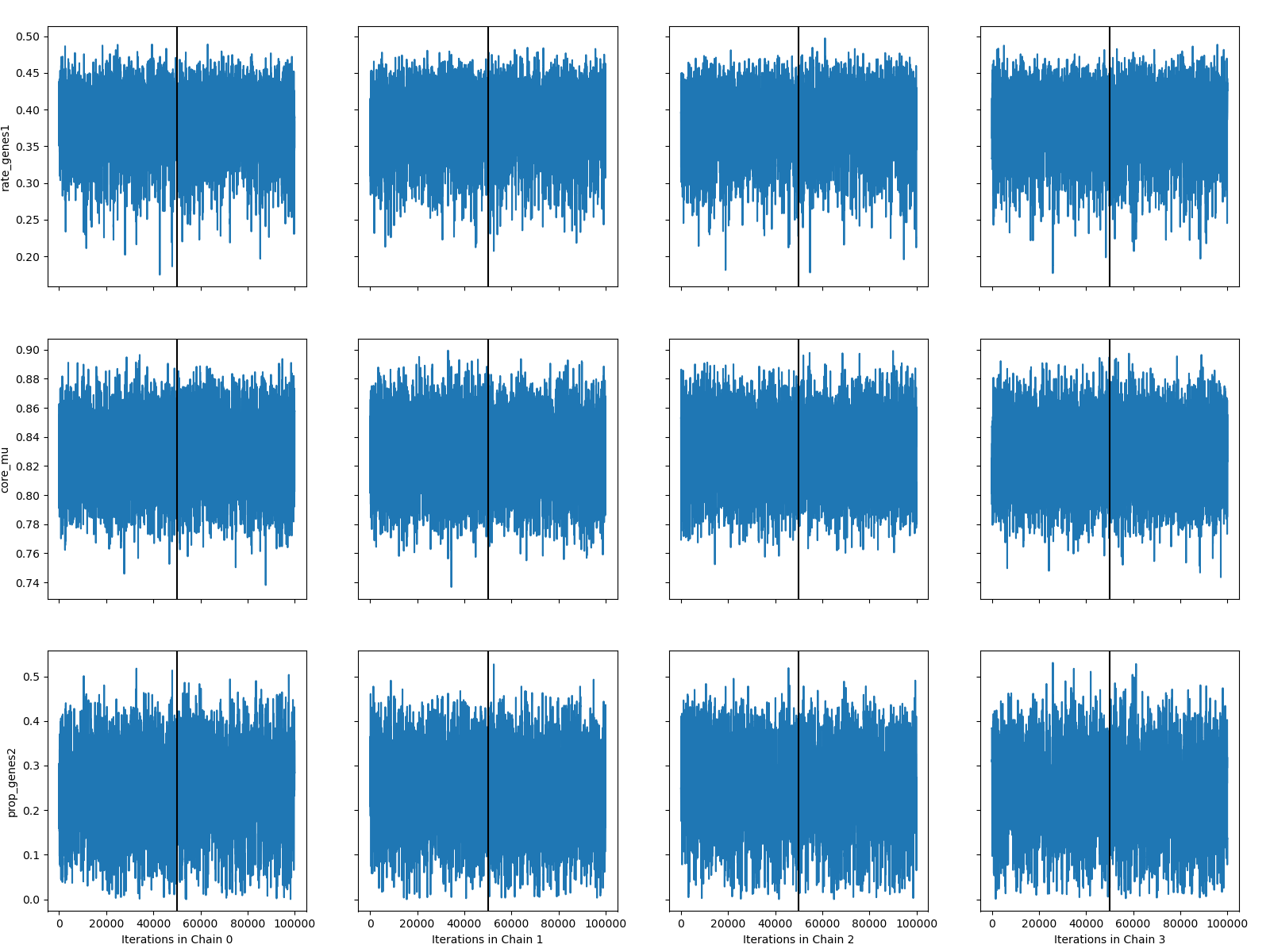

*Listeria monocytogenes*

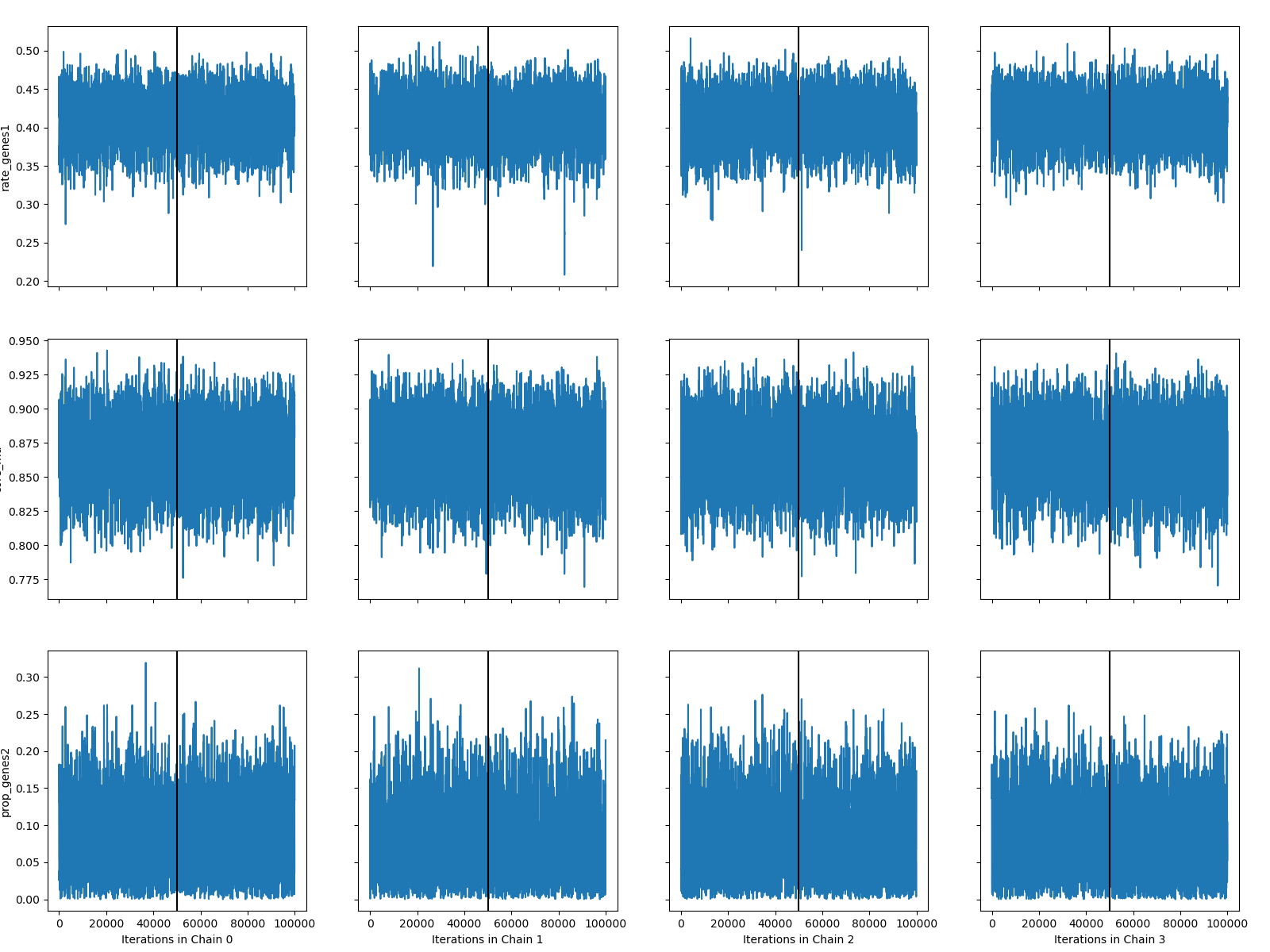

*Moraxella catarrhalis*

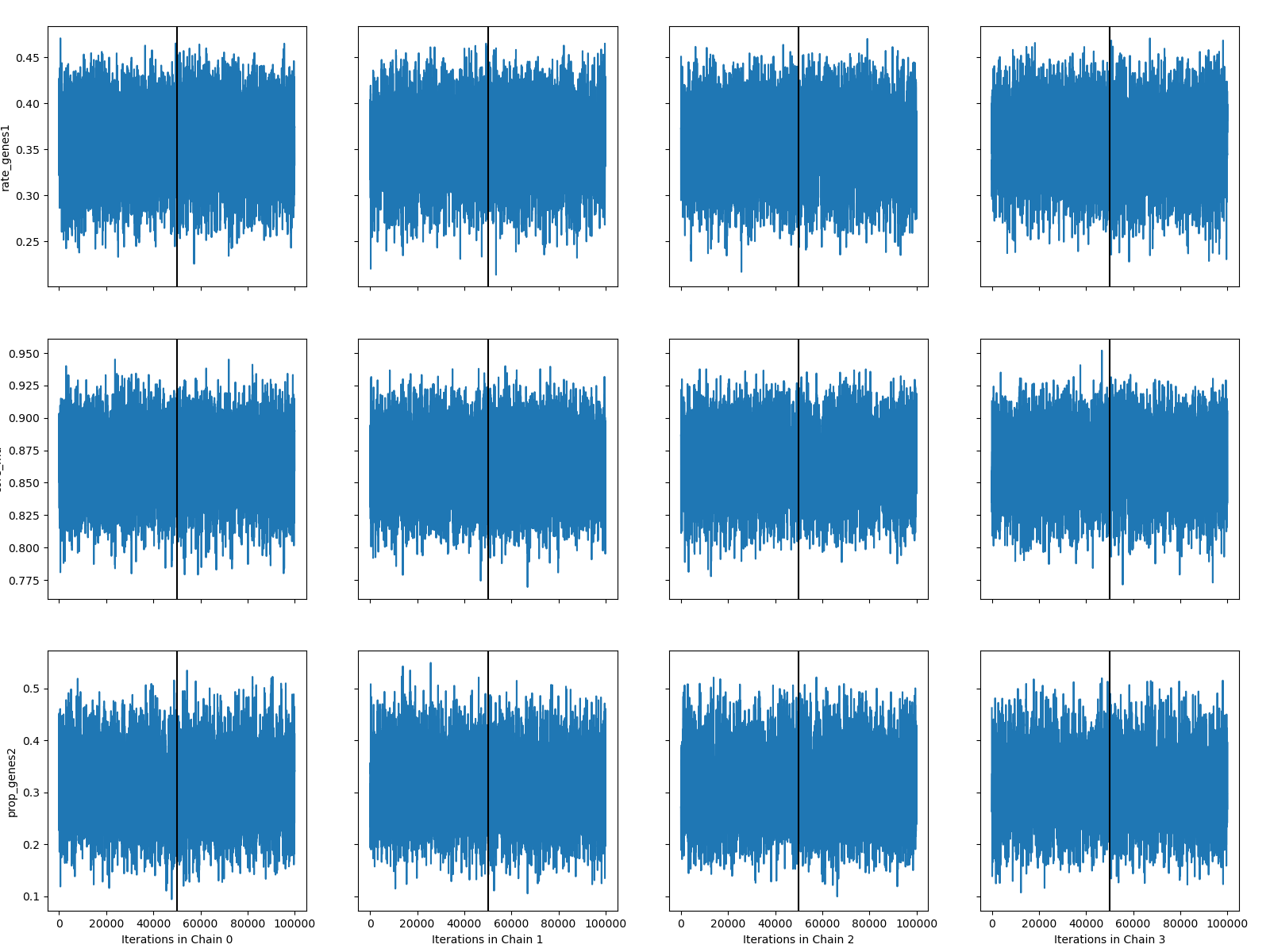

*Mycobacterium abscessus*

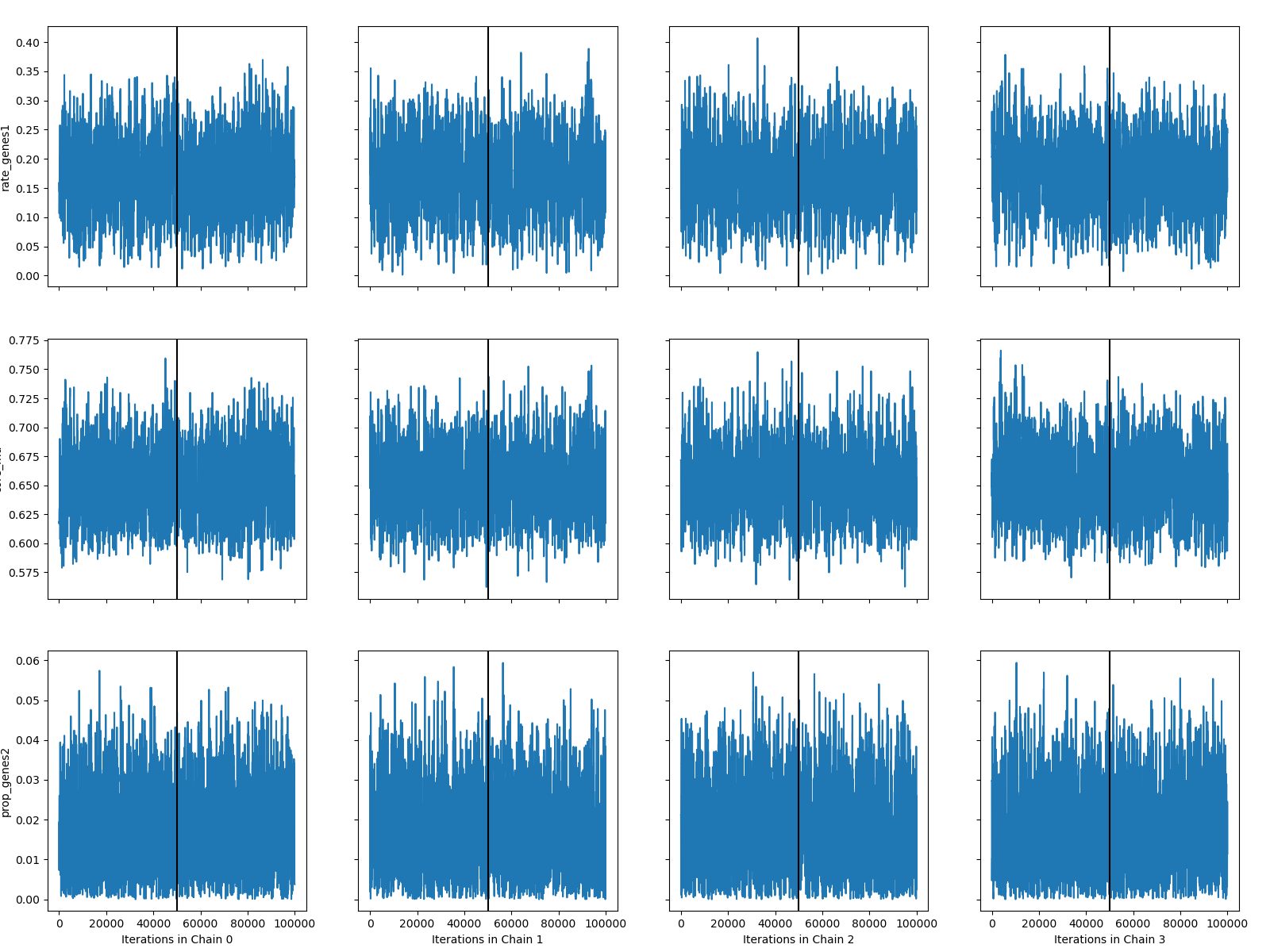

*Mycobacterium tuberculosis*

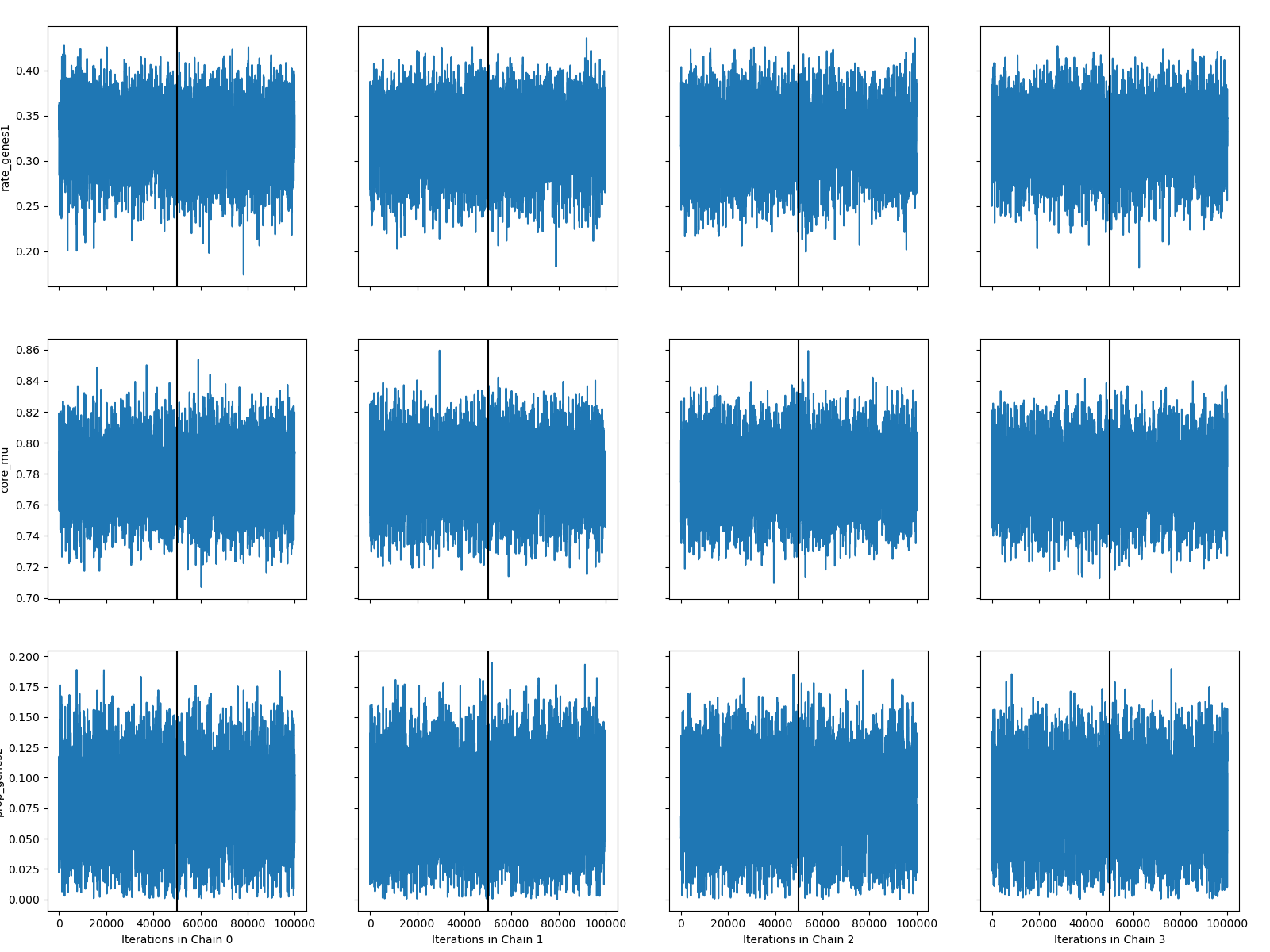
*Neisseria gonorrhoeae*

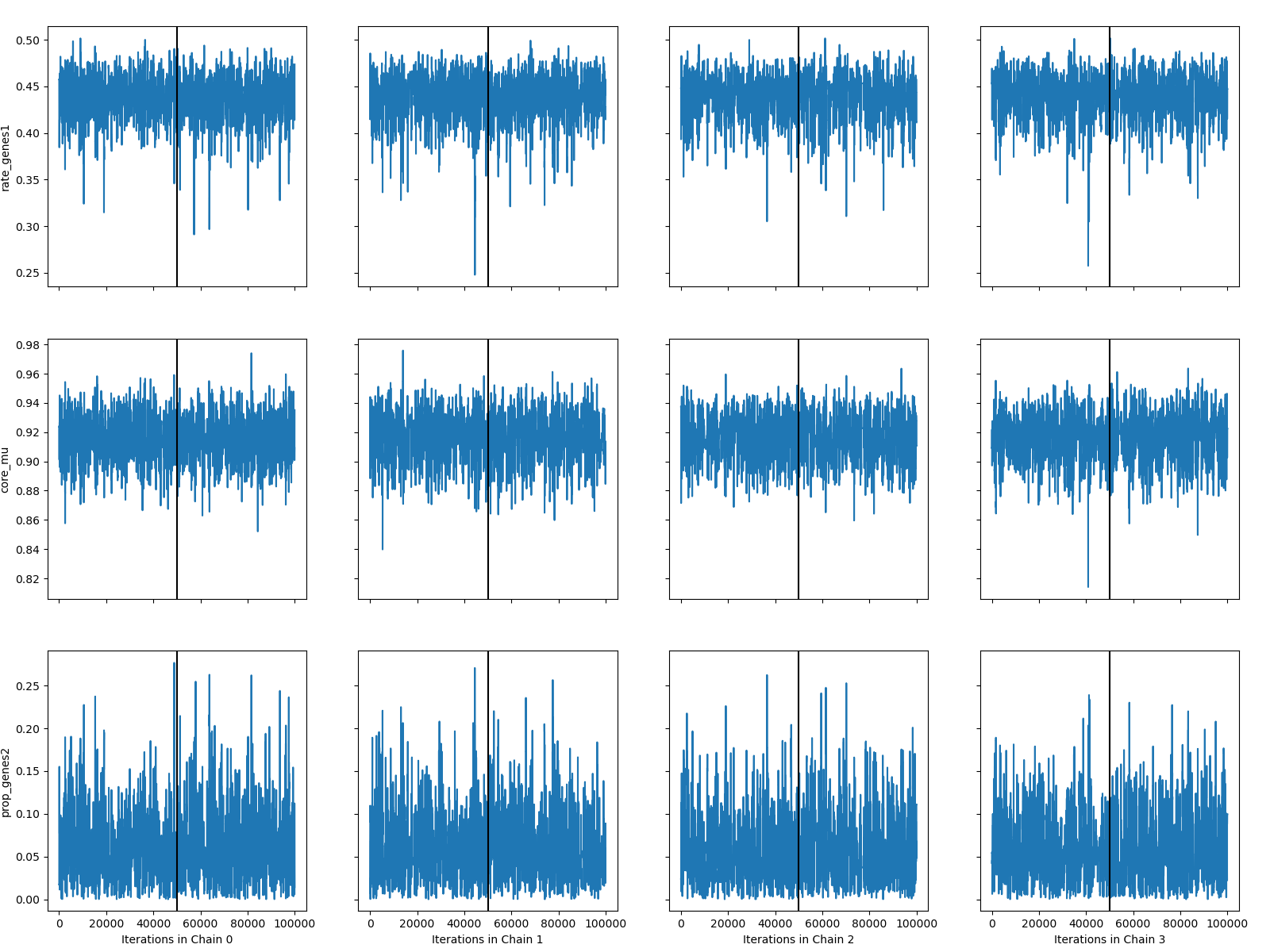

*Neisseria meningitidis*

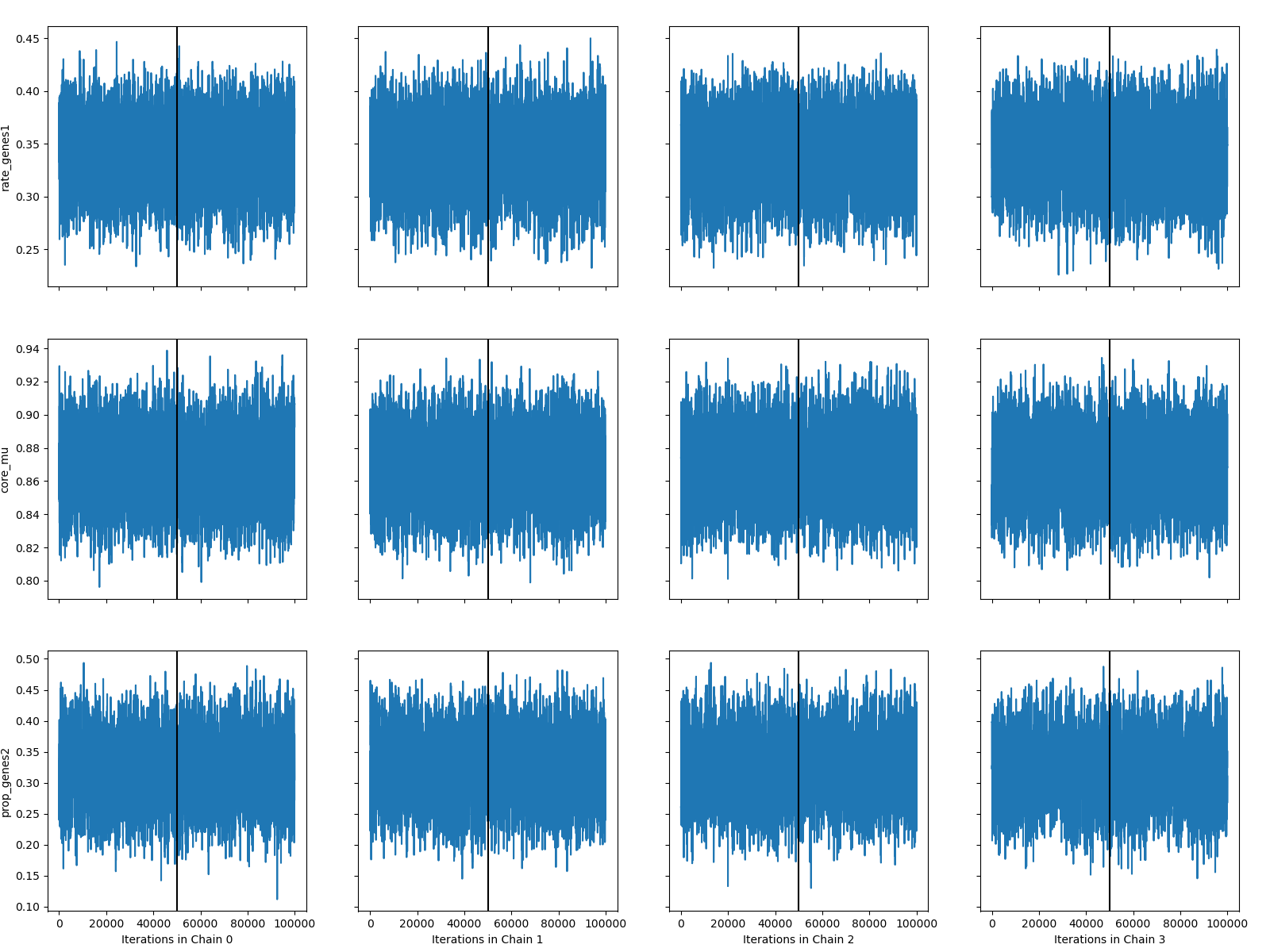

*Pseudomonas aeruginosa*

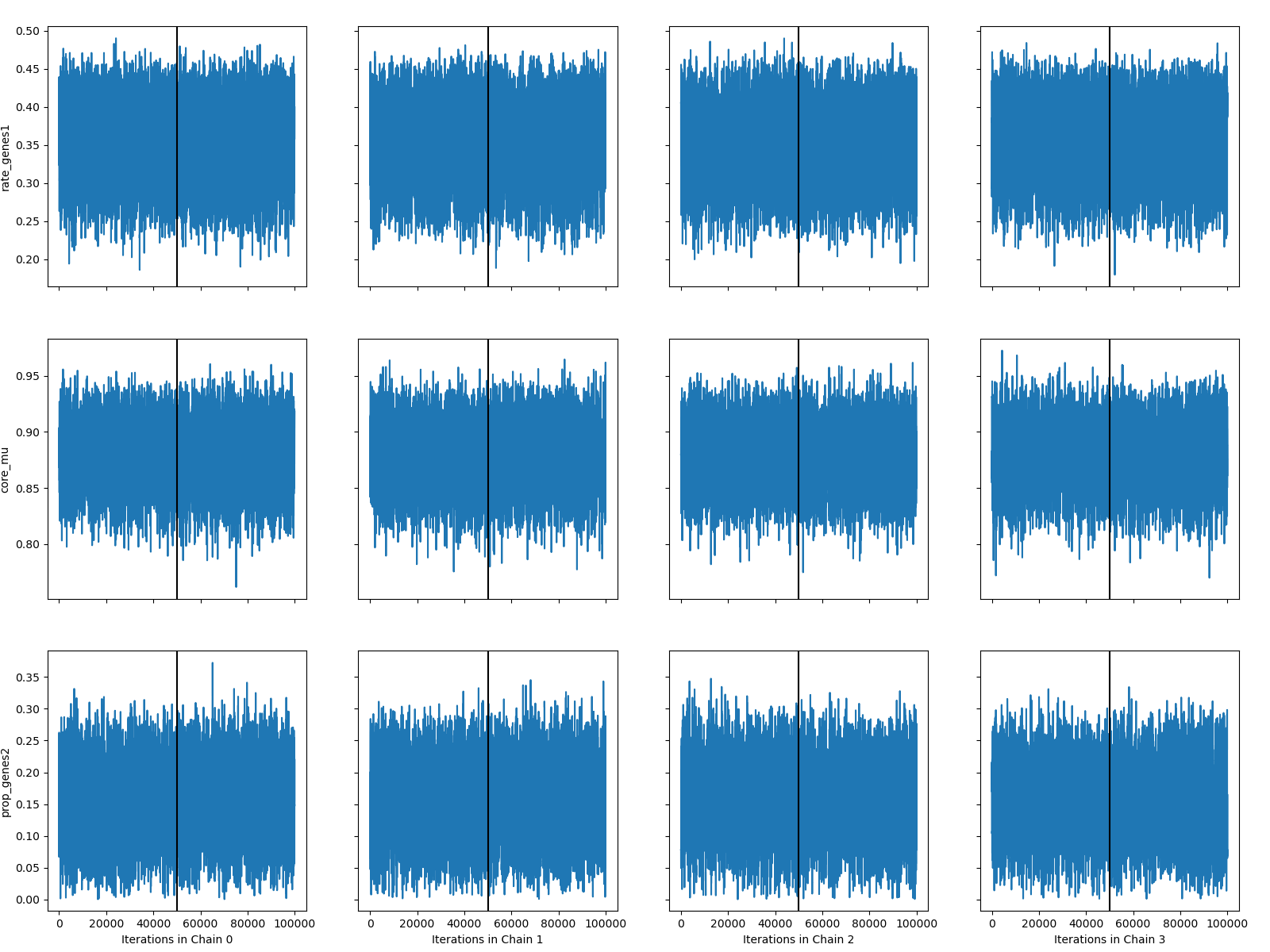

*Salmonella enterica*

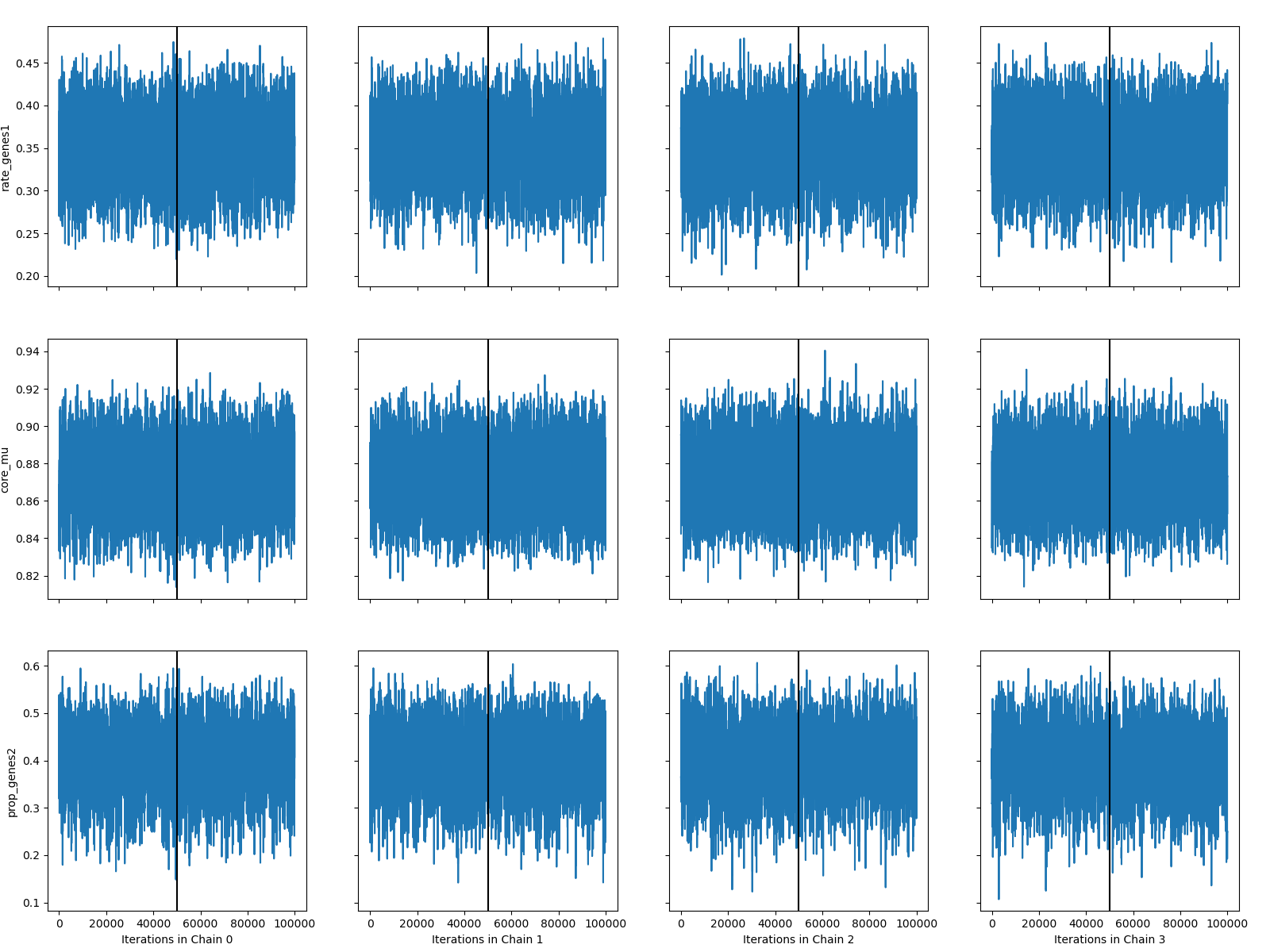

*Staphylococcus aureus*

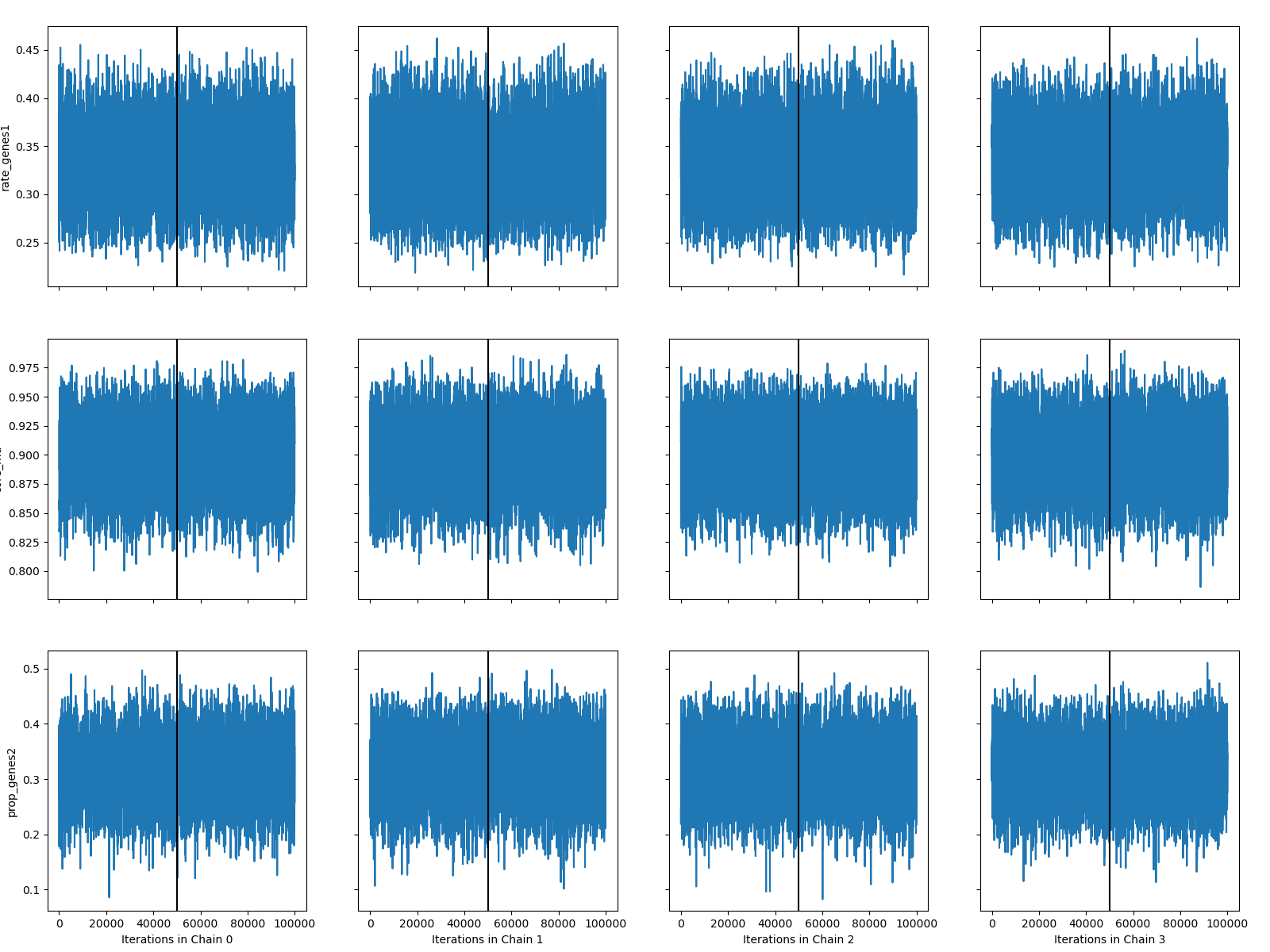

*Stenotrophomonas maltophilia*

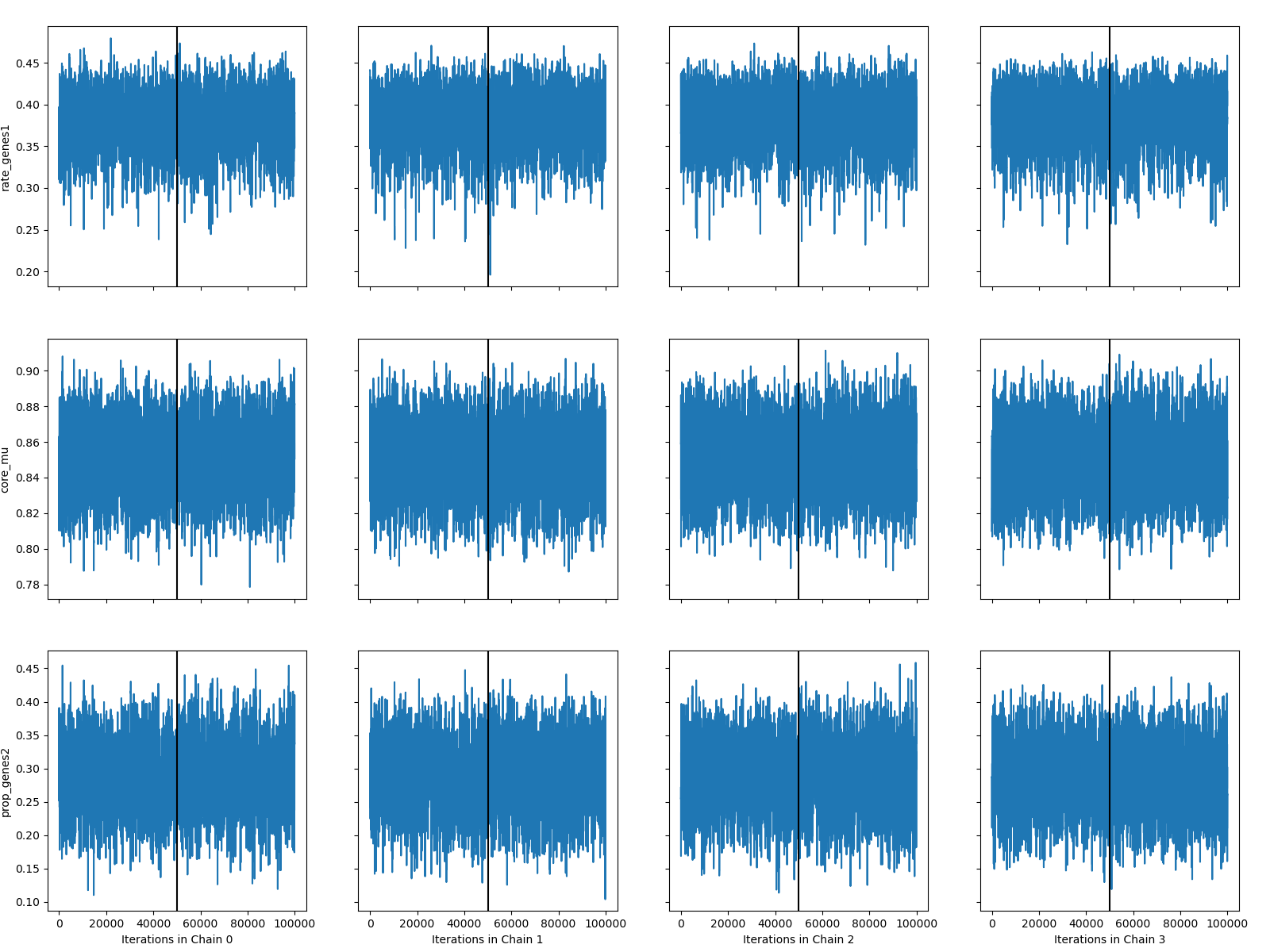

*Streptococcus agalactiae*

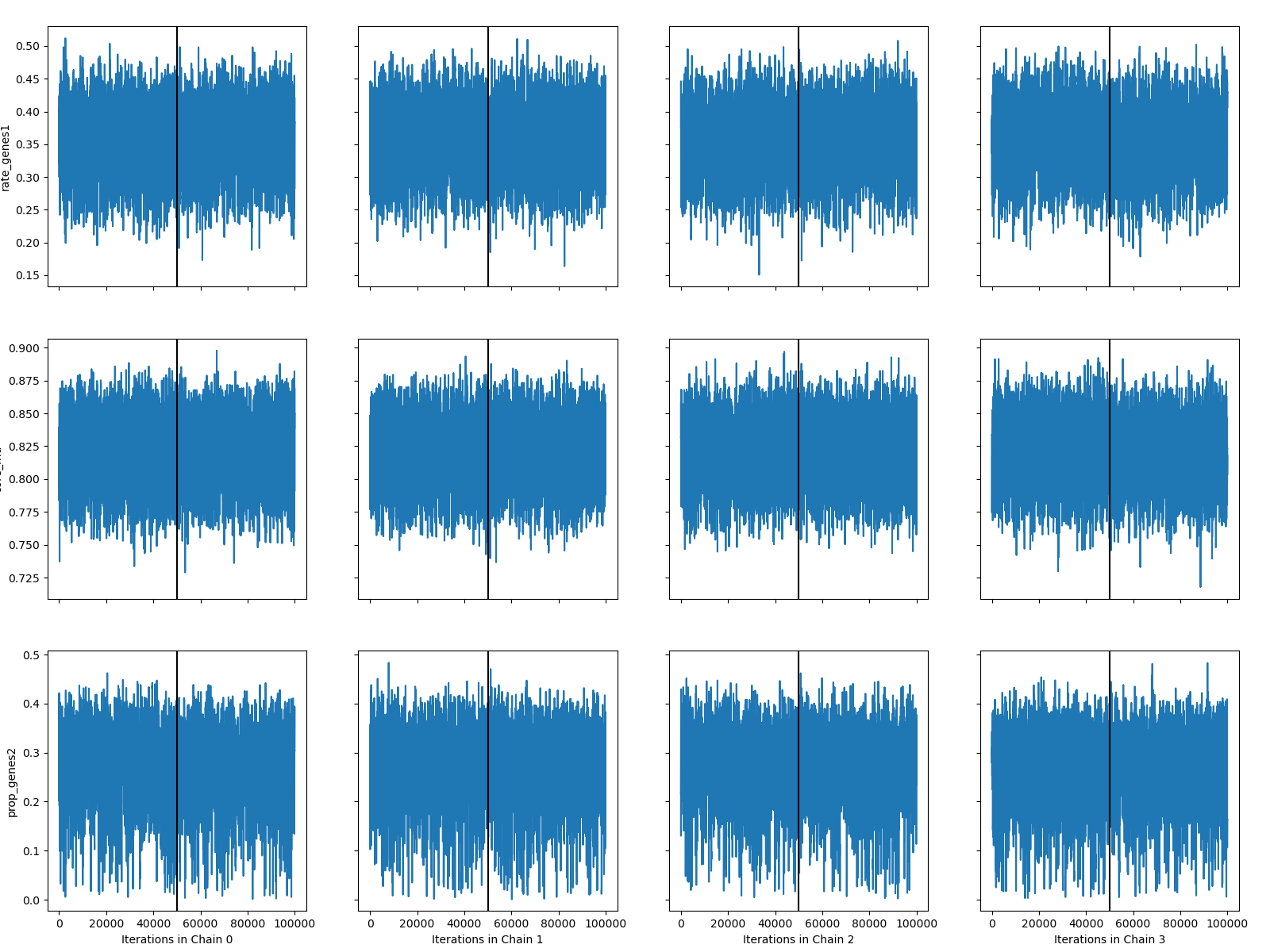

*Streptococcus dysgalactiae*

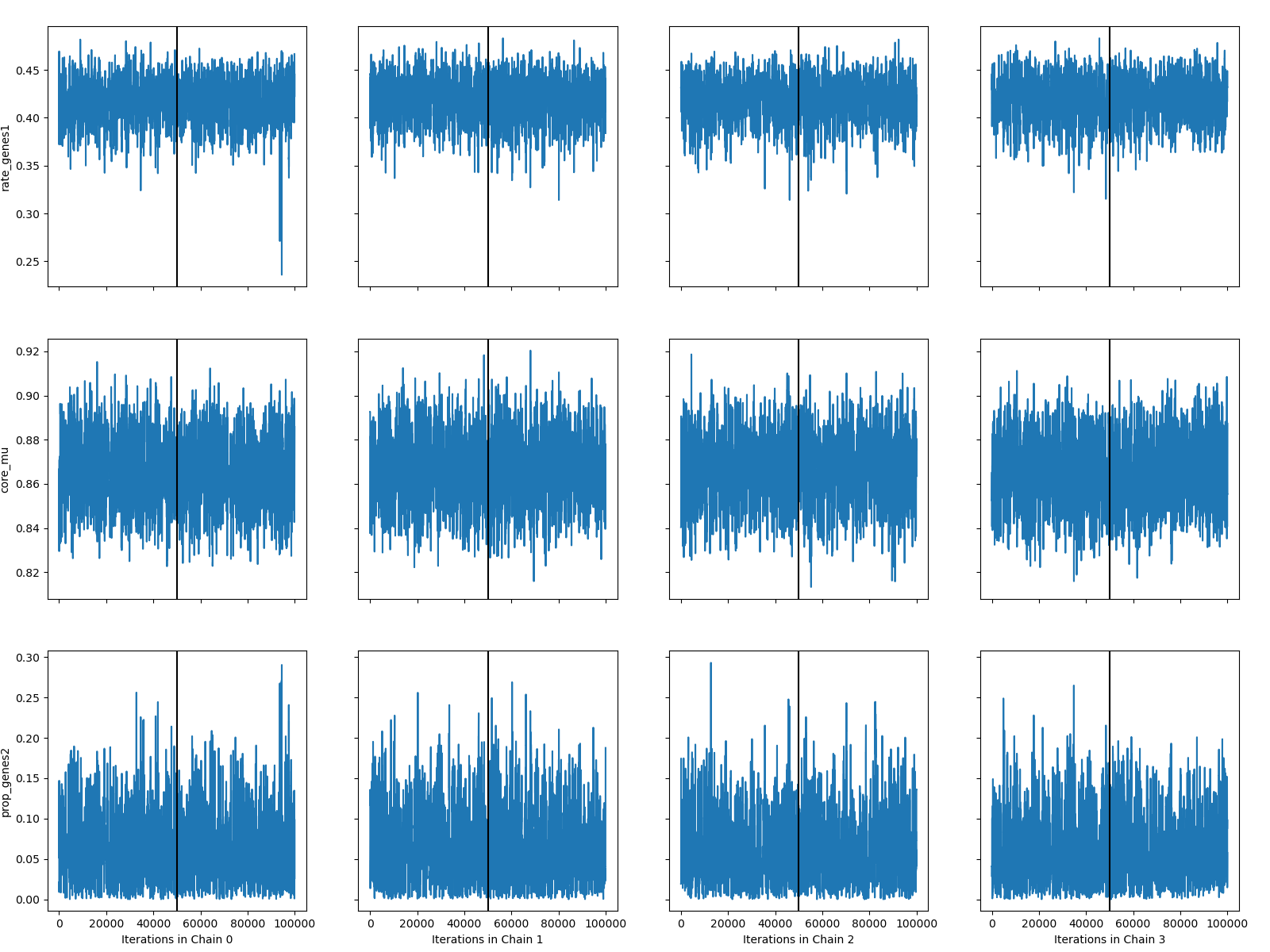

*Streptococcus pneumoniae (Global)*

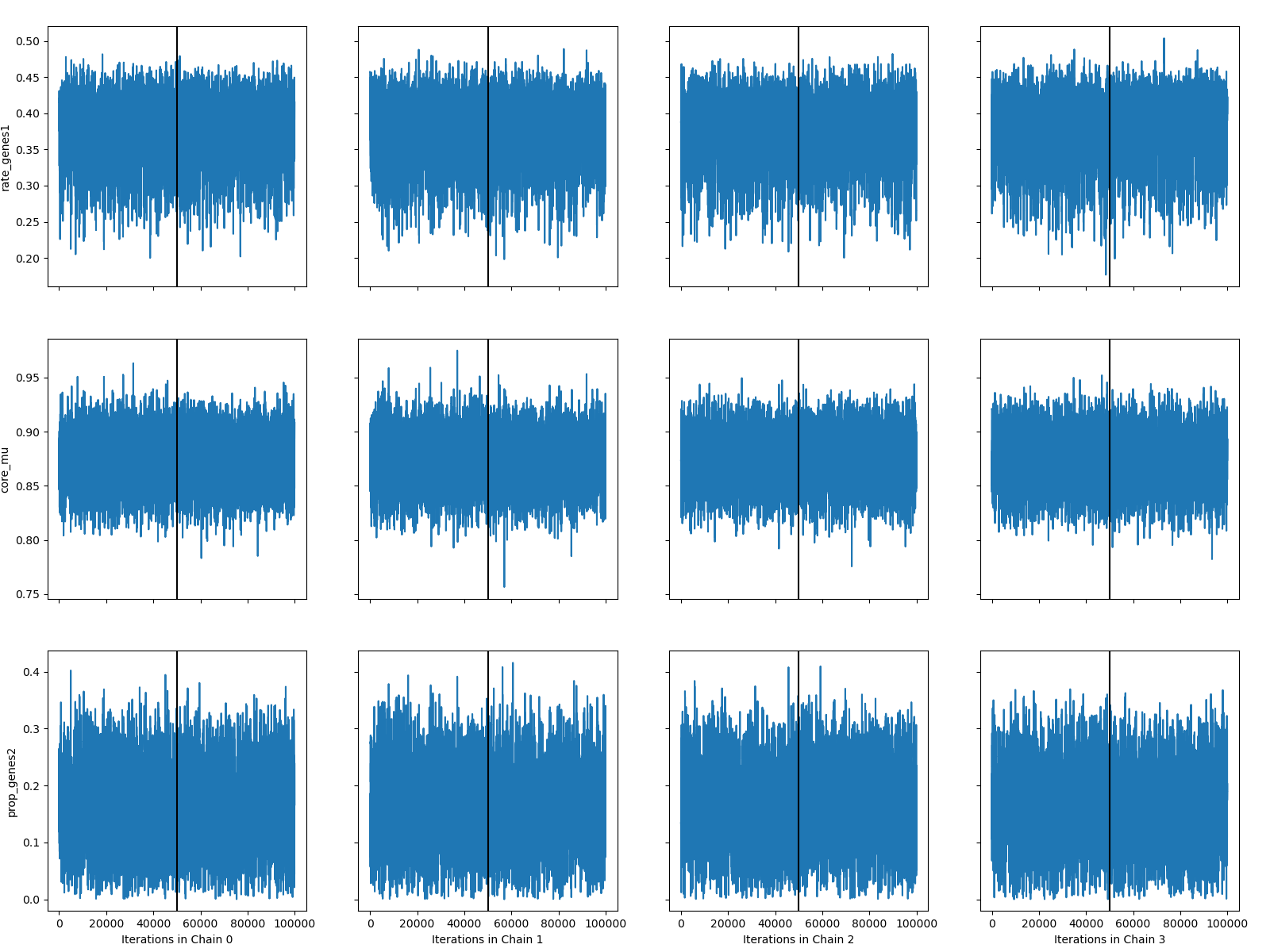

*Streptococcus pneumoniae (Local)*

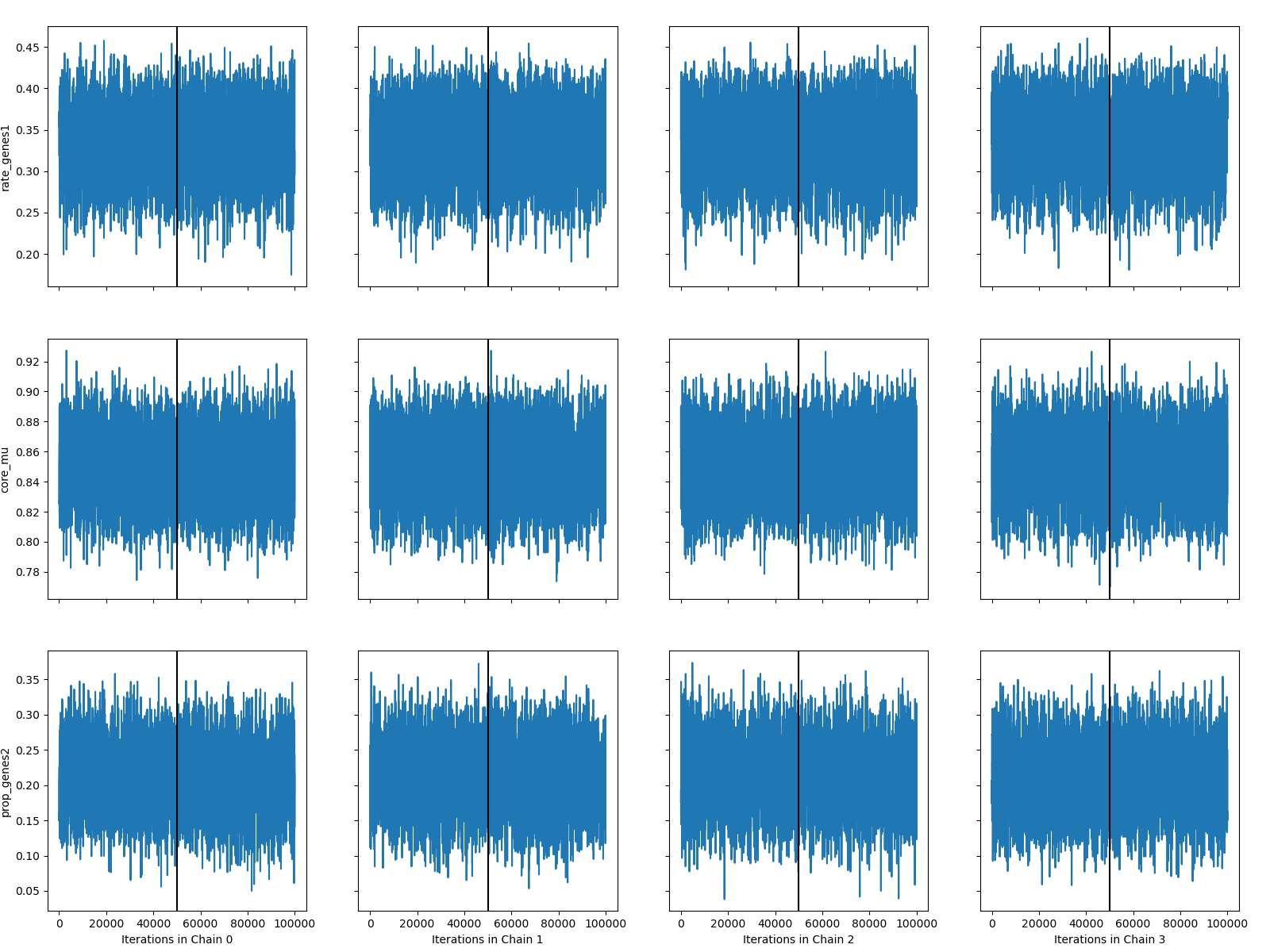

*Streptococcus pyogenes*
